# A multicellular actin star network underpins epithelial organization and connectivity

**DOI:** 10.1101/2024.07.26.605277

**Authors:** Barai Amlan, Soleilhac Matis, Xi Wang, Lin Shao-Zhen, Karnat Marc, Bazellières Elsa, Richelme Sylvie, Lecouffe Brice, Chardes Claire, Berrebi Dominique, Rümmele Frank, Théry Manuel, Rupprecht Jean-François, Delacour Delphine

**Author notes:** Authors participated equivalently (Barai Amlan, Soleilhac Matis). Authors participated equivalently (Lin Shao-Zhen, Karnat Marc).

## Abstract

Epithelial tissues serve as physical barriers against various external pressures yet remarkably maintain structural stability. Various cellular apparatus including bicellular junction and actomyosin network contribute to the epithelial integrity, packing and remodelling. Although their role in morphogenetic and mechanical processes have been extensively studied during embryogenesis and disease development, their synergistic effects in maintaining tissue organization and connection remain poorly understood. In this study, we discovered a tissue-scale actomyosin network connected through bicellular junctions and manifested in the villi of adult murine intestinal tissue. Later we reproduced such supracellular structure in the differentiated compartment of *ex vivo* intestinal epithelium model. The self-organized actomyosin networks comprised individual actin nodes in each hexagonal cell at the epithelial base with six radial actin branches, presenting an ‘actin star’ unit. The repeated units were connected through the bicellular junctions, forming a large, multicellular array covering the differentiated domains. Functionally, actin stars contribute to epithelial morphological stability by maintaining cell hexagonality and packing, thereby preserving the solid-like order of the epithelium. Laser ablation experiments validate a modified vertex theoretical model that connects the emergence of such solid-like order to the onset of tension along the actin star branches. Actin stars also acted as locks at the basal side minimizing protrusive activity in the epithelial layer, hindering cell migration and disorganization of the epithelial tissue. Altogether, the supracellular actin star network constitutes a basal biomechanical apparatus coordinating epithelial tissue stability and organization.

## INTRODUCTION

The actomyosin network ensures numerous morphogenetic processes through the generation and transmission of tension forces ^1,2,3^, which are primarily facilitated by the presence and activation of the molecular motor myosin-II (non-muscle myosin-II, NM-II) along actin filaments ^4^. The pivotal functions of the contractile apparatus are particularly emphasized in epithelial tissues, which form resilient cellular assemblies lining the body surfaces. Monolayered epithelial behave as crucial interfaces between the internal milieu and the external environment, orchestrating essential physiological processes such as protection, secretion, and absorption. Thus, the preservation of epithelial tissue integrity stands as a paramount concern. Any disruptions in this integrity can yield profound consequences, including compromised organ formation and functionality, and potential tumorigenesis ^5,6^.

Epithelial monolayers possess distinct characteristics, marked by the polarization of densely packed cells into cohesive sheets. Hence, hexagonal packing represents an efficient way of covering a surface and distributing force equally among epithelial cells ^7^. Moreover, cell polarization gives rise to discrete apical and basolateral domains, each endowed with unique molecular compositions and functions that dictate directional epithelial functions ^8,6^. To maintain epithelial monolayer cohesion and coordination, cells establish robust intercellular junctions, notably adherens junctions mediated by cadherins and linked to actomyosin network via catenins, particularly enriched in the actin belt on apical side ^9,10,11^. It is now clear that the apical coupling of E-cadherin, catenin and actomyosin is a powerful component for tissue mechanics, as it serves as a mechanosensor unit by responding to applied forces ^11,12,13^. Therefore, the integration of mechanical information at the tissue level, achieved through adaptation of the actomyosin cytoskeleton or junctions themselves, is a major component of epithelial morphogenesis ^11,13^. For instance, the apical-medial actomyosin drives apical constriction during mesoderm invagination in *Drosophila* embryos for apical area deformation. Apical constriction allows apical area deformation and ultimately tissue folding through mechanical forces applied on adherens junctions ^14,15,16,17,18^. In addition, anisotropic changes of tension along the cell surface control cell deformation that result in cell intercalation. During this process, cells remodel their junctions with neighbouring cells to permit cellular rearrangement during tissue elongation ^19^. Altogether, these morphogenetic events directly impact the tissue physical state or tissue fluidity. Tissues in a solid-like state are barely remodelled and able to resist mechanical stress to maintain their architecture, while tissues in a fluid-like state are more prompt to remodelling due to exerted forces ^20^.

Here, we used the mammalian intestinal tissue to address mechanisms that control epithelial stability. We revealed the development of a multicellular star-shaped actomyosin lattice in the basal domain of differentiated epithelial cells, that behaves as a new mechanical unit at play for morphological and functional stability in the intestinal tissue.

## RESULTS

### Differentiated intestinal epithelial cells develop basal actin star-like structures

The mouse intestinal tissue is a suitable working model for investigating the collective organization of an adult mammalian epithelium. Its functional unit, known as the crypt-villus axis, exhibits distinct compartments: a proliferative domain represented by the crypt (in blue, Figure 1A) and a differentiated domain comprising the villus (in yellow, Figure 1A). Examination of adult mouse villi offered a direct glimpse into the *in vivo* organization of this columnar epithelium (Figure 1B). Through actin labelling in whole-mount tissue, we achieved a clear visualization of the brush border at the apical pole and the basolateral cell contacts (Figure 1B a,d). Notably, our observations revealed a regular actin meshwork located beneath the nuclei on the basal side of epithelial cells along the villus (Figure 1B b-d, Video S1). This meshwork comprised a dense actin core centrally located in each cell basal side, with discrete branches of actin bundles seemingly linking to cell-cell contacts (Figure 1B b-c). These ‘star-shaped’ actin-based structures were commonly observed in most epithelial cells along the villus body, except for those at the villus base (where cells complete terminal differentiation) and the villus tip (where cells undergo preparation for death and extrusion) (Supplementary Figure 1A-B) ^21^. Additionally, such ‘actin stars’ (AcSs) were conspicuously absent in crypt cells (Supplementary Figure 1C-E, Video S2), underscoring their specificity to the intestinal differentiated domain.

**Figure 1:**
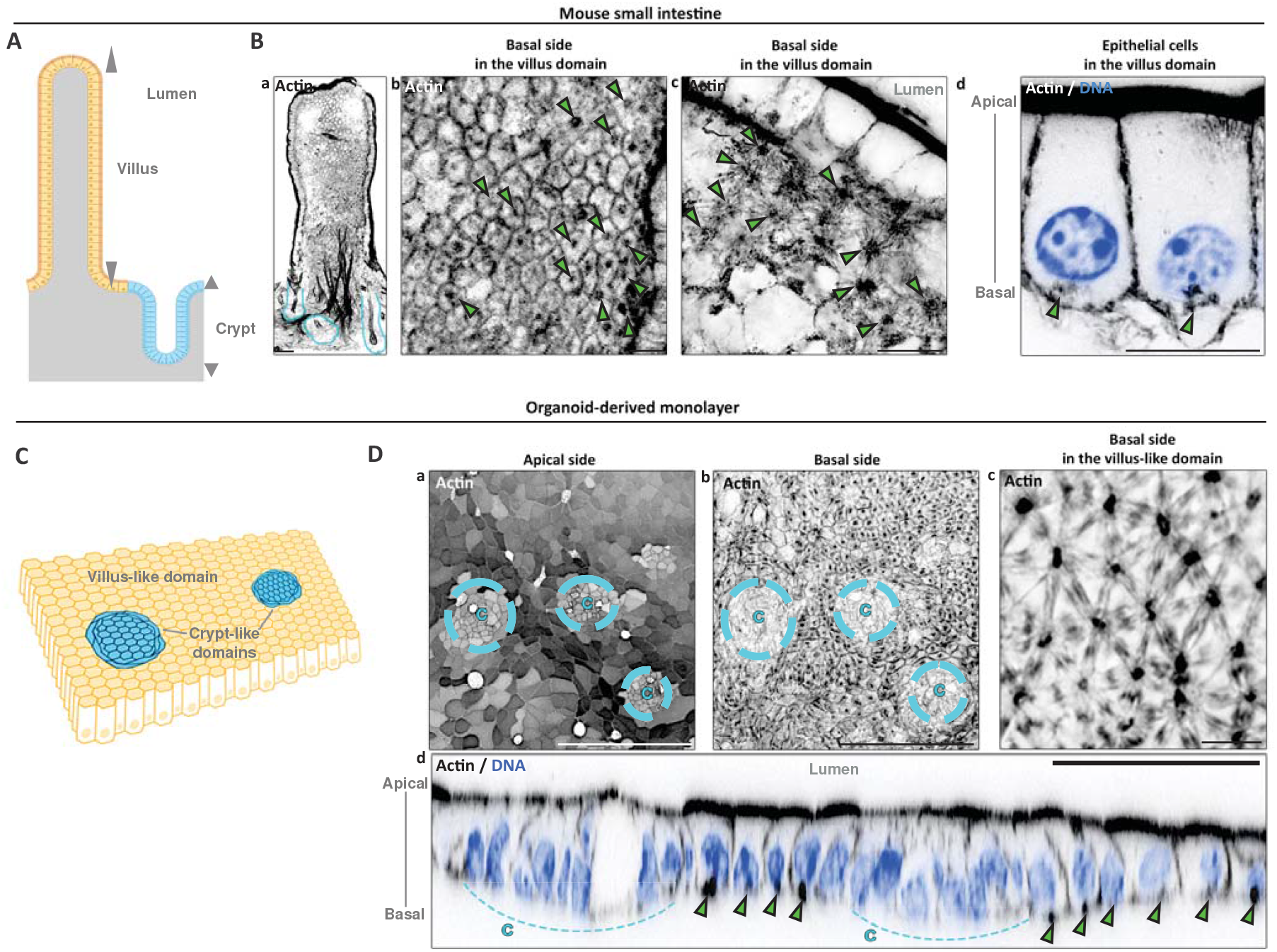
Differentiated intestinal epithelial cells display star-shaped actin cytoskeleton. **(A)** Scheme showing the functional organization of the mouse small intestinal tissue where the proliferative crypt domains are shown in blue and the differentiated villus compartment in yellow. **(Ba-d)** Confocal analysis of actin distribution in the villus domain. Nuclei are stained in blue in (d). Green arrowheads point to actin stars (AcSs). Scale bars, (a) 50 μm, (b-d) 10 μm. **(C)** Scheme showing the self-organization of intestinal organoid-derived monolayer with the proliferative crypt-like domains in blue and the differentiated villus-like compartment in yellow. **(Da-d)** Confocal analysis of actin distribution in the apical or basal side of an organoid-derived monolayer. xz view is presented in (d). Crypt-like domains are delimited in blue. Green arrowheads point to actin stars. C, crypt-like domain. Scale bars, (a-b) 100 μm, (c) 10 μm, (d) 50 μm.

To delve deeper into the characterization of AcS structures and enhance their spatial resolution, we embarked on reproducing such network within a previously established 2D intestinal organoid culture ^22^. In brief, organoid-derived monolayers were grown on soft substrates made of cross-linked (CL) Matrigel^®^ matrix, on which they mimicked the patterning of intestinal tissue *in vitro* (Figure 1C). These include self-organizing crypt-like domains enriched with proliferative cells (i.e. EdU-positive cells, Supplementary Figure 2A), encircled by large villus-like domains with differentiated cells (i.e. cytokeratin-20- and ezrin-positive cells, Supplementary Figure 2B,D) ^23,24,22^. Within the differentiated domain, we noticed a discernible brush border at the apical surface of enterocytes (Figure 1D a, Supplementary Figure 2D) and a similar star-shaped actin structure was present at the basal surface of nearly all individual cells (Figure 1D b-c, Supplementary Figure 2C-D), closely mirroring our *in vivo* findings (Figure 1Bb-d). Furthermore, MUC2-positive goblet cells also exhibited AcSs, with actin branches extending continuously into the AcS of neighbouring enterocytes (Supplementary Figure 2E-F). Remarkably, AcSs were absent in the crypt-like domain, like in the *in vivo* tissue (Figure 1D b,d, Supplementary Figure 2D). Thus, the development of basal AcSs network in the 2D differentiated domains emulated those in the intestinal villi. The development of basal AcSs emerged as a significant hallmark of the differentiated domain in intestinal epithelium, alongside the apical brush border.

### The assembly of basal actin stars creates an epithelial supracellular cytoskeletal network

Further structural analysis using high-resolution microscopy revealed that each epithelial cell of the differentiated domain exhibited a single AcS unit, featuring an actin node positioned at the centre of mass of the basal surface (Figure 2A,C) and accompanied by ∼ 6 actin branches extending from it (Figure 2A-B,D). AcS branches measured approximately 5 μm in length (Figure 2E) and exhibited a regular radial organization, with an angle of *α* = 59±15.75° between actin branches (Figure 2F). Moreover, the actin branch was oriented perpendicular (*β* = 89±13.13°) to an adjacent bicellular contact (Figure 2A-B,F), and appeared to maintain directional continuity with a branch of the neighbouring cell’s AcS (Figure 2B, insert). Consequently, the collective disposition of AcSs within the differentiated epithelial layer delineates a triangular network (Figure 2A-B) with AcS playing the role of a Delaunay tessellation of the Voronoi-like cell membrane network.

**Figure 2:**
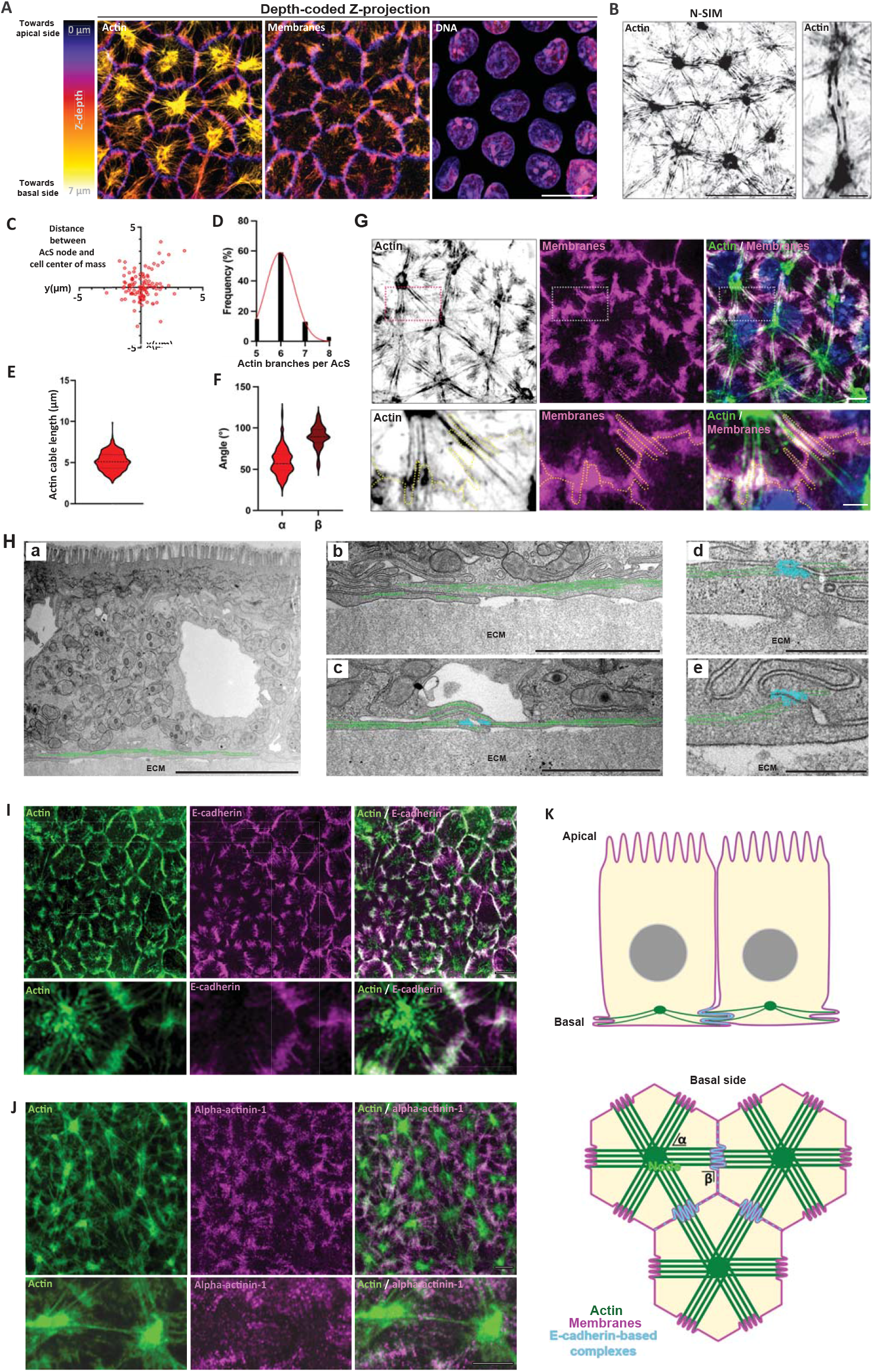
Basal actin star assembly creates a multicellular connecting network. **(A)** Airyscan microscopy and depth-coded z-projection of actin, membranes and DNA localization in the basal side of organoid-derived monolayers. Scale bar, 10 μm. **(B)** N-SIM microscopy analysis of basal actin distribution in organoid-derived monolayers. Scale bar, 15 μm, insert 2 μm. **(C)** Statistical analysis of the distance between the AcS node and the cell center of mass in organoid-derived monolayers. Mean distance = 1.48±1.46 μm (mean±S.D.). N = 3 experiments, n = 113 cells. **(D)** Statistical analysis of the number of actin branches per AcS. Mean number of actin branches per star = 6.04±0.67 (mean±S.D.). N = 3 experiments, n = 90 cells. **(E)** Statistical analysis of the actin cable length in AcSs. Mean length = 5.16±1.13 (mean±S.D.). N = 4 experiments, n = 331 cells. **(F)** Statistical analysis of the angle formed between actin cables (*α*) or between actin cables and plasma membranes (*β*). Mean *α* = 59±15.75° (mean±S.D.), mean (*β*) = 89.3±13.13°. N = 4 experiments, n (*α*) = 162 cells, n (*β*) = 55. **(G)** Airyscan analysis of actin (green), membranes (magenta) and DNA (blue) in organoid-derived monolayers. Scale bar upper row 5 μm, low row 2 μm. **(Ha-e)** Transmission electron microscopy analysis of the basal domain of differentiated cells in organoid-derived monolayers. Actin cables are pseudo-colored in green, adherens junction-like adhesions in light blue. ECM, extracellular matrix. Non-pseudo-colored images are shown in Supplementary Figure 2 C. Scale bars, (a) 5 μm, (b-c) 2 μm, (d-e) 0.5 μm. **(I-J)** Confocal analysis of E-cadherin and α-actinin-1 at the basal side of organoid-derived monolayers. Lower panel shows zoomed-in representative image of cell-cell junctions. Scale bar, 5 μm. **(K)** Scheme describing the architecture of AcS network (green) and cell membranes (magenta) in the basal side of organoid-derived monolayers. E-cadherin-based complexes observed in (H-I) are depicted in blue.

We investigated the potential connection between AcSs and the microtubule network. To assess the localization of the microtubule-organizing centre (MTOC), we generated centrin-1-GFP organoids (Supplementary Figure 3A). While AcSs developed at the basal side of differentiated cells within the villus-like domain, the MTOC was localized apically, consistent with previous studies on enterocytes ^25,26,27^. Additionally, the majority of the microtubule network was oriented toward the apical side of the villus-like domain (Supplementary Figure 3B). Only a few microtubules were observed along the lateral membranes at the basal side, with no apparent colocalization with AcSs (Supplementary Figure 3C-E). This suggested that microtubules may not be essential for AcS formation. To test this hypothesis, we treated organoid-derived monolayers with nocodazole. Since microtubule depolymerization did not affect the integrity of AcS structures (Supplementary Figure 3F), we concluded that microtubules were not required for AcS development.

Furthermore, we could not detect canonical focal adhesion components, such as paxillin (Supplementary Figure 4A-B) or talin (not shown) at the AcS branches. While paxillin-positive focal adhesions were detected along actin fibres in cells located at the periphery of crypt-like domains, only faint and small paxillin dot patterns were observed in differentiated cells (Supplementary Figure 4A-B), suggesting that AcS branches are not primarily dedicated to cell-substrate adhesion. Subsequently, we examined the spatial arrangement of AcS branches at basal cell contacts. AcS branches from neighbouring cells did not exhibit physical continuity nor penetrate the plasma membrane (Figure 2B, insert). Instead, AcS branches snugly fit into finger-like medial membrane folds of bicellular membranes (Figure 2G-H, Supplementary Figure 4C), wherein E-cadherin-based adhesion sites and actin cross-linkers were localized (Figure 2Hc-e,I-J, Supplementary Figure 4C-E). We further explored the potential mechanical coupling origin of AcSs through basal E-cadherin-based contacts. After chelating extracellular calcium with EDTA treatment, cell-cell contacts were disrupted, which broke down the interconnected network. However, the AcS nodes remain intact (Supplementary Figure 4F). This suggested that while cell-cell adhesion may help maintain the continuity of the actin supracellular network, it may not contribute to AcS node formation. Nevertheless, the arrangement of finger-like and adhesive membranes at the extremities of AcS branches suggested a “zippering” effect at this level, potentially providing structural and architectural stability to the AcS network that connects each epithelial cell within the differentiated domain (Figure 2K).

### Actin star formation relies on significant cell contractility and minimal substrate adhesion

Similar self-organizing structures resembling AcSs have been shown to emerge *in vitro* within minimal actin cortices, upon the addition of myosin-II filaments and permissive contractility ^28,29,30,31^. Consequently, we speculated that cell contractility might be a prerequisite in our system for the generation of AcSs. Analyses of myosin-IIA-KI-GFP organoid monolayers revealed a concentrated GFP signal at the AcS node and its surrounding area (Figure 3A-B). This finding was confirmed with endogenous phosphorylated myosin light chain 2 (P-MLC2) in 2D organoid monolayers (Figure 3C-D) and with endogenous myosin-IIA in adult mouse intestine (Supplementary Figure 5A). Remarkably, a substantial proportion of total cell contractility localised at the AcS level in the basal domain (Figure 3E), with approximately 61.5% of myosin-IIA-GFP and 61.3% of P-MLC2 signal intensity (Figure 3F). This suggests that the bulk of cell contractility accumulates at basal AcS structures. Furthermore, the contractile capacity of the differentiated cell type was particularly elevated compared to the proliferative cells of the crypt-like domain (Figure 3G), with myosin-IIA-GFP and P-MLC2 signal intensities averaging 1.30 and 2.37 times higher, respectively (Figure 3H). Similarly, P-MLC2 signal intensity was significantly lower in crypts compared to villi in adult small intestine (Supplementary Figure 5B-C). These data highlighted a distinct tissue patterning of contractility levels within this epithelium. Interestingly, live imaging analysis indicated that AcSs formed from actin foci that rapidly coalesced in cells that transverse the periphery of the crypt-like domain into the differentiated domain (Supplementary Figure 5D; Video S3). Thus, the emergence of AcS coincided with the transition of cells into a highly contractile tissue domain.

**Figure 3:**
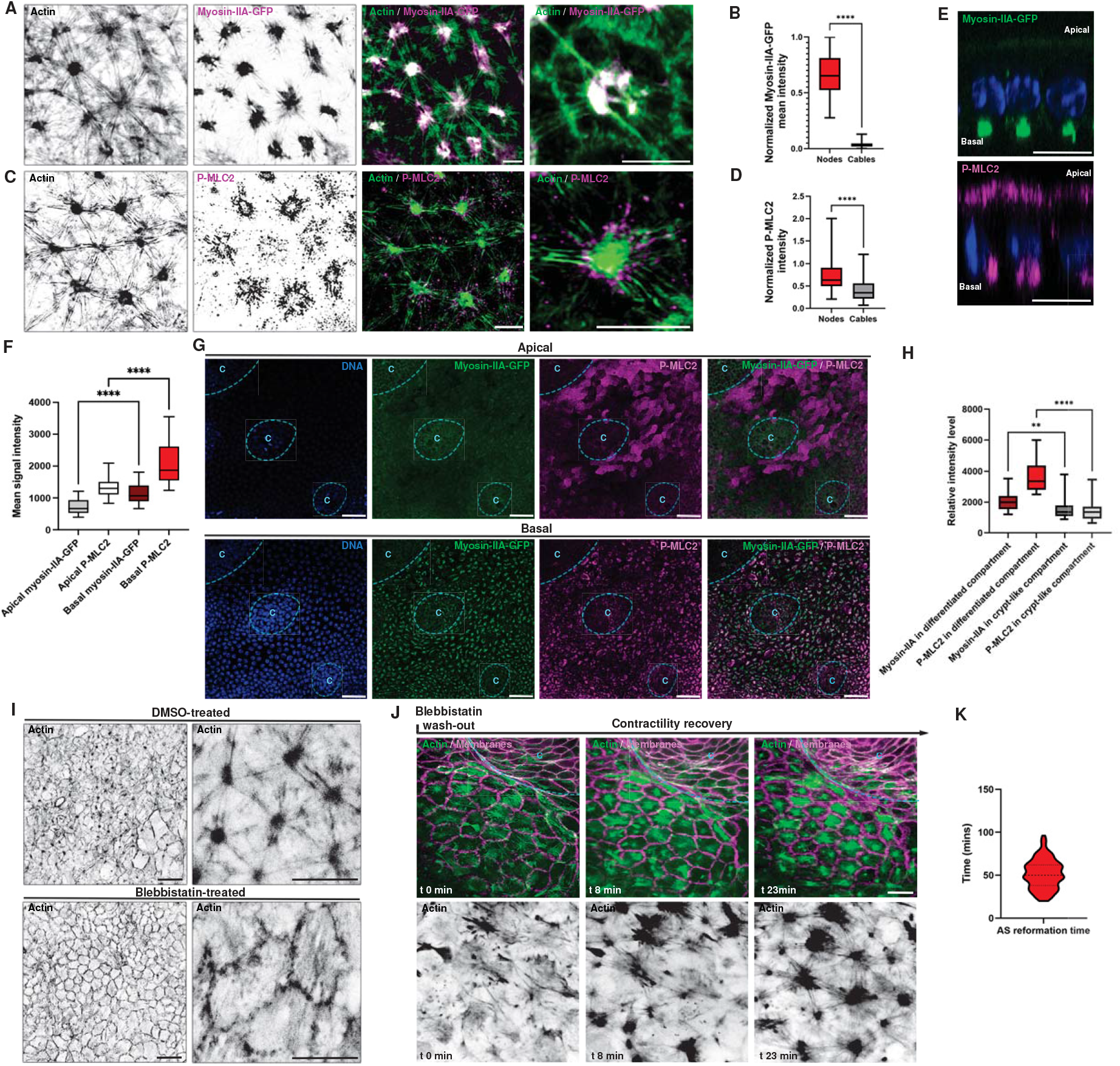
Cell contractility triggers actin star formation. **(A)** Confocal analysis of myosin-IIA-KI-GFP (magenta) localization in AcSs (green). Scale bar, 5 μm. **(B)** Statistical analyses of the signal intensity level of myosin-IIA-GFP in nodes and cables of AcSs. Myosin-IIA-GFP signal intensity in AcS nodes = 0.653 (0.525-0.812) (median (IQR)), in AcS cables = 0.029 (0.018-0.047). N = 3 experiments, n = 154 cells. Mann-Whitney test, ****p<0.0001. **(C)** N-SIM analysis of P-MLC2 (magenta) localization in AcSs (green). Scale bar, 5 μm. **(D)** Statistical analyses of the signal intensity level of P-MLC2 in nodes and cables of AcSs. P-MLC2 signal intensity in AcS nodes = 0.632 (0.496-0.909), in cables = 0.348 (0.216-0.558). N = 3 experiments, n = 98 cells. Mann-Whitney, ****p<0.0001. **(E)** Confocal analysis of the apico-basal distribution of myosin-IIA-GFP (green) or P-MLC2 (magenta) in organoid-derived monolayers. Scale bar, 10 μm. **(F)** Statistical analyses of the signal intensity level of myosin-IIA-GFP and P-MLC2 in the apical and basal domain of differentiated cells. Mean apical myosin-IIA-GFP signal intensity in differentiated compartments = 725±48 (mean±S.E.M), basal myosin-IIA-GFP = 1159±72, apical P-MLC2 = 1314±63, basal P-MLC2 = 2078±135. N = 3 experiments, n (cells in differentiated compartments) = 23 cells. Paired t-tests, ****p<0.0001. **(G)** Confocal analysis of myosin-IIA-GFP (green) and P-MLC2 (magenta) distribution in an organoid-derived monolayer. Nuclei (DNA) are stained with Hoechst33342 (blue). Crypt-like domains (c) are delimited with a dotted blue line. Scale bar, 20 μm. **(H)** Statistical analyses of the signal intensity level of myosin-IIA-GFP and P-MLC2 in differentiated and crypt-like compartments. Myosin-IIA-GFP signal intensity in differentiated compartments = 1987 (1548-2394) (median (IQR)), in crypt-like compartments = 1356 (1088-1791), P-MLC2 signal intensity in differentiated compartments = 3351 (2797-4369), in crypt-like compartments = 1360 (936.5-1701). N = 3 experiments, n (cells in differentiated compartments) = 23 cells, n (cells in crypt-like compartments) = 19 cells. Mann-Whitney, **p = 0.0022, ****p<0.0001. **(I)** Confocal analysis and z-projection of actin distribution in the basal domain of control or blebbistatin-treated organoid-derived monolayers. Scale bar, left panel 20 μm, right panel 10 μm. **(J)** Time-lapse images of CellMask actin (green) in tdTomato (magenta) organoid-derived monolayer after 1h blebbistatin treatment and then wash-out (t = 0min). Crypt-like domains (c) are delimited with a dotted blue line. Scale bar, 10 μm. **(K)** Statistical analysis of the mean time of AcS re-formation after blebbistatin treatment and wash-out. Mean time (min) = 51.41±16.81 (mean±S.D.). N = 3 experiments, n = 101 cells.

Furthermore, we directly probed the necessity of contractility for AcS formation. As illustrated in Figure 1D and Supplementary Figures 1D-E, the crypt compartment lacked AcS structures. However, enhancing contractility through calyculin-A treatment of organoid-derived monolayers led to the formation of AcS-like structures within the crypt-like compartment (Supplementary Figure 6, Video S4). In contrast, treatment with blebbistatin, an inhibitor of actomyosin activity, resulted in the disappearance of typical AcS structures from the basal surface within the villus-like compartment (Figure 3I). Notably, blebbistatin treatment had only a minor impact on the pools of myosin-IIA and P-MLC2 at the apical domains of organoid-derived 2D monolayers (Supplementary Figure 7A-B). Moreover, under downregulation of actomyosin activity, the actin cytoskeleton predominantly organized along the lateral membranes and formed thin stress fibre cables (Figure 3I). Similarly, inducible myosin-IIA-KO caused a significant dismantling of AcSs (Supplementary Figure 7C). In addition, restoring contractility in blebbistatin-treated cells through subsequent wash-out led to the reformation of AcSs in approximately 50 min (Figure 3 J-K; Video S5). Live recording of actin cytoskeleton dynamics during this process revealed a reorganization of actin cables (Video S6). Local inward contraction within each cell triggered the massive coalescence of actin into central foci and the accumulation of small actin bundles into thick radially arranged actin bundles, i.e. the forming AcS node and branches, respectively (Figure 3J; Video S6). Collectively, these findings pointed out the pivotal role of contractility in AcS generation and suggested that the star-like organization of the actomyosin network may be mechanically sensitive.

We then manipulated the mechanical environment of the organoid-derived monolayer by varying substrate rigidities using various polyacrylamide substrates (Matrigel-coated PAA gels) ^32^. AcSs grew on soft substrates with a rigidity of 300 Pa (Figure 4A-B), a rigidity comparable to the *in vivo* condition ^22^. However, stiffer substrates caused basal actin network remodelling, leading to the disappearance of AcSs (Figure 4A-B). Specifically, on 2.4 kPa substrates, there was a shift towards the formation of numerous interconnected actin foci instead of a single actin node, while on 5.2 kPa substrates, stress fibre-like actin bundles predominated (Figure 4A-B). Thus, the mechanical properties of the cell substrate and the resulting level of cell-substrate adhesion seem to be important for AcS formation. Notably, substrate stiffness not only enhanced cell contractility but also promoted cell adhesion, anchoring actin bundles to the substrate and preventing their coalescence into larger bundles ^33^. As substrate stiffness increased, the monolayer also exhibited the formation of prominent focal adhesions, which were connected to actin stress fibres (Supplementary Figure 8). These focal adhesions likely strengthened the cell-substrate coupling, allowing contractile forces to be transmitted to the substrate rather than to neighbouring cells, thereby inhibiting the formation of an interconnected actin network ^34^.

**Figure 4:**
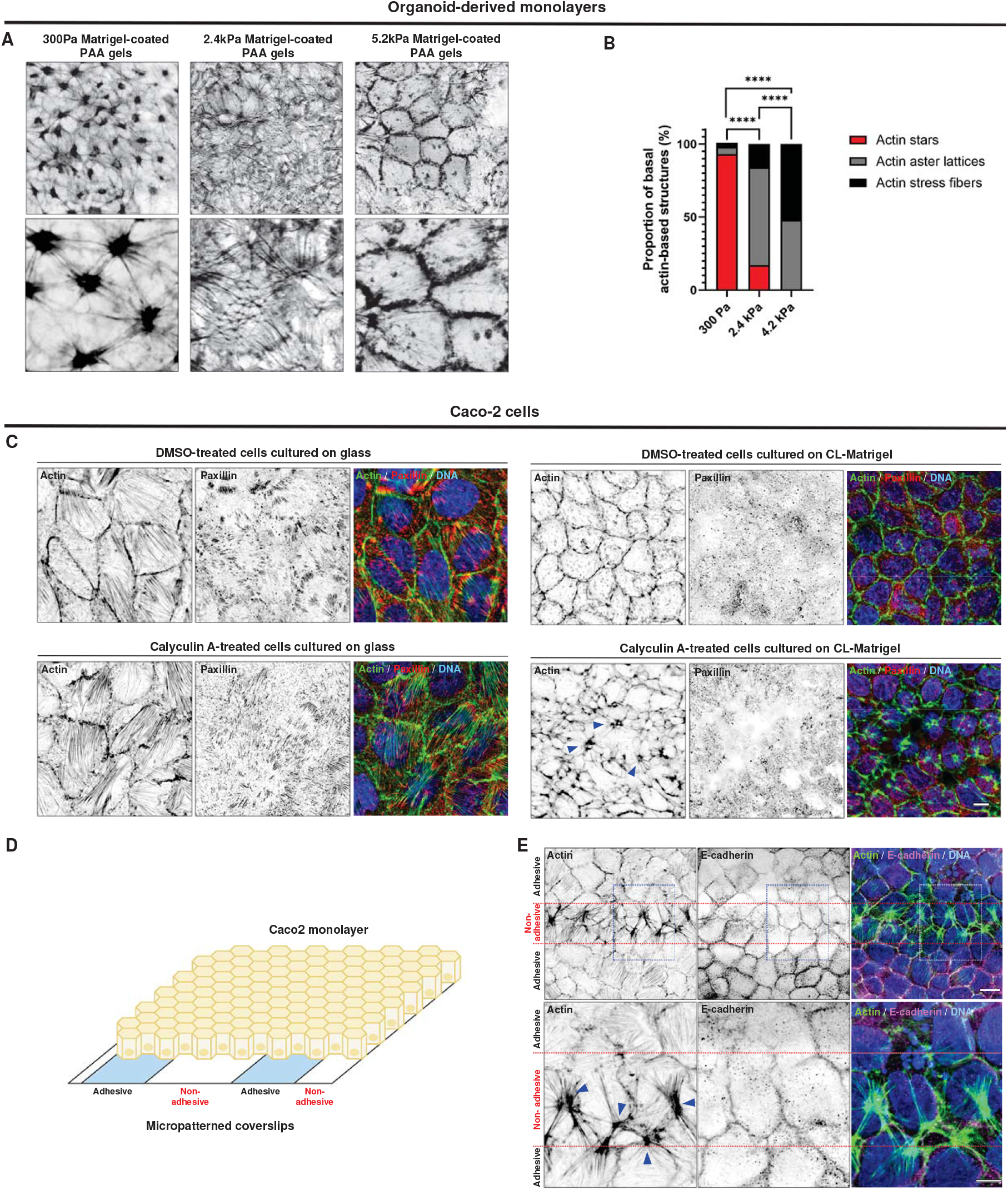
Epithelial mechanical properties and its environment condition on actin stars development. **(A)** Confocal analysis of the basal actin arrangement in organoid-derived monolayer grown on 300Pa, 2.4 or 5.2kPa PAA gels. Scale bar, 10 μm. **(B)** Statistical analyses of basal actin-based structures formed on different rigidity PAA gels. Mean AcS proportion at 300Pa = 93±4.33% (mean+S.E.M), at 2.4kPa = 17±10.02% and at 5.2kPa = 0±0%. Mean actin aster lattice proportion at 300Pa = 5±2.33%, at 2.4kPa = 67±14.58%, at 5.2kPa = 48±14.86%. Mean actin stress fibers proportion at 300Pa = 3±3%, at 2.4kPa = 16±11.12%, at 5.2kPa = 52±14.86% N = 3 experiments, n (300Pa) = 107 cells, n (2.4kPa) = 160 cells, n (5.2kPa) = 229 cells. Fisher’s exact test, ****p<0.0001. **(C)** Confocal analysis of basal actin (green), paxillin (red) and E-cadherin (magenta) in Caco2 cells grown on glass coverslips, uncoated or on cross-linked Matrigel, and treated with either DMSO (control) or 20nM calyculin-A. Cells stained with phalloidin and paxillin show actin (green) or focal-adhesion (red) organization respectively. Nuclei (blue) are stained with Hoechst33342. Scale bar, 5 μm. Blue arrows show AcS nodes in Caco2 cells. Scale bar, 2 μm. **(D)** Scheme showing the micropatterned substrates used for Caco2 monolayer culture in adhesive / non-adhesive conditions. **(E)** Confocal analysis basal distribution of actin (green) and E-cadherin (magenta) in Caco2 cells grown on adhesive / non-adhesive micro-patterns. Red dotted lines delimit the adhesive from the non-adhesive areas. Blue arrows in the inset indicate AcS nodes. Nuclei (blue) are stained with Hoechst33342. Scale bars, upper row 10 μm, lower row 5 μm.

To disentangle the respective roles of cell contractility and adhesion, we employed surface micropatterning to selectively modulate cell adhesion and assess its impact on the contractile bundle assembly. For this purpose, we used the well-established intestinal epithelial cell line, the adenocarcinoma Caco2 cells in which the AcS formation has not been previously documented. To investigate contractility independently of cell adhesion, we cultured Caco2 cells on stiff, adhesion-permissive substrates such as glass, and found canonical stress fibres instead of AcSs (Figure 4C, upper left panel). While Caco2 cells on soft CL-Matrigel substrates ^22^ (∼300 Pa, comparable with the PAA gels used in Figure 4A) show significant loss of stress fibres (Figure 4C, upper right panel), AcS organization was still absent. We then elevated actomyosin contractility through calyculin-A treatment and discovered while no discernible modulation of the basal actin arrangement on the glass substrates (Figure 4C, lower left panel), the enhanced contractility caused the formation of AcS-like structures on soft CL-Matrigel substrates (Figure 4C, lower right panel). Of note, we found Caco2 cells developed large FAs (paxillin staining) on stiff substrates which were largely absent on soft crosslinked Matrigel substrates (Figure 4C). Along this line, we then tested if weakening cell-substrate adhesion might facilitate the development of AcS networks.

We cultured these cells on micropatterned substrates featuring alternating adhesive and non-adhesive regions (Figure 4D). While Caco2 cells displayed conventional stress fibres on the adhesive areas, intriguingly, they developed patterns resembling AcSs on the non-adhesive regions (Figure 4E). These findings suggested that the cell-substrate machinery restricted the physical integration of tensile forces within the basal domain and prevented the coalescence of bundles into a radial AcS network. Altogether, these results emphasized the critical roles of both the mechanical properties of epithelial cells and their surrounding environment in AcS development. They also implied that cell-substrate adhesion limited the coalescence of contractile bundles into an AcS lattice. We concluded that the development of star-like actin structures likely arose from a combination of mechanical properties inherent to the intestinal epithelium: low adhesion to the basal substrate and high cellular contractility.

### The actin star network provides epithelial morphological and dynamical stability

What are AcS’ functions in the epithelial tissue? We found that inhibition of AcS formation with blebbistatin treatment in the organoid-derived monolayer induced a decrease in cell height (Figure 5A-B), indicating a shift from a columnar to a cuboid cell shape in the absence of the AcS network. We did not observe any noticeable effect of blebbistatin treatment on apico-basal cell polarity or on the integrity of tight and tricellular junctions (Supplementary Figure 9A-C). Moreover, the treatment increased basal cell area (Figure 5C-D) and reduced cellular hexagonality, as quantified through a decrease in local triangular order (see Methods; Figure 5G). In addition, local inhibition of AcSs via light-inducible blebbistatin activation resulted in similar cell area modifications (Figure 5E-F). Conversely, the gradual re-establishment of AcSs following blebbistatin treatment wash-out coincided with basal cell shape remodelling, characterized by the reduction of cell area and the acquisition of a more circular and hexagonal form (Supplementary Figure 9D-F; Video S6). Consistently, we observed an enhanced cell density within the epithelial tissue (Supplementary Figure 9G). Hence, AcSs likely contributed to the maintenance of a tightly packed columnar epithelial monolayer. Moreover, as shown in Figure 2B, 2G-J, AcSs contained in each epithelial cell were mechanically connected to each other via finger-like E-cadherin-based junctions on the basal surface, and we thus hypothesized that the AcS network may place the differentiated tissue under a homogenous tension.

**Figure 5:**
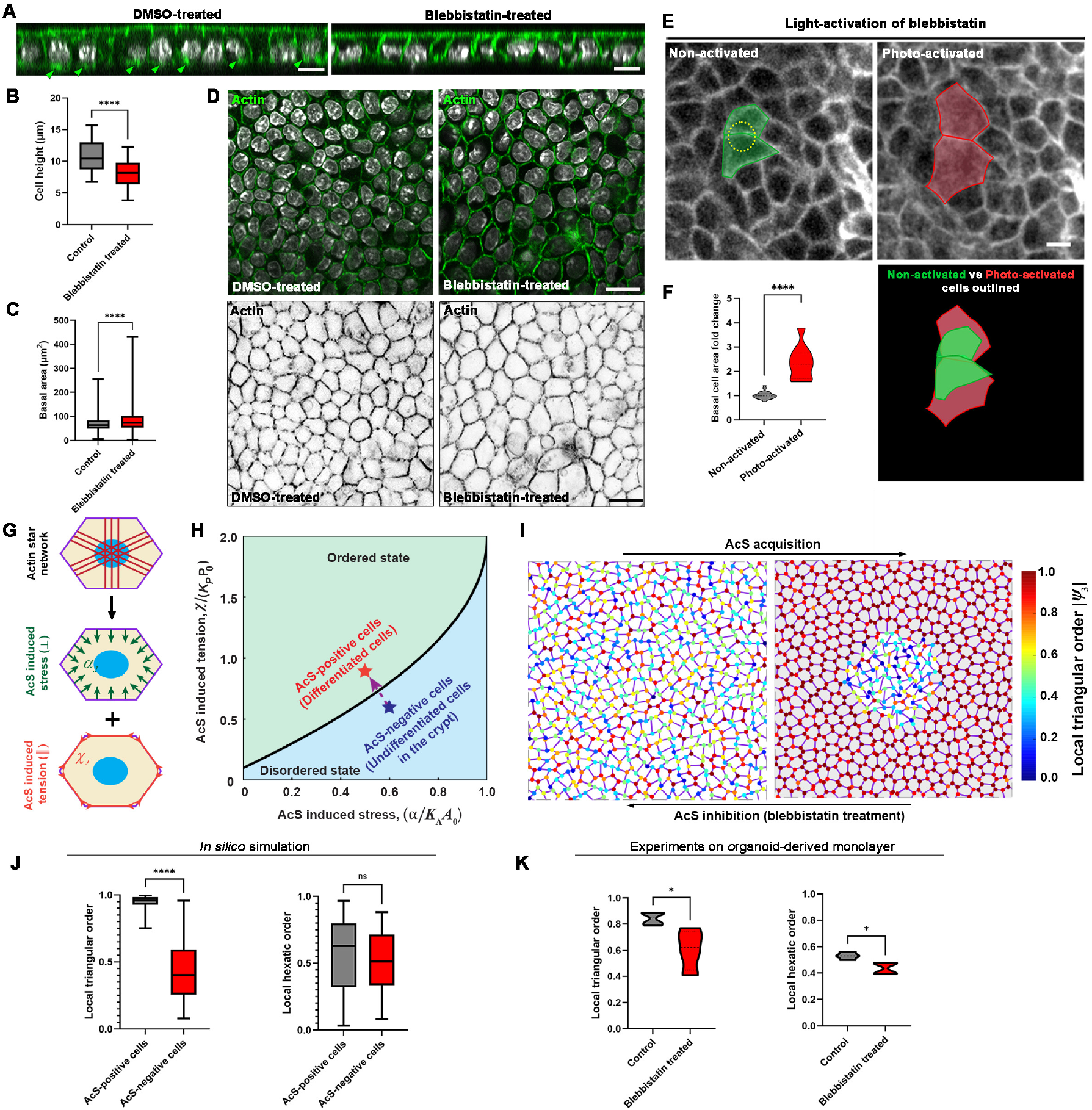
The actin star network ensures epithelial morphological stability. **(A)** Confocal analysis and xz view of actin distribution in DMSO-treated or blebbistatin-treated organoid-derived monolayers. Nuclei (gray) are shown. Green arrowheads point to AcSs. Scale bar, 10 μm. **(B)** Statistical analysis of mean cell height in control or blebbistatin-treated cells. Cell height in control = 10.43 µm (8.731-12.99) (median (IQR)), in blebbistatin-treated = 8.128 (6.392-9.742). N = 3 experiments, n (control) = 60 cells, n (blebbistatin-treated) = 65 cells. Mann-Whitney, ****p<0.0001. **(C)** Statistical analysis of mean basal cell area in control or blebbistatin treated cells. Basal cell area in control = 63 µm^2^ (49.34-82.33) (median (IQR)), in blebbistatin treated = 73.14 (53.30-102). N = 3 experiments, n (control) = 675 cells, n (blebbistatin treated) = 585 cells. Mann-Whitney, ****p<0.0001. **(D)** Confocal analysis illustrating cell shape changes in blebbistatin-treated organoid-derived monolayers compared to DMSO-treated. Top panel shows merged image with actin (green) and nuclei (gray), and bottom panel shows actin in inverted grey. Scale bar, 20 μm. (**E)** Images of membranes-tdTomato before (green) and 10 min after (red) photo-activation of azidoblebbistatin. Scale bar, 5 μm. **(F)** Statistical analysis of the basal cell area modification after photo-activation of azidoblebbistatin. Mean basal cell area fold change in non-activated condition = 1.015±0.03 (mean±S.E.M), in photo-activated condition = 2.353±0.21. N = 3 experiments, n (non-activated) = 23 cells, n (photo-activated) = 10 cells. Unpaired t-test, ****p<0.0001. **(G)** Mechanical description of the AcS network contractility within a cell by AcS-cable stresses. In Supplementary Information, we show that a AcS-cable-tension model (model 2) maps into two contributions, an AcS-induced contractile stress *α* and an AcS-induced contribution to the junction tension *χ*, defining our model 1. **(H)** Theoretical analysis of the order/disorder rigidity transition in terms of the AcS-induced pulling stress *α* and tension *χ*. The black solid line represents the theoretically predicted critical rigidity transition line; see Eq. (S19) in Supplementary Information. The symbols and arrow denote the proposed path of cell differentiation. **(I)** Simulation of the local triangular order parameter upon AcS formation and subsequent optimal cell differentiation, or AcS inhibition under blebbistatin treatment and deficient cell differentiation. **(J)** Statistical quantification of the local triangular and hexatic order *in silico*. Local triangular order in AcS-positive cells = 0.959 (0.927-0.984) (median (IQR)), in AcS-negative cells = 0.402 (0.256-0.592). Local hexatic order in AcS-positive cells = 0.629 (0.321-0.800), in AcS-negative cells = 0.5130 (0.335-0.715). Mann-Whitney, ****p<0.0001. **(K)** Statistical quantification of the local triangular and local hexatic order from experiments. Local triangular order in control cells = 0.847 (0.795-0.885) (median (IQR)), in blebbistatin-treated cells = 0.622 (0.448-0.746). Local hexatic order in control cells = 0.530 (0.505-0.555), in blebbistatin-treated cells = 0.436 (0.393-0.475). Unpaired t-test, *p (triangular) =0.027, *p (hexatic) =0.011.

To test this, we next incorporated our experimental observations into a computational model of the cellular assembly, called vertex model (see Supplementary Information). We first considered a coarse-grained model, where AcS formation triggers a homogeneous contractile response, through a reduction of the preferred cell area and cell perimeter (Figure 5G-H; Supplementary Information, Section I Figure S1), in proportion set by experiments. As in experiments, we observed an increase in the level of hexagonality, as measured through the triangular order, within the cells undergoing the contractile stresses as compared to other cells (Figure 5I-K; Supplementary Information Section II Figure S6G; see Methods, section Computational modelling). This effect was reversible upon the removal of the contractile stress, thus mimicking the effect of blebbistatin. We also considered a second model where AcSs are modelled by a discrete set of force dipoles between the cell-cell junction midpoint and the cell barycentre (Supplementary Information, Section II Figure S3). Such a second model leads to a similar path in the preferred area and perimeter space (see Supplementary Information, section II). These simulations also suggested that the resulting tissue was under high tension. To experimentally assess the level of tissue tension, we next laser-dissected a single AcS and then monitored the behaviour of surrounding AcSs (labelled as actin nodes N1-6) (Figure 6A-B; Video S7). We found an almost immediate recoil of neighbouring AcS nodes after the ablation and existence of a visible recoil at large distances (Figure 6A-C). Notably, laser ablation at the cell-cell junction caused recoil of both actin branches from the neighbouring cells that connect at the junction (Supplementary Figure 10A-C, Video S8). In contrast, ablation at the AcS branch between the cell-cell junction and AcS node resulted in recoil of only the ablated branch, while the branch in the adjacent cell remained intact (Supplementary Figure 10D-F, Video S9). In this case, the junction itself was displaced away from the ablation site (Supplementary Figure 10D-F, Video S9). Together, these experiments indicated that in the differentiated epithelial area, AcSs of neighbouring cells were interconnected, facilitating the transmission of tension across several rows of cells. This tension induced a rapid recoil of the AcS branch post-ablation, occurring at a speed of approximately 5 µm/s (Supplementary Figure 10G). Tension propagation was abolished when the AcS network was disrupted by blebbistatin treatment (Supplementary Figure 11; Video S10). Importantly, laser ablation experiments only affected the basal actomyosin network associated with AcSs, leaving the apical actomyosin organization apparently intact (Supplementary Figure 12). Based on our vertex model simulation, in which a cell is ablated within a tensile tissue, we justified that the recoil speed is informative of the cell-substrate friction (Figure 6D-E). In our simulations, the delay in the strain propagation scaled with the distance to the ablated cell according to τ*∼*L^2/K, i.e. with K an effective diffusion of elasticity that is inversely proportional to the friction to the substrate (Figure 6F). These results showed a diffusion effect of the initial local elastic response after laser dissection and the existence of a mechanical diffusion of elasticity away from the ablation site, testifying of the large-scale connectivity provided by the AcS lattice.

**Figure 6:**
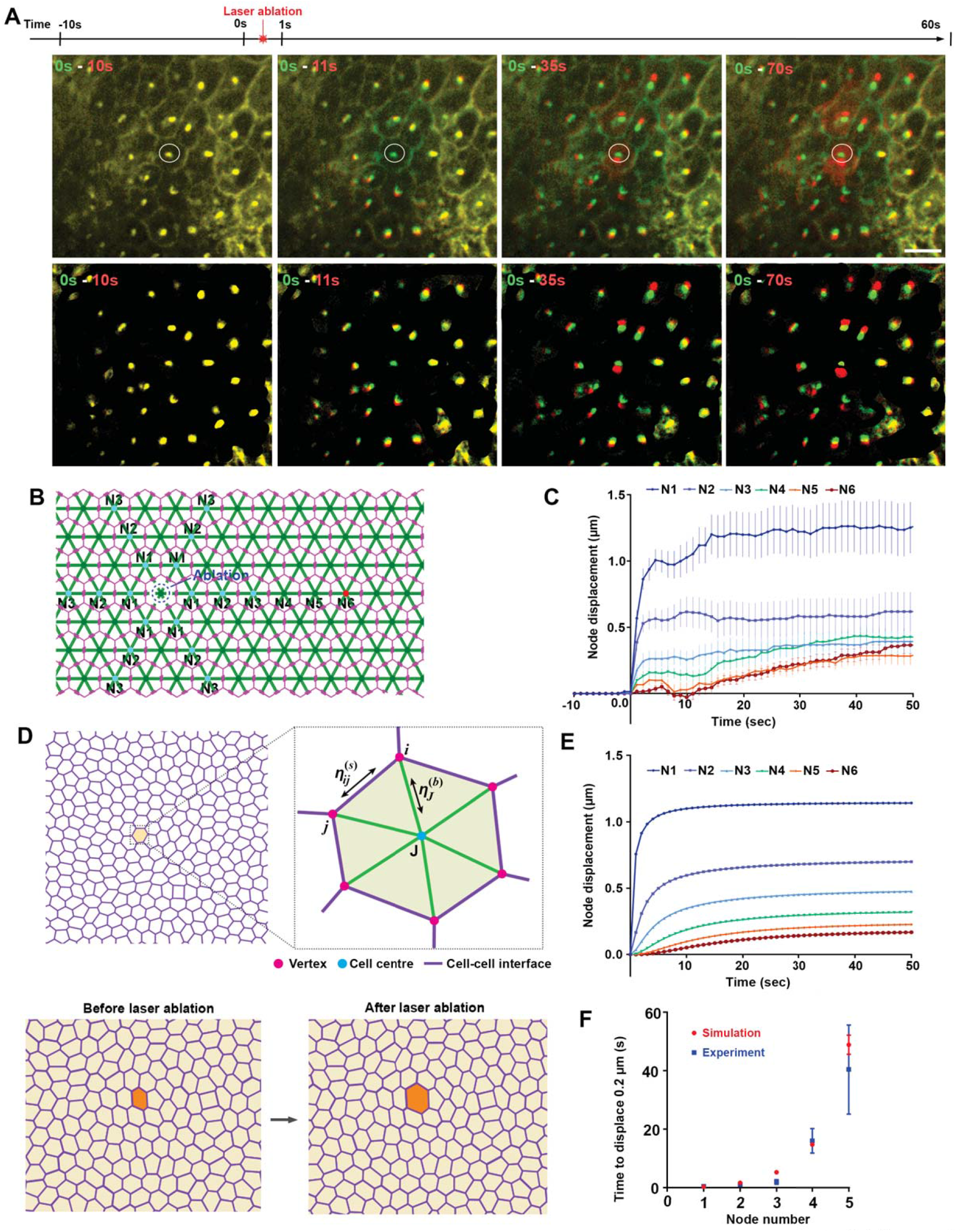
The actin star network is associated with increased level of tension. **(A)** Color-coded time-lapse analysis of AcS displacement after laser ablation of a given AcS (white circle). The first image (t = -10s) is false-colored green, the image at a given time, i.e. before (t = 0s) or after (t = 1, 25 or 60s) laser ablation is false-colored in red. Masks of AcS nodes are also presented in the lower panel. Scale bar, 10 μm. **(B)** Scheme showing the site of laser ablation with dotted circle in the AcS lattice (green) within the epithelial monolayer (magenta), and depicting the AcS nodes (N) annotation for quantification. **(C)** Average radial displacement of AcS nodes that neighbour the laser ablation point. N0, point of AcS laser-ablation. **(D)** Upper panel, sketch of the AcS-induced vertex model with cell-cell junction viscosity. Lower panel, laser ablation protocol: the cell at the model ablation site has its activity set to zero. **(E)** Average radial displacement of cell barycentres at the *N*-th row as a function of time *t* after ablation (see Methods). **(F)** Time to reach 0.20 μm radial displacement, *t*_*0*.*2µm*_, (mean ± S.E.M) as a function of the row of cells (i.e., distance to the laser ablation site), in experiments (blue, *n* = 13 – 18 tracked AcS nodes, N = 3 experiments) and simulations (red, averaged over *n* = 5 independent simulations).

Furthermore, we observed that the blebbistatin-treated organoid-derived monolayer displayed numerous and large basal cell protrusions resembling lamellipodia in the villus-like domain (pseudo-coloured in green, Figure 7A-B, t = 0 min; raw images in Supplementary Figure 13). Upon recovery of contractility following blebbistatin wash-out, AcSs re-growth occurred (as discussed earlier), which was accompanied by the gradual disappearance of basal lamellipodia (Figure 7A-B t = 2 to 25 min; Supplementary Figure 13; Video S6). We thus propose that AcSs might play a role in inhibiting basal cell dynamics. To directly test this hypothesis, we proceeded to laser dissection of a specific AcS and monitored its impact on the basal cell surface (Figure 7D; Supplementary Figure 14A). At 125s after AcS removal, the actin network reorganized and exhibited multiple actin foci within the cell which was targeted by laser dissection, indicating that AcS had not yet reformed. During this period, the basal cell surface expanded and exhibited actin-positive cell protrusions (Figure 7D; Supplementary Figure 14A). Indeed, the perimeter of basal protrusive extension in the laser-dissected cell increased from 2.6±2.6 μm in the pre-ablation condition to 75.8±3.7. μm after the ablation (Figure 7C; Video S11-S12). When two adjacent cells were subjected to laser ablation, both exhibited similar responses, characterized by basal expansion and the formation of actin protrusions (Supplementary Figure 14B, Video S13). When half of the actin branches connected to an AcS node were ablated, only the ablated half produced actin protrusions (Supplementary Figure 15, Video S14). Moreover, the nodes in neighbouring cells did not disassemble, but instead moved significantly away from the cell centroid (Supplementary Figure 16A-B). Based on these findings, we concluded that the AcS lattice restricted epithelial basal protrusive activity by placing the tissue under tension. These data prompted us to explore the participation of the AcS network in global epithelial tissue behaviour. As compared to the control condition, a blebbistatin treatment led to more persistent cell trajectories (Figure 7E), and more correlated velocities, both temporally (*D*, see Methods, Figure 7F-I), and spatially (*λ*_WT_ = 40 +/-20 *μ*m against *λ*_Blebbistanin_ = 70 +/-15 *μ*m), see *Methods*, Figure 7J). These observations of an increase in the velocity correlations *D* and *λ* hint at not coherent with a lower tension in blebbistatin. However, as predicted by the generic theory of Henkes et al. ^35^, these are coherent with the observed concomitant increase in the protrusion persistence time.

**Figure 7:**
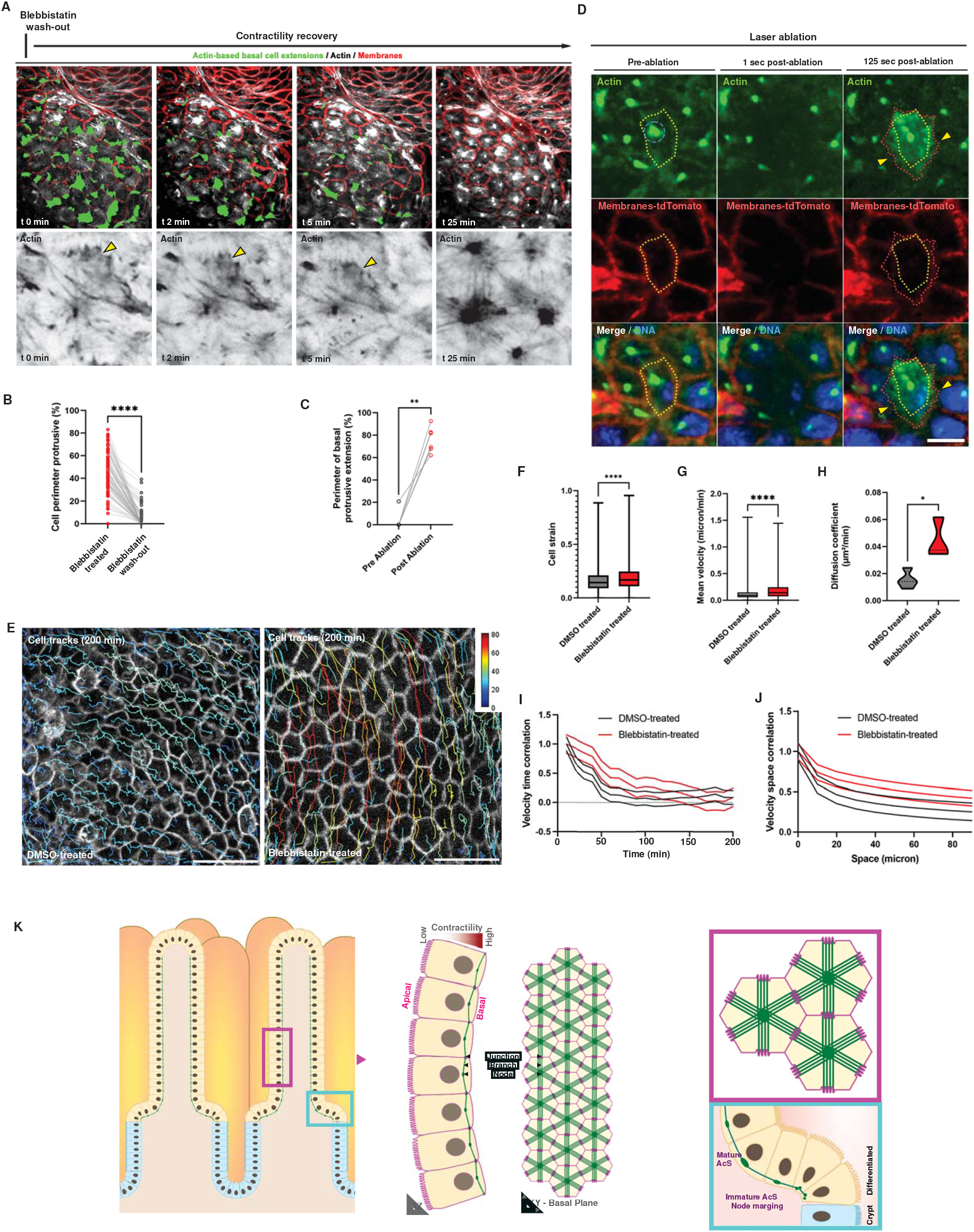
The actin star assembly restricts epithelial dynamics. **(A)** Time-lapse of CellMaskActin (white) and membranes-tdTomato (red) organoid-derived monolayer after 1h blebbistatin treatment and then wash-out (t = 0 min). Actin-based protrusive structures are outlined in green, and also shown in the lower black and white panel. Yellow arrowheads point to actin-based protrusive extension. Scale bars, 10 μm. **(B)** Statistical analysis of cell perimeter protrusive before and after blebbistatin wash-out. P erimeter protrusive before = 41.01 (32.04-55.07) µm (median (IQR), after = 0.000 (0.000-3.491). N = 4 experiments, n = 100 cells. Wilcoxon test, ****p<0.0001. **(C)** Statistical analysis of the perimeter of basal protrusive cell extension after laser ablation of the AcS node. P erimeter in pre-ablation condition = 0.000 (0.000-0.000) µm (median (IQR), in post-ablation condition = 75.63 (67.83-82.60). N = 3 experiments, n = 8 cells. Wilcoxon test, ****p<0.0001. **(D)** Time-lapse of CellMaskActin (green) and membranes-tdTomato (red) before or after laser ablation (blue dotted circle) of an AcS node. Cell perimeter is outlined before ablation in yellow dotted line and after ablation in red dotted line. Yellow arrowheads point to actin-based protrusive extension. Nuclei are stained in blue. Scale bar, 5μm. **(E)** Automated cell tracking in DMSO- or blebbistatin-treated tdTomato organoid-derived monolayers, and t-projection of 20 frames time-lapse series. Monolayer background corresponds to t = 0. Color bar indicates the track length (µm)). Scale bar, 50 μm. **(F)** Statistical analysis of the mean cell strain in DMSO-treated or blebbistatin-treated organoid-derived monolayers. C ell strain in DMSO-treated cells = 0.143 (0.096-0.211) (median (IQR)), in blebbistatin-treated = 0.169 (0.109-0.247). n (DMSO-treated cells) = 95863, n (blebbistatin-treated cells) = 21950. Mann-Whitney test, ****p<0.0001. **(G)** Statistical analysis of the mean cell velocity in DMSO-treated or blebbistatin-treated organoid-derived monolayers. Velocity in DMSO-treated cells = 0.092 μm/min (0.056-0.145) (median (IQR)), in blebbistatin-treated = 0.141 µm/min (0.073-0.244). n (DMSO-treated cells) = 3116, n (blebbistatin-treated cells) = 1622. Mann-Whitney test, ****p<0.0001. **(H)** Statistical analysis of the mean diffusion coefficient in DMSO-treated or blebbistatin-treated organoid-derived monolayers. Mean diffusion coefficient in DMSO-treated cells = 0.015±0.003 (mean±S.E.M), in blebbistatin-treated = 0.044±0.009. n (DMSO-treated cells) = 4, n (blebbistatin-treated cells) = 3. Unpaired t-test, *p=0.0159. **(I-J)** Analysis of velocity correlations in DMSO-treated or blebbistatin-treated organoid-derived monolayers: **(I)** autocorrelations, as a function of time, **(J)** same-time correlations, as a function of the point-to-point distance (see Methods). N = 4 in DMSO-treated and N = 3 in blebbistatin-treated experiments. **(K)** Scheme depicting the proposed model of AcS development in the mammalian intestinal epithelium.

## DISCUSSION

### Actomyosin star-shaped assemblies develop in differentiated intestinal epithelial cells

In this study, we unveil a supracellular star-shaped organization of actin network in mammalian intestinal epithelial cells and demonstrated its implication in various cellular and tissue functions (Figure 7K). To our knowledge, such actin meshwork development has not been previously documented in mammals, and contrast to the conventional basal actin organization, which predominantly exhibit stress fibres and/or lamellipodia ^36,37,38^. However, some studies have reported unusual basal actin organization in specific contexts. In rat embryonic cells, geodesic-like arrangements of multiple actin foci transiently form before the formation of stress fibres, under the control of actin cross-linking factors (i.e. alpha-actinin, tropomyosin) ^39^. Similar polygonal actin meshes, named cross-linked actin network (CLANs), organize in individual human trabecular meshwork cells ^40,41^. In the same line, super-resolution imaging has shown a multitude of actin foci cortex organization within individual mouse embryonic stem cells ^42^. Importantly, these structures did not denote a multicellular organization and were not regulated by myosin-II-mediated contractility; instead, they relied on actin polymerization. In addition, when mouse embryonic fibroblasts were cultured on non-adhesive/adhesive micropatterns, dynamic actin foci network developed on non-adhesive areas and were exacerbated under inhibition of actin polymerization ^43^. Bershadsky and colleagues then showed through modelling that in this model system, myosin II-mediated contractility propels small actin assemblies towards each other, converging into a single large actin node ^43^. Authors then hypothesized that multiple actin foci network may provide cytoskeletal connectivity to distal parts of the cell ^43^. Furthermore, star-like actin networks have been previously reported *in silico* cell-free systems ^28,44,45^, where authors demonstrated that incorporation of myosin-II-based contractility on an F-actin monolayer induces the formation of a star-shaped actomyosin network and places the actin array under a ‘dynamical steady state’ ^28,44,45^. Altogether, these studies underline how plastic the basal actin network can be in response to environmental factors ^46^. They also pinpoint two pivotal factors: contractility and competition with cell-substrate adhesion, which may explain why AcS organization has not been observed yet in mammalians *ex vivo*. One possibility is that the conventional substrates employed in routine cell culture experiments are glass or plastic, which are exceedingly rigid and foster robust focal adhesions ^47,48,49^, impeding the formation of AcSs. In our experiments, we cultivated organoid-derived monolayers under conditions that closely emulate the *in vivo* rigidity, significantly softer than a typical cell culture dish ^22^. However, using soft substrates alone was not enough to facilitate the formation of AcSs in the cancerous Caco2 cell line (Figure 4C). Another critical factor is the degree of contractility activation. Indeed, elevating contractility on a soft substrate provoked the appearance of AcSs in Caco2 cells. Similarly, intestinal differentiated compartments with heightened contractility, developed the AcS network, while low-contractile crypt compartments lacked AcSs (Figure 3G-H, Supplementary Figure 5B-C). Moreover, the AcS network was exclusively formed when organoid-derived monolayers were grown on soft substrates that hinder geodesic-like arrangements of multiple actin foci or mature focal adhesion formation (Figure 4A-B). Additionally, Caco2 monolayers displayed AcS formation under weakly adherent culture conditions solely when elevated contractility activation is induced via calyculin-A treatment (Figure 4E). Even the low-contractility crypt compartment displayed AcS formation when contractility was enhanced through calyculin-A treatment (Supplementary Figure 6, Video S4). It is tempting to suggest that these transformed/cancerous cells might have lost their innate capacity to promote optimal contractility, at least on the basal side when grown on soft substrate and in a weakly adherent state. Subsequent experiments will be imperative for definitely clarifying this hypothesis.

### The basal actin star assembly generates a supracellular connecting network

The AcS network may represent a distinctive multicellular mechanical entity within the basal region of differentiated epithelial cells. AcS self-organized structures were reminiscent of the apico-medial actomyosin, well-described in the Drosophila epithelium and which is required for cell shape remodeling through apical constriction ^14,15,16,17,18^. Here, while the apical actin network remained localized to the subcortical zone behind adherent and tight junctions ^11^, the basal contractile AcS network extended in a more centripetal manner in intestinal epithelial cells at equilibrium, yet remained interconnected with E-cadherin-based junctions (Figure 2). When tension increases on the apical side, an apical-medial actomyosin network orchestrates tissue folding ^14,15,16,17,18^. During intestinal development, apical constriction also occurs in mammalian intestine for crypt invagination ^50,51,52^. It might be tempting to draw parallels with basal AcSs in differentiated intestinal epithelial cells, which naturally colonize curved geometry of villi in adults (Figure 1A), but AcSs also developed on flat surfaces in *in vitro* cultures (Figure 1D). This implies that they must serve functions beyond differentiated tissue folding.

In contrast to other epithelia, we and others showed that the majority of cell contractility is localized on the basal side of differentiated intestinal cells ^52^ (Figure 3E-F; Supplementary Figure 5B-C). Among the main differentiated gut cell types, the epithelial apical side gets specialized for nutrient absorption with the creation of a brush border of microvilli in enterocytes or for mucus secretion in goblet cells ^21,53^. Although brush border anchors in the terminal web composed of actomyosin and intermediate filaments, this is a robust apical assembly that undergoes little remodelling in physiological conditions once formed, and for which a role for its contractile property outside of the context of brush border formation remains elusive ^54,53,55^.

Star-shaped actomyosin assemblies on the basal cell surface have only rarely been described in epithelia. An ultrastructural study conducted by M. De Ceccatty in 1986 focusing on morphological investigations of sponge epithelial cytoskeletons, reported a similar star-shaped cytoskeletal organization on the basal epithelial layer ^56^. Interestingly, our study echoes a recent study conducted by Harvey and colleagues ^57^. These authors have also shown the presence of a basal-medial actomyosin network in Drosophila epithelium, which appears to be related to our AcS network. However, this study lacks a detailed characterization of basal actomyosin structures and their multicellular connectivity property, as the primary focus was on investigating a newly identified basal cell-cell adhesion complex named ‘basal spot junctions’ ^57^. In the differentiated intestinal epithelium, AcS branches connected E-cadherin-based basal contacts that localized at structures between engulfed fingers and intercellular bridges with basolateral cell-cell contacts, such as discussed by Svitkina and colleagues ^58^, testifying that AcS and basal contacts were under substantial tension. In addition, the presence of mechanosensitive elements like E-cadherin, β-catenin, α-catenin, vinculin, and α-actinin further hints towards the mechano-responsive nature of this network. The actin star network may thus represent a new way to create a supracellular actomyosin organization to transmit force imbalance through the tissue.

### The actin star network constitutes a new mechanical apparatus for epithelial tissue organization and coordination

The AcS network represents a mechanical subunit at the basal cell side that connects surrounding epithelial cells. As such, it exerted functions at cellular and multicellular levels. At the cellular level, the AcS cytoskeletal lattice conferred morphological and functional stability to cells. It modulated cell shape by constraining the basal cell surface, thereby enhancing the columnar appearance and refining cell shape (Figure 5A-F; Supplementary Figure 9D-G). Additionally, it maximized epithelial cell packing and order (Figure 5G-K), which represents an important factor for stabilizing the epithelial assembly and optimizing the intestinal surface for nutrient absorption. Moreover, the disappearance of basal cell protrusive activity upon the formation of the AcS network suggested that AcSs restricted lamellipodia formation. Thus, the AcS network may facilitate cell transition from a motile, fluid-like, to a static, solid-like, morphology, as observed in mature epithelia ^20^. To characterize this transition, we first considered the hexatic order which is a well-known measure in condensed matter ^59^. Nevertheless, here we found that the triangular order, rather than the hexatic one, was the best probe of the AcS formation in experiments, and of the onset of an AcS-induced tension in simulations: the spatial correlation function *g*_6_(*r* = *r*_*i*_ − *r*_*j*)_ = ⟨*ψ*_6_ (*r*_*i*_)*ψ \**_6_ (*r*_*j*_)⟩/ ⟨*ψ*_6_ (*r*_*i*_)*ψ \**_6_ (*r* )⟩ decays over a single cell length even in control experiments and does not exhibit strikingly different behaviour as a function of *r* in our simulated differentiated and undifferentiated cases (see Supplementary Information Figure S7).

Beyond its cellular functions, the AcS network adopted a central role in orchestrating tissue architecture and homeostasis. By enhancing spatial coupling of differentiated epithelial cell clusters, it ensured the mechanical continuity of the epithelial monolayer, thereby upholding tissue integrity. Furthermore, the cytoskeletal network regulated cell shape, promoting a uniform cell hexagonalization that modulates an epithelial order akin to a close-to-solid phase ^20^. By constraining lamellipodia, AcSs hampered cell dynamics, pivotal for the formation of a tissue at equilibrium. This restriction mitigated random cell movement within the monolayer, leading to a relatively homeostatic and uniformly structured tissue architecture. In addition, at a broader tissue scale, the AcS network interconnected differentiated cells within the villus-like domain, and may facilitate the controlled collective cell movement observed along the crypt-villus axis ^60^. An earlier study on gut epithelia turnover reported it to be mediated by actin-rich basal protrusions driven collective cell migration ^61^. There, it was also suggested that front-back polarity in the migrating cell is mediated by basal protrusions. Here, while observing the endogenous basal actin along the villi, we were unable to detect lamellipodial structures in the enterocytes. The majority of enterocytes were uniformly hexagonal and lacked front-back polarity. There was merely an evolution of the enterocyte basal cell shape at the very tip of the villus, which could resemble lamellipodial activity at that point (not shown) and might represent some migrating cells.

In summary, we describe here a unique tissue-spanning multicellular organization of star-shaped actin networks in differentiated basal mammalian epithelium. Along the line of the De Ceccatty’ hypothesis of “histoskeleton” in sponge ^56^, we believe that the AcSs represent a cytoskeletal network for coordination of mammalian epithelial tissue organization and its functioning. Our *in vivo* observations, as well as *ex vivo* and *in vitro* experiments, further confirmed the presence of this network in the intestinal epithelium, its mechanism of formation, and its essential role in tissue organization. We demonstrated that this network emerged under a high contractility / low cell-substrate adhesion state that naturally existed in the differentiated intestinal epithelium. We further showed its importance in regulating various functions from cell to tissue levels, that ensured tissue homeostasis and maintain healthy epithelial architecture. Together, this hints towards a mechanism of AcS network-mediated complex self-organization of the differentiated tissue.

## Acknowledgements

We thank Renata Basto, Véronique Marthiens, Ana-Maria Lennon-Dumenil, Robert S. Adelstein and Danijela Vignjevic for providing mice. We thank Pierre-François Lenne for help with laser ablation equipment. We thank Arnaud Echard (Institut Pasteur, Paris), Guillaume Salbreux (UNIGE, Geneva), Benoit Ladoux, René-Marc Mège, Guillaume Romet-Lemonne and Antoine Jegou (IJM, Paris) for helpful discussions. Confocal microscopy and spinning-disc analyses were performed in the ImagoSeine microscopy facility (IJM) and PICSL imaging facility (IBDM). This work was supported by grants from the Groupama Foundation – Research Prize for Rare Diseases 2017 (to D.D), France 2030, the French Government program managed by the French National Research Agency (ANR-16-CONV-0001) and from the Excellence Initiative of Aix-Marseille University - A*MIDEX (to J.F.R.), as well as by ANR-20-CE30-0023 (COVFEFE) (to J.F.R.), the Fondation ARC (to A.B., M.S and D.D), the Fondation pour la Recherche Médicale FRM (to A.B.), the Marie Skłodowska-Curie Actions, Postdoctoral Fellowships, project 101108750 (to A.B.), the Marie Skłodowska-Curie Actions Postdoctoral Fellowship, project 846449 (to W.X.), the Université de Paris IdEx UP 2021-I-050 funded by the French Government through its “Investments for the Future” (to D.D.), the ANR-19-CE13-0014-01 (to D.D.), the ANR-20-CE13-0015 (to D.D.), the Human Frontier Science Program (RGP0038/2018) (to D.D.), and the INCA PLBIO20-150 – Cancéropole Ile-de-France (to D.D.) and the CNRS through the MiTi interdisciplinary programs (to D.D.).

## Supplemental Information

Supplementary Figures S1-S16, Videos S1-S14, and Supplementary Information file detailing the vertex model simulation procedure are in the online version of the paper.

## Author contributions

A.B., M.S., W.X., S.Z.L., M.K., E.B., S.R., B. L., C.C., D.B., M.T., J.F.R. and D.D. designed and performed experiments and required analyses. A.B., M.S., S.Z.L., M.K., D.B., F.R., M.T., J.F.R. and D.D. coordinated the overall research and experiments, and wrote the manuscript.

## SUPPLEMENTARY VIDEOS

**Video S1:** Z-stack confocal imaging of actin distribution through the differentiated epithelial monolayer of the villus compartment in the adult mouse small intestine. Left panel, distance between z-slices, 0.42 µm. Scale Bar, 20 µm. Right panel, distance between z-slices, 0.23 µm. Scale bar, 5 µm.

**Video S2:** Z-stack confocal imaging of actin distribution through the proliferative epithelial monolayer of the crypt compartment in the adult mouse small intestine. Distance between z-slices, 0.23 µm. Scale bar, 10 μm.

**Video S3:** Time-lapse imaging of CellMask-Actin-GFP (green) and Membranes-tdTomato (magenta) during cell exit from the crypt-like domain and cell entry in the differentiated domain of organoid derived-monolayers. White arrowheads point toward one AcS formation. Crypt-like domains are delimited in yellow circle. Right side shows magnified region of interest. Scale bars, 10 μm (left) and 5 µm (right). Images were acquired every 3 min. Frame rate is 4 fps.

**Video S4:** Time-lapse imaging of CellMask-Actin-GFP (green), Membranes-tdTomato (magenta) and nuclei (DNA, Blue) in organoid derived-monolayers during 50nM calyculin-A treatment. Scale bar, 10 μm. Images were acquired every 3 s. Frame rate is 10 fps.

**Video S5:** Time-lapse imaging of CellMask-Actin-GFP (green) and Membranes-tdTomato (magenta) in organoid derived-monolayers after blebbistatin treatment and subsequent wash-out. Scale bar, 20 μm. Images were acquired every 120 s. Frame rate is 10 fps.

**Video S6:** Time-lapse imaging of CellMask-Actin-GFP (inverted grey or green) and Membranes-tdTomato (magenta) in organoid derived-monolayers after blebbistatin treatment and subsequent wash-out. Membranes-tdTomato (magenta) organoid derived monolayer was stained with CellMask-Actin-GFP and was treated for 1 hr with 10 µM Blebbistatin. Movie was recorded immediately after replacing blebbistatin treatment with blebbistatin free media. Scale Bar, 10 μm. Images were acquired every 120 s. Frame rate is 10 fps.

**Video S7:** Time-lapse imaging of CellMask-Actin-GFP (grey) in organoid derived-monolayers after laser ablation of around a single AcS node along the yellow circle. Tracked AcS nodes trajectories from ablation center are indicated with colored lines, with node ‘1’ (N1) in red indicating immediate neighbour to the ablated node, N2 in green neighbouring N1, N3 in cyan neighbouring N2, N4 in blue neighbouring N3, N5 in magenta neighbouring N4, N6 in yellow neighbouring N5. Scale bar, 10 μm. Images were acquired every 1.1 s. Frame rate is 10 fps.

**Video S8:** Time-lapse imaging of CellMask-Actin-GFP (green or inverted grey), Membranes-tdTomato (magenta) in organoid derived-monolayers during cell-cell junction laser ablation. Laser ablation area is indicated with yellow line. Scale bar, 3 μm. Images were acquired every 0.5 s. Frame rate is 10 fps.

**Video S9:** Time-lapse imaging of CellMask-Actin-GFP (green or inverted grey), Membranes-tdTomato (magenta) in organoid derived-monolayers during laser ablation on AcS branch between AcS node and cell-cell junction. Laser ablation area is indicated with yellow line. Scale bar, 3 μm. Images were acquired every 0.3 s. Frame rate is 25 fps.

**Video S10:** Time-lapse imaging of 1 hr 10 µM blebbistatin treated organoid derived-monolayers during laser ablation of around the yellow circle. Actin is stained with CellMask-Actin-GFP (green), Membranes-tdTomato (magenta) shows cell membrane. Scale bar, 10 μm. Images were acquired every 5 s. Frame rate is 10 fps.

**Video S11:** Time-lapse imaging of CellMask-Actin-GFP (inverted grey or green), Membranes-tdTomato (red) and Hoechst 33342 stained nucleus (blue) in organoid derived-monolayers after laser ablation around a single AcS node indicated with orange circle. Scale bar 10 µm. Images were acquired every 1.56 s. Frame rate is 10 fps.

**Video S12:** Time-lapse imaging of CellMask-Actin-GFP (inverted grey or green) and Membranes-tdTomato (magenta) in organoid derived-monolayers after laser ablation around a single AcS node indicated with yellow circle. Scale bar 5 µm. Images were acquired every 1 s. Frame rate is 33 fps.

**Video S13:** Time-lapse imaging of CellMask-Actin-GFP (inverted grey or green) and Membranes-tdTomato (magenta) in organoid derived-monolayers after laser ablation around AcS node in two neighbouring cells. Scale bar 5 µm. Images were acquired every 1 s. Frame rate is 10 fps.

**Video S14:** Time-lapse imaging of CellMask-Actin-GFP (inverted grey or green) and Membranes-tdTomato (magenta) in organoid derived-monolayers after laser ablation of half

of the branches connected to one AcS node. The ablation site is marked with a yellow line Scale bar 5 µm. Images were acquired every 1 s. Frame rate is 10 fps.

## MATERIAL AND METHODS RESOURCE AVAILABILITY

### Lead contact

Further information and requests for resources and reagents should be directed and will be fulfilled by the lead contact Delphine Delacour.

### Materials availability

Materials generated in the current study are available from the lead contacts upon request. There are restrictions to the availability of due to collaborations or MTAs.

### Data and code availability

Source data files and codes have been submitted as supplementary files. Microscopy data reported in this paper will be shared by the lead contact on request. Custom codes are deposited on GitHub (https://github.com/DelphineDelacour/EpiOrderSeg2). Any additional information required to re-analyze the data reported in this paper is available from the lead contact upon request.

## METHOD DETAILS

### Mice

Wild-type C57/Bl6 adult male mice were provided by the animal house facility of the Institut Jacques Monod and by Janvier Labs company (France). Mice were housed in EOPS (Environment without Specific Pathogenic Organisms) environment, and handled in accordance with French regulation for animal care. Centrin-1-GFP / H2B-mCherry organoids were generated from mice provided by Renata Basto (Institut Curie, Paris) ^62,63^. VillinCreERT2-tdTomato organoids were generated from mice provided by Danijela Vignjevic (Insitut Curie, Paris) ^61,64^. Myosin-IIA-GFP-knock-in organoids ^65^ were generated from mice provided by Robert S. Adelstein (NHLBI, Bethesda) and Ana-Maria Lennon-Dumesnil (Institut Curie, Paris). Myosin-IIA-KO/mTmG mice were kindly provided by Danijela Vignjevic (Institut Curie, Paris) ^61,64^, and generated by crossing myosin-IIA-KO ^66^ and mT/mG mice ^67^.

### Organoid preparation and culture

6 to 12 weeks-old mice were used for organoid generation. After euthanization by cervical dislocation, the small intestine was harvested, flushed with PBS to discard luminal content and cut longitudinally open. The tissue was then cut into small pieces of 3-5 mm and further washed in PBS.

For organoid preparation, the pieces of intestinal tissue were then incubated on ice for 10 min in a tube containing 5 mM EDTA. The tube was then vortexed for 2 min to release villi from the tissue. After EDTA removal, the intestinal pieces were placed in cold PBS and vortexed vigorously for 3 min to ensure crypt release. This process was repeated 3 times, with each fraction recovered. The third and fourth fractions are usually concentrated in crypts, so these are combined and passed through a 70-µm cell strainer to remove remaining villi and centrifuged at 1000 RPM for 5 min. The pellet (crypts) was then washed in advanced DMEM/F12 (#12634010 Thermo Fisher Scientific, Waltham, Massachusetts, USA) and centrifuged. The final pellet is resuspended in 50 µl of 1:1 ratio of advanced DMEM/F12 and ice-cold Matrigel (#734-1100 VWR, Radnor, PA, USA) and plated as domes. Incubation at 37°C for 20-30 min allowed Matrigel polymerization. 3D organoid culture was performed in IntestiCult™ Organoid Growth medium (#06005 StemCell Technologies, Vancouver, Canada), from here on termed ENR medium. Organoid stocks were routinely grown in Matrigel with IntestiCult™ Organoid Growth medium and passaged every 7 to 10 days. Medium was changed every 2 days.

For 2D organoid monolayer preparation, 3D organoids were cultured in L-WRN conditioned media for at least 3 days before use. Organoids were harvested with cold advanced DMEM/F12 and transferred in a falcon tube. Organoids were mechanically broken through a P200-filtered tip 150 times. The solution of broken organoids was centrifuged at 72g for 3 mins at 4°C. The supernatant was removed and 5 mL of fresh F12 was added. The breaking step and centrifugation were repeated once. Then the cell pellet was filtered through a 30 µm cell strainer. The pellet was resuspended in warm L-WRN conditioned media + 10µM Y27632 (StemCell #72302) and 150 µL were gently seeded on the crosslinked Matrigel substrate (18 mm coated coverslips on a 12 well plate) making sure the solution stays on the coverslip. After 4h of incubation in a 37°C incubator, more L-WRN conditioned media + 10 µM Y27632 (up to 1 mL/well) were added for the first 24h culture. After 24h, cells were subsequently grown on L-WRN conditioned media (without Y27632). Culture media were replaced every 24-48 hours with fresh L-WRN conditioned media and the organoid-derived monolayers were cultured for 10 days before immunostaining or live imaging.

For myosin-IIA-KO induction, Cre recombinase was induced in organoid-derived monolayers with 100nM of 4-hydroxytamoxifen (#SML1666, Sigma-Aldrich) for 24h.

### Preparation of L-WRN conditioned medium

The L-WRN conditioned media was prepared from L-WRN cells, acquired from ATCC (ATCC CRL-3276), according to Miyoshi and Stappenbeck ^68^. L-WRN cells were cultured in L-Cell medium (DMEM high glucose (Sigma-Aldrich, cat. no. D6429) supplemented with FBS (10% v/v), Glutamax (2 mM) and penicillin/streptomycin (100 units/mL). After the first day, selection media was added, containing Geneticin (500 µg/mL) and hygromycin (500 µg/mL). Once confluent cells were passed into five T175 flask and cultured in L-cells media until confluency. Then cells were cultured in Primary Cells Medium (PCM) (Advanced DMEM/F12 supplemented with FBS (20% v/v), Glutamax (2 mM) and penicillin/streptmycin (100 units/mL). The PCM supplemented with Wnt-3a, R-spondin and Noggin secreted by the L-WRN cells was collected every 24h and mixed with freshly made PCM at a 1:1 ratio and was vacuum filtered through 0.22 µm membrane membrane to make EM.

### Human biopsies and preparation of tissue samples

Tissue samples of human duodenum were provided by Necker-Enfants Malades Hospital (Paris, France) and were collected from the Necker Paediatric Anatomo-Pathology Department for retrospective analyses. The biopsies analyzed here comprised 1 patient with gastralgia, 1 patient with anemia, and 1 patient with coeliac disease under gluten-free diet. Duodenal biopsies were collected during endoscopic procedures for diagnosis and/or monitoring of patients. All parents signed informed consent forms approved by the local ethics committee for biopsy exploitation (Unité de Recherche Clinique (URC) of Necker Hospital, URC). For immunohistochemical analyses, biopsies were fixed for 2 h in 4% formaldehyde. The samples were then paraffin embedded. 5 μm sections were de-waxed in a xylene bath, rehydrated in isopropanol and in solutions with decreasing ethanol concentrations, and were processed for immunostaining. De-waxed tissue sections were blocked in 1.5% donkey serum (Sigma-Aldrich, St Louis, Missouri, USA) for 1 h. Primary antibody incubations were performed at 4°C overnight and secondary antibody incubations at room temperature for 2 h, both in 1.5% donkey serum solution. Hoechst 33342 staining (Life Technologies, Paisley, UK) was used to detect nuclei. Tissue sections were mounted in home-made Mowiol 488 solution.

### Caco2 cell culture

Caco2 cells, originally acquired from ATCC, were kindly provided by Dr. S. Robine (Curie Institute, Paris). Caco2 cells were routinely grown in DMEM 4.5 g/l glucose supplemented with 20% fetal bovine serum and 1% penicillin-streptomycin (Gibco, Thermo Fischer Scientific, Waltham, MA, USA) for a maximum of 9 passages. The culture medium was renewed every 2-days.

### Cross-linked Matrigel substrates (CL-Matrigel)

To produce cross-linked Matrigel (CL-Matrigel) substrates, a fresh-made cross-linker solution was prepared by mixing 100 mM NHS (Sigma-Aldrich) and 400 mM EDC (Sigma-Aldrich) in cold PBS 4°C. Glass coverslips (Ø = 18 mm) were plasma treated and then were cooled in a fridge (-20°C, 3 min). Cross-linker solution then was mixed well with thawed pure Matrigel at a ratio of 1:10 (v/v) and 50µL drops of the mixture were then poured on top of the cooled plasma treated coverslip and were spread by tilting the coverslips. Subsequently, the coverslips were placed in a 37°C incubator in a 12 well plate for 2h to form CL-Matrigel layers. While incubation, PBS were poured in 2 empty wells of the 12 well plate ware to prevent gel dehydration. The CL-Matrigel substrates were washed with PBS once and incubated in PBS at 37°C for 24 h to remove unreacted EDC and NHS. The substrates can be stored up to a week in incubator 1X PBS for future utilization. Before use, the coverslips were washed 2 times with 1X PBS.

### Glass coverslip micropatterning with deep UV and cell seeding

The micropatterning protocol was adapted from ^69^. For polystyrene coating, 20x20 glass coverslips (1304369, Schott) were cleaned for 10min in acetone then for 10min in isopropanol in a bath sonicator and then dried with compressed-clean air under a laminar flow hood. They were first coated with adhesion promoter Ti-Prime (MicroChemicals) using a spin-coater (WS-650m2-23NPPB, Laurell) at 3000 rpm for 30s and baked on top heater for 2min at 120°C. Then a 1% polystyrene (MW 260,000, 178891000, Acros Organic) solution in toluene (179418, Sigma-Aldrich) was spin-coated on the coverslip at 1000 rpm.

For coverslip passivation, polystyrene layer was oxidized by exposure to air-plasma as described above and immersed into a solution of poly(L-lysine)-graft-poly(ethylene glycol) (PLL-g-PEG) (ZZ241PO22, JenKem Technology, Beijing) at 0.1 mg/mL in HEPES (10mM, pH 7.4) for 1 hour at room temperature. Coverslips were then washed in HEPES buffer and air dried.

For coverslip micropatterning, passivated coverslips were put in tight contact with a chromed etched photomask (Toppan Photomask). Tight contact was maintained using a homemade vacuum holder. The PLL-PEG layer was burned with deep UV (λ=190nm) through the etched windows of the photomask, using UVO cleaner (Model No. 342A-220, Jelight), at a distance of 1cm from the UV lamp with a power of 6mW/cm2, for 3 min. Exposed coverslips were then incubated with a solution of 10µg/ml fibronectin (F1141, Sigma) in carbonate buffer (100 mM NaHCO_3_ buffer, pH 8.5) for 30min at room temperature. Micropatterned coverslips were then washed with the carbonate buffer. Caco2 cells were directly seeded on these micropatterned coverslips on a 35 mm petri dish and were grown for 4 – 5 days.

### Immunostaining

Routinely, organoid-derived monolayers were fixed using 4% paraformaldehyde for 30 min, then permeabilized using 0.1% triton-x-100 in PBS for 10-15 min. The blocking step was performed in 4% goat serum /1% BSA solution for 1 hour, before proceeding to incubation with primary antibody at 4°C overnight. The next day, the primary antibody was removed and the monolayers were washed 3 times in PBS for 10 min each, before adding the secondary antibody and left to incubate for 2h at room temperature. Finally, monolayers were washed 3 times again for 10 minutes before incubating in Hoechst 33342 for 15 min to stain nuclei. Immunostained samples were mounted in homemade Mowiol solution. For F-actin staining, fluorescently labelled phalloidin was added during secondary antibody incubation.

For immunostaining of *in vivo* mouse intestine, 0.5-mm pieces of mouse jejunum were fixed in 4% PFA at 4°C overnight under shaking for 1 hour. After PBS washes, tissue permeabilization was performed in 1% Triton X-100 / PBS solution for 1 hour at RT with rotation. After a few PBS washes, incubations with primary and subsequent secondary antibodies were done in 0.1% BSA / 0.3 % goat serum / 0.2 % Triton X-100 / PBS overnight at 4°C with rotation. F-actin was stained using phalloidin (Life Technologies Ltd. (Paisley, UK) A12379 - A12380) like a secondary antibody. Hoechst33342 staining was used was used for 30 minutes to detect nuclei followed by PBS washes and a final H2O wash. Immunostained samples were mounted in Vectashield (Vector Laboratories, Burlingame, CA).

### Antibodies and reagents

Mouse monoclonal antibody directed against beta-catenin (clone 14, #610154, IF dilution, 1:100) was from BD Biosciences. Rabbit polyclonal antibodies directed against paxillin (#ab32084, IF dilution 1:100), alpha-1 catenin (#ab51032, IF dilution 1:100), MUC2 (ab#272692, IF dilution 1:200) and alpha-tubulin (#ab18251, IF dilution 1:100) were from Abcam. Mouse monoclonal directed against cytokeratin-20 was from Agilent Dako (#M7019, IF dilution 1:200). Mouse monoclonal directed against alpha-1 actinin (#TA500072S) was from Origene. Rabbit monoclonal antibody directed against E-cadherin (clone 24E10, #3195S, IF dilution 1:100), rabbit polyclonal antibodies directed against P-MLC2 (#3674, IF dilution, 1:100, and #95777S, IF dilution, 1:100) were from Cell Signalling Technology. Rabbit polyclonal antibody directed against non-muscle myosin heavy chain II-A antibody (clone poly19098, #909801, WB dilution 1:500) was from Biolegend. Rabbit polyclonal directed against Par3 (#07-330) was from Millipore. Rabbit polyclonal directed against ZO-1 (#R26.4C) was from DHSB. Rabbit polyclonal directed against tricellulin (#48-8400) was from Thermo Fisher Scientific. Rabbit polyclonal directed against ezrin was a gift from Dominique Lallemand (Institut Cochin, Paris). Goat anti-mouse-Alexa-488, 568, anti-rabbit-Alexa488, 568 or 647 were from Life Technologies (Paisley, UK). Alexa Fluor-488 and -568 phalloidin were from Thermo Fisher Scientific. Nuclei were stained with Hoechst 33342 solution incubation (Life Technologies) at a 1:1000 dilution. CellMask Actin Deep Red actin tracking stain (#A57245) was from Invitrogen (Thermo Fisher Scientific). Azidoblebbistatin (#MPH-198) was from MotorPharma (Budapest, Hungary). Blebbistatin, and Y-27632 were from Sigma Aldrich (Saint-Louis, MO, USA).

### Drug/EDTA treatments and live imaging

Drug treatments were done using the following conditions: 10μM blebbistatin for 1.5h, 50nM calyculin A for 1h, or 3µM nocadazole for 3h. Organoid-derived monolayers were incubated in drug containing medium, then washed out with PBS and prepared for immunostaining or live-imaging. Cells were incubated for 1.5h in DMSO as controls. For live blebbistatin washout experiments, organoid monolayer was incubated in 10 µm blebbistatin in culture media for 1h, then was washed 2 times with warm PBS and then was immediately imaged live in blebbistatin free culture media. For live blebbistatin photo-activation, organoid derived monolayers were first stained with CellMask actin-GFP and then were incubated with 10μM azidoblebbistatin (Cat#DR-A-081, Motor pharma) or DMSO controls for 1h. The samples were then taken for live imaging in a LSM 980 scanning probe microscope equipped with a multiphoton setup. Azidoblebbistatin was photoactivated in the selected ROI using a 2-photon 800 nm laser irradiation ^70^ and live images were acquired. For all the live imaging monolayers were stained with live probes like CellMask actin-GFP (F-actin labelling) and Hoechst 33342 (nucleus labelling).

For chelation of extracellular calcium, CellMask actin-GFP-stained organoid-derived monolayers were treated with 0.5mM EDTA in the culture media for 40min in live imaging using a LSM 880 scanning probe microscope.

### Laser ablation experiments

For Laser ablation experiments, F-actin was first labelled in live organoid-derived monolayers by incubating with CellMask™ Green Actin Tracking stain (Invitrogen) for 1 hour in culture media. Labelled monolayer was then taken in a live imaging setup on a confocal microscope equipped with laser ablation setup. The ablation was done using either a spinning disc microscope (CSU XI) with a pulsed 355nm UV laser (iLas system, Roper Scientific) at 25% power in Metamorph, or a LSM 880 microscope with a tunable two-photon laser set to 710 nm at 20-25% power for 20-30 iterations, based on a previously described protocol ^71^. Node displacement was manually tracked using TrackMate in imageJ.

### Live imaging

Dynamics experiments on live organoid-derived monolayers were performed on a live imaging setup on an inverted Zeiss microscope equipped with a CSU-X1 spinning disk head (Yokogawa – Andor), using Zeiss 40X and 63X objectives.

### Segmentation and analyses

#### Segmentation

We used CellPose 2.0 (https://www.cellpose.org/; ^72^) to perform cell segmentation. Mask dilatation was used to extract the cell contours and location of tricellular junctions, called vertices.

#### Hexatic order analysis

For each cell (indexed by *J* = 1,2,3, … , *N* with *N* the total number of cells), we define a hexatic order parameter value:

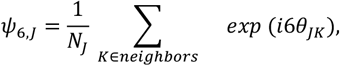

where *θ*_*JK*_= *arg* (*r*_*K*_− *r*_*J*_), is the angle between the considered cell center, denoted *r*_*J*_, and the one of its neighbour, *r*_*K*_; *N*_*J*_ is the number of neighbor cells of the *J*-th cell. The overall hexatic order is then defined as 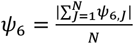. We then estimated the normalized spatial correlation function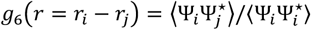, where Ψ_*i*_= (*ψ*_6_)_*i*_.

#### Triangular order

We define a triangular order parameter value:

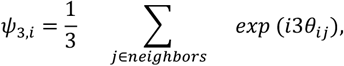

where *θ*_*ij*_ = *arg* (*r*_*j*_ − *r*_*i*_), is the angle of the cell-cell junction *ij*. The overall triangular order is then defined as 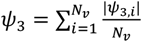.

#### Strain

For each detected cells, we estimate a cell shape tensor *Λ*whose *xx* components read:

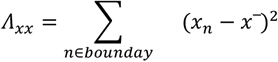

where (*x*_*n*_, *y*_*n*_) is the coordinate of the boundary pixels *n* and 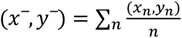 is the cell barycenter; the *xy* and *yy* components are defined similarly. The matrix *Λ* has positive eigenvalues *Λ*_1_ and *Λ*_2=_ with *Λ*_1_ ≥ *Λ*_2_. We then define the cellular strain as:

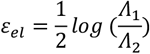

In the strain quantification, we exclude detected objects with an area smaller than 2 microns.

#### Tracking

We used StarDist in TrackMate (FiJi ^73^) for cell tracking analyses. We used a Kalman tracker parameterized with a search radius of approximately half the cell typical size, an initial search radius 10% higher, and a maximum frame gap of 2 ^74,75^.

#### Velocity Correlation Functions

For every cell, with position *r*_*i*_, we compute the velocity autocorrelation function:

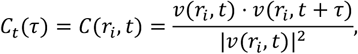

where *τ* is the delay time. We also evaluate the spatial velocity correlation function:

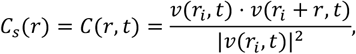

where *r* = *r*_*j*_ − *r*_*i*_ and *r*_*i*_ is the position of the cell *j*. For each experiment, we estimate the average correlation function ⟨*C*_*s*_ (*r*)⟩ over all tracks and all times (see Figure 7E-J in the main text). Practically, in our evaluation of the velocity functions, we discarded every track which was shorter than 4 frames. The spatial correlation length (*λ*) was then obtained using the nonlinear least-squares fit (nlinfit) of Matlab R2023a and the fitting function y = exp(-x/*λ*).

### Computational model

We employed a cell-based computational model, called the vertex model ^76,77,78^, to simulate the multicellular response of cell differentiation, cell ablation, and contractility recovery of multicellular actin star network. In this model, the cell monolayer is represented as a tiling of polygons, see Figure 6D. The dynamics of cells are determined by force balance equations at each vertex:

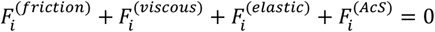

where (1) 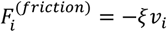 is the friction force between the monolayer and the substrate, with *ξ* being the friction coefficient and 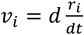 being the velocity of the vertex *i*. (2) 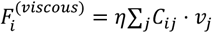 is the viscous force at vertex *i*, which depends on the velocities of the neighbor vertices *j* and scales proportionally to a viscous modulus *η* that models dissipation along the cell-cell junction and within the bulk cytoplasm; *C*_*ij*_ is a viscous structure tensor that depends on the vertices’ positions (*r*_*i*_), and the topological relation of cells and vertices ^79^ (see Supplementary Information); (3) 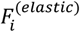 is the elastic force stemming from variations in the cell shape, classically expressed as 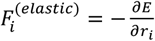 with ^77,80,79,81,82^ :

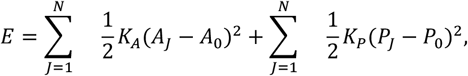

where *K*_*A*_ and *K*_P_ are the rigidities associated with cell area and cell perimeter; *A*_0_ and *P*_0_ are the preferred cell area and the preferred cell perimeter, respectively; *A*_*J*_ and *P*_*J*_ are the actual area and perimeter of the *J*-th cell, respectively. (4) the force 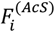 corresponds to the forces induced by the multicellular actin star network; we decompose this force into two contributions, resulting from: (4.1) an extra pulling stress 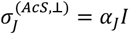 with *α*_*J*_ ≥ 0 quantifies the intensity of the contractility of the actin star network within the *J*-th cell, see Supplementary Information, Figure S1; and (4.2) an AcS-induced tension *χ*_*J*_ > 0 parallel to edges of the *J*-th cell, see Supplementary Information, Figure S1. We show that the AcS-induced pulling stress *α*_*J*_ and the AcS-induced tension *χ*_*J*_ are equivalent to renormalizing *E* into *E*_*eff*_ with a renormalized target area 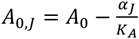 and a renormalized target perimeter 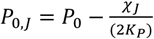 (see Supplementary Materials). Consequently, we obtain the following dynamic equation on the velocities:

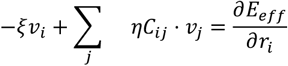

We then solve the latter equation to estimate the vertex displacement at each simulation time step. Last, we mention that in the laser ablation simulations, the cells at the *k*-th row are defined by the distance *d* of the cell centre to the ablation site if it satisfies *k*-1/2 < *d*/*L*_cell_ < *k*+1/2 with *L*_cell_ being the cell size.

### Statistical analysis

All statistical analyses were performed using Prism (GraphPad Software, San Diego, CA, USA, version 9.0). Statistical details of experiments can be found in the figure legends. Unless otherwise stated, experiments were replicated 3 times independently.

