## Supplementary Figures and Supplementary Information for "A multicellular actin star network underpins epithelial organization and connectivity"

**Barai et al.**

**SUPPLEMENTARY INFORMATION FILE**

**Supplementary Figures 1- 16**

**Supplementary Information: File detailing the vertex model simulation procedure are in the online version of the paper.**

#### SUPPLEMENTARY FIGURES

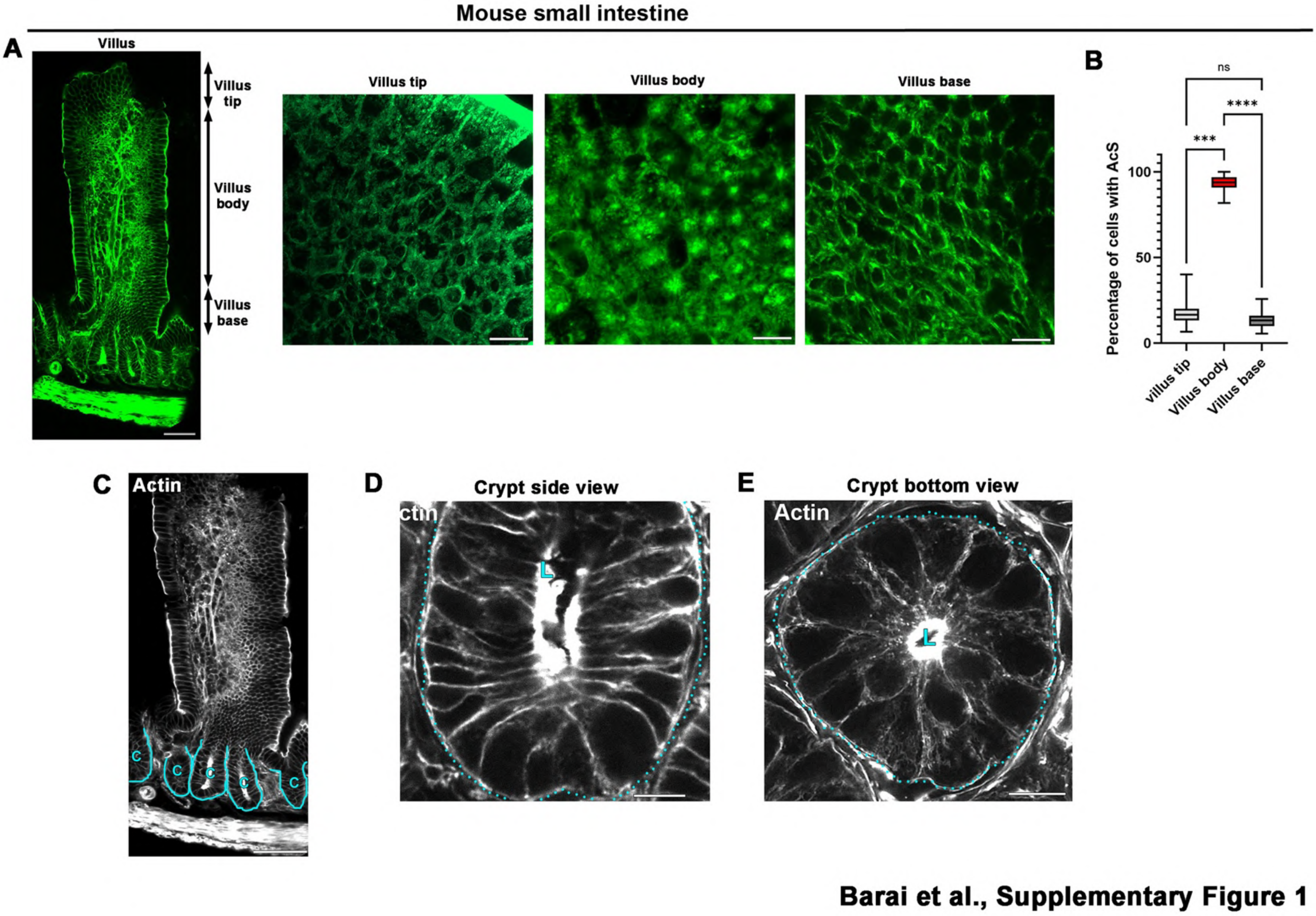


**Supplementary Figure 1: (A)** Confocal analysis of basal actin distribution in the mouse villus compartment. Villus base, body and tip views are shown on the right. Scale bar, left panel 65 μm, right panels 10 μm. **(B)** Statistical analysis of the proportion of cells containing AcS at the base, body and tip parts of the villus. Percentage of AcS-positive cells at the villus tip = 16.67% (13.33-20.00) (median (IQR)), at the villus body =93.64% (90.80-96.84), at the villus base = 13.33% (10.00-16.08). N (villus tip) = 15, N (villus body) = 16, N (villus base) = 16, n (cells at villus tip) = 435, n (cells at villus body) = 525, n (cells at villus base) = 425. Kruskal Wallis Test, Dunn's multiple comparison test, ***p=0.0001, ****p<0.0001. **(C-E)** Confocal analysis of actin distribution in the mouse crypt compartment. Crypt side view (D) and bottom view (E) are shown. Crypt domains are delimited with blue line. L, crypt lumen. Scale bar C 100 μm, D-E 10 μm.

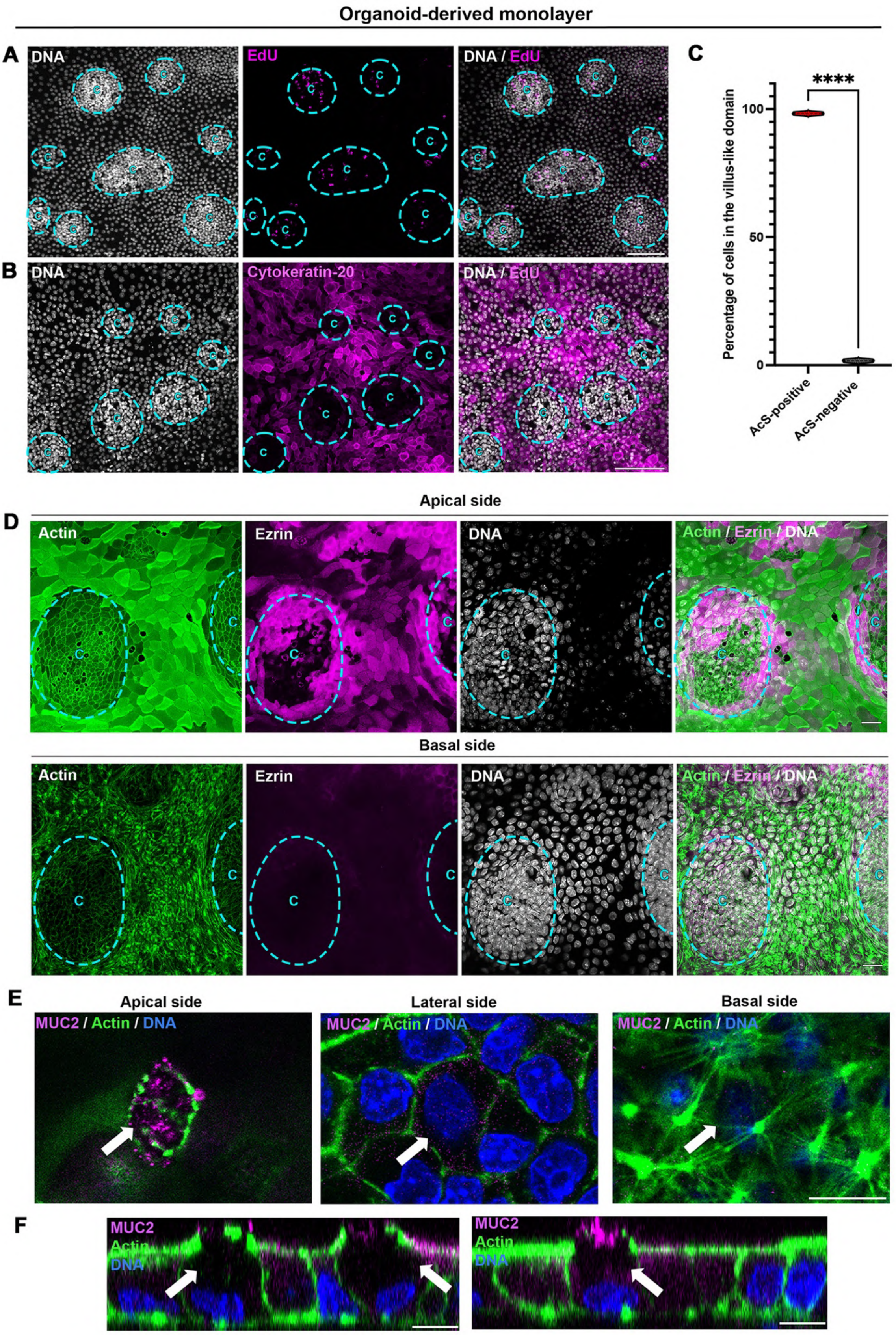


**Barai et al., Supplementary Figure 2**

**Supplementary Figure 2: (A-B)** Confocal analysis of EdU (A) and cytokeratin-20 (B) distribution in organoid-derived monolayers. Nuclei are shown in gray. Crypt-like domains are delimited in blue. C, crypt-like domain. Scale bar, 100 μm. **(C)** Statistical analysis of the proportion of AcS-positive or -negative cells in the villus-like domain of organoid-derived monolayers. Mean percentage of AcS-positive cells = 98.26±0.33% (mean±S.E.M), mean percentage of AcS-negative cells = 1.737±0.33% N = 3 experiments, n = 331 cells. Unpaired t-test, ****$p<0.0001$. **(D)** Confocal analysis of actin (green), ezrin (magenta) and DNA (gray) in the apical or basal side of organoid-derived monolayers. Crypt-like domains are delimited in blue. C, crypt-like domain. Scale bar, 20 μm. **(E-F)** Confocal analysis of MUC2 (magenta) and actin (green) in organoid-derived monolayers. xy (E) and xz (F) views are presented. Scale bar, 10 μm.

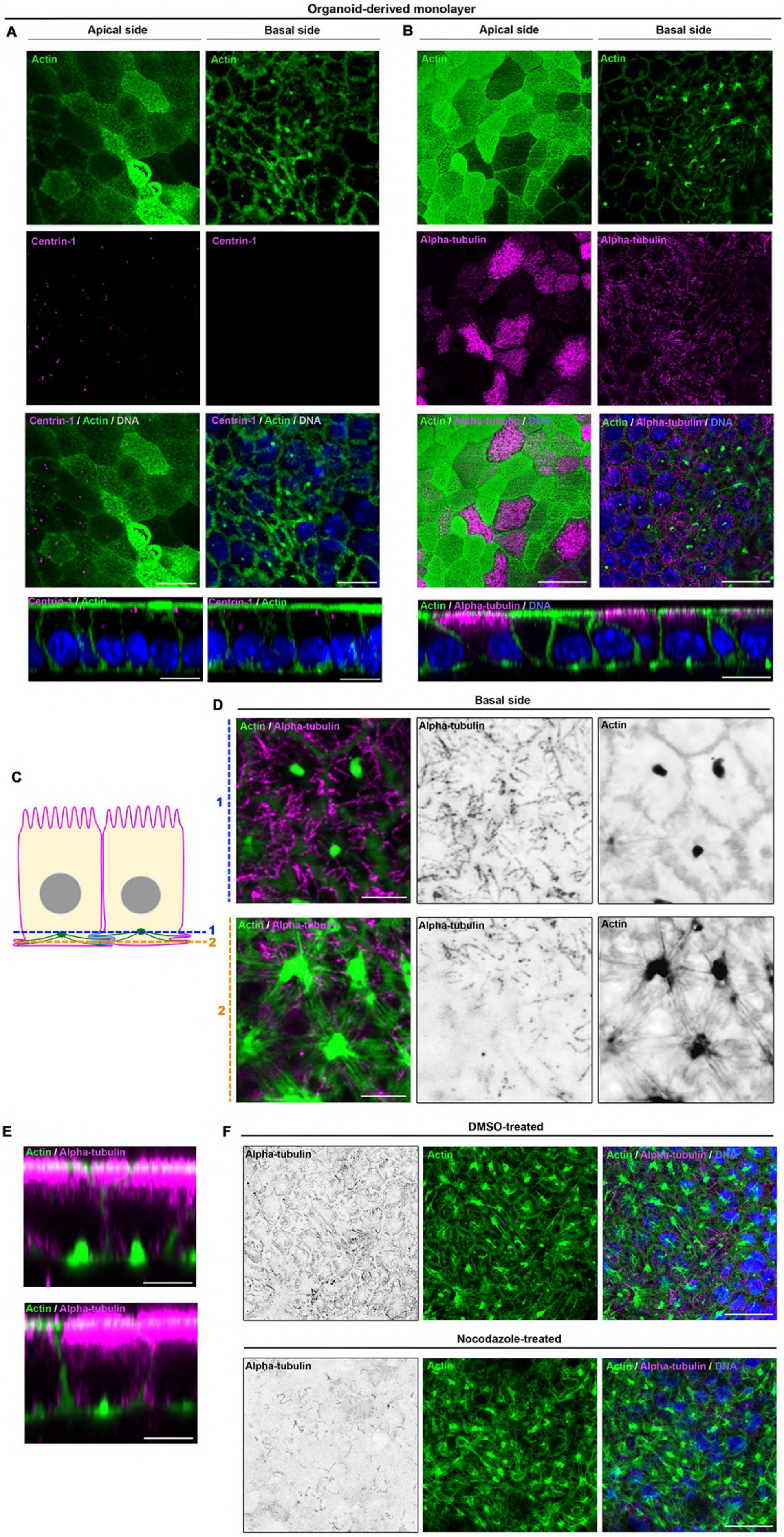


Barai et al., Supplementary Figure 3

**Supplementary Figure 3: (A)** Confocal analysis of the distribution of actin (green), centrin-1-GFP (magenta) and H2B-mCherry (blue) in the apical and basal sides of the villus-like domain in organoid-derived monolayers. Representative xz views are presented in the lower panel. Scale bar, xy views 10 µm, xz views 10 µm. **(B)** Confocal analysis of the distribution of actin (green), alpha-tubulin (magenta) and DNA (blue) in the apical and basal sides of the villus-like domain in organoid-derived monolayers. A representative xz view is presented in the lower panel. Scale bar, xy views 20 µm, xz view 10 µm. **(C)** Scheme showing the cell area where high-magnification imaging was done in (D) in sub-basal area (1, blue dotted line) or basal area of enterocytes (2, orange dotted-line). **(D-E)** High-magnification of actin (green) and alpha-tubulin (magenta) focusing on the basal side of enterocytes in the villus-domain of organoid-derived monolayers. Representative xz views are presented in (E). Scale bar, xy views (D) 5 µm, xz views (E) 5 µm. (**F**) Confocal analysis of the distribution of actin (green), alpha-tubulin (magenta) and DNA (blue) in the basal side of the villus-like domain in organoid-derived monolayers treated with DMSO or nocodazole. Scale bar, 20 µm.

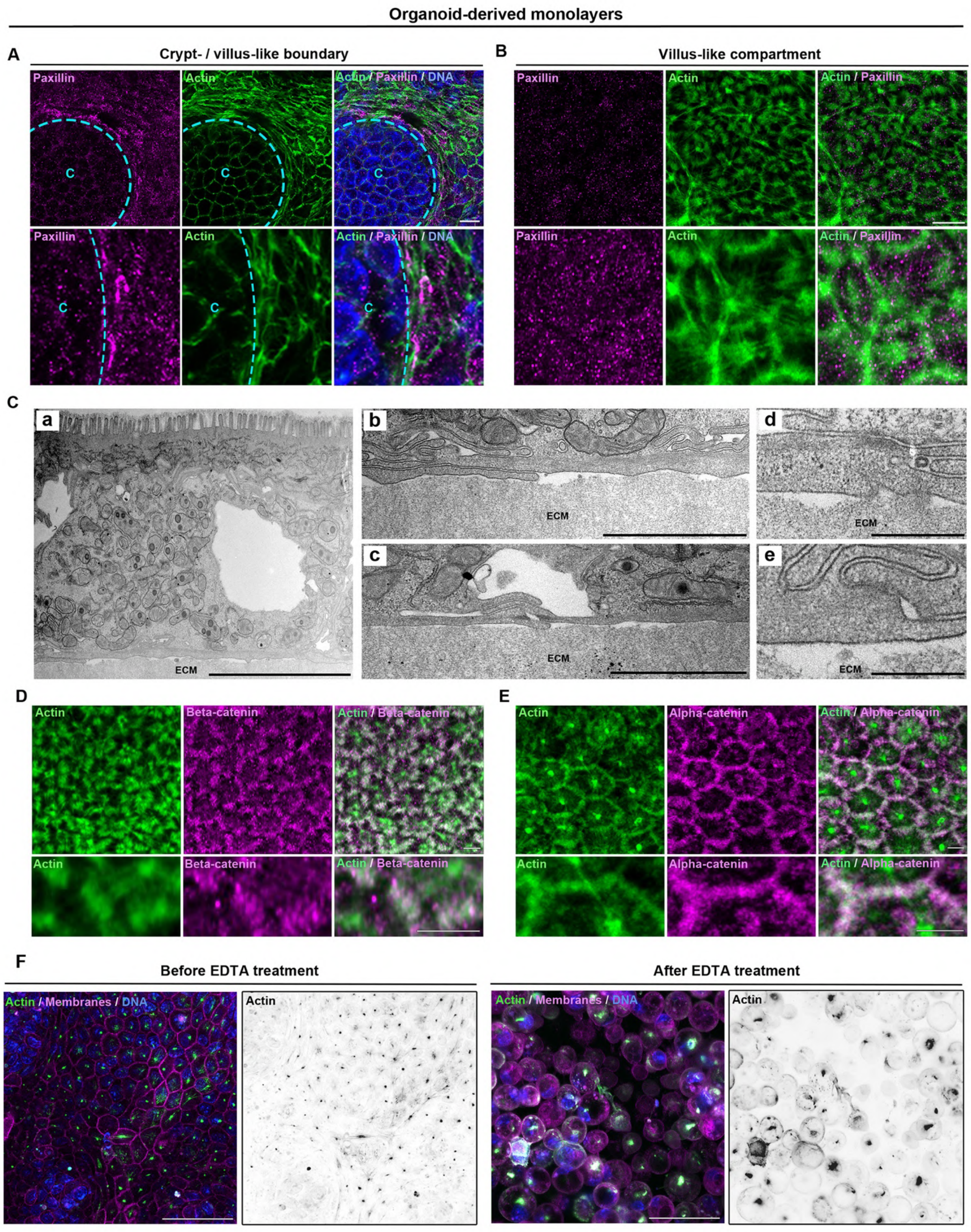


**Barai et al., Supplementary Figure 4**

**Supplementary Figure 4: (A-B)** Airy scan confocal analysis of paxillin (magenta) and actin (green) in the area of crypt- / villus-like boundary or in the differentiated compartment. Crypt-like domains are delimited in blue. C, crypt-like domain. Scale bar, 10 μm. **(Ca-e)** Transmission electron microscopy analysis of the basal domain of differentiated cells in organoid-derived monolayers. ECM, extracellular matrix. Scale bars, (a) 5 μm, (b-c) 2 μm, (d-e) 0.5μm. **(D-E)** Confocal analysis of β-catenin (magenta) or α-catenin (magenta) and actin (green) in the differentiated compartment of organoid-derived monolayers. Scale bar, 5μm. **(F)** Confocal analysis of actin (green), membranes (magenta) and DNA (blue) in DMSO- or EDTA-treated organoid-derived monolayers. Scale bar, 50μm.

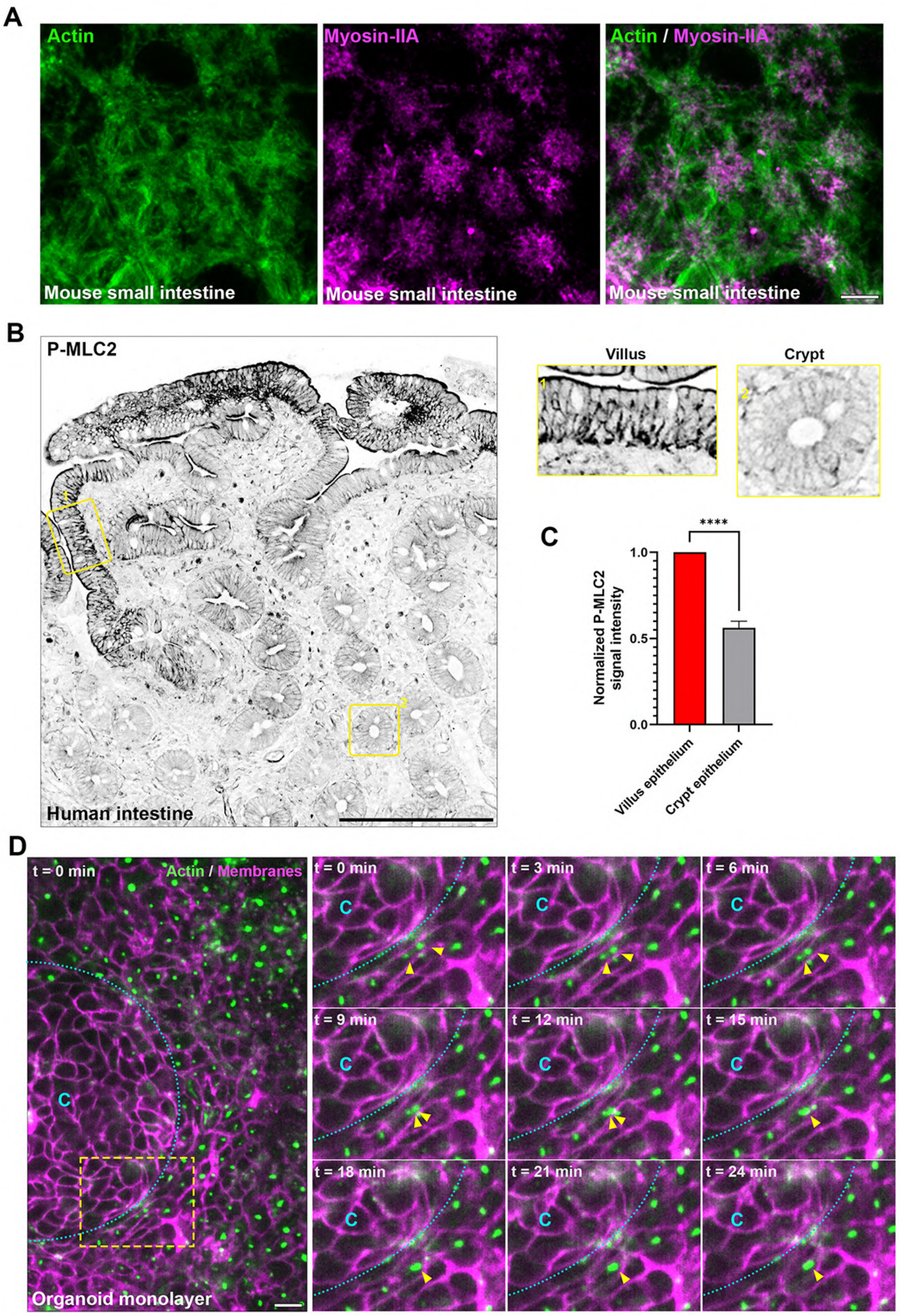


Barai et al., Supplementary Figure 5

**Supplementary Figure 5: (A)** Confocal analysis of actin (green) and myosin-IIA (magenta) in the basal domain of differentiated cells along the villus of the mouse small intestine. Scale bar, 5 µm. **(B)** Confocal analysis of P-MLC2 distribution in the human intestinal tissue. Areas boxed in yellow are presented on the right. Scale bar, 200 µm. **(C)** Statistical analyses of the signal intensity of P-MLC2 in the villus epithelium and the crypt epithelium in the human small intestine. Normalized signal intensity in the villus epithelium = 1 (1-1) (median (IQR)), in the crypt epithelium = 0.5615 (0.5330-0.6008). N (intestinal biopsies) = 3. Mann Whitney, ****$p<0.0001$. **(D)** Time-lapse analysis of AcS formation at exit of the crypt-like domain in organoid-derived monolayers. Actin (green) and membranes (red) are shown. Yellow arrowheads point toward one AcS formation. Crypt-like domains are delimited in blue. C, crypt. Scale bar, 10 µm.

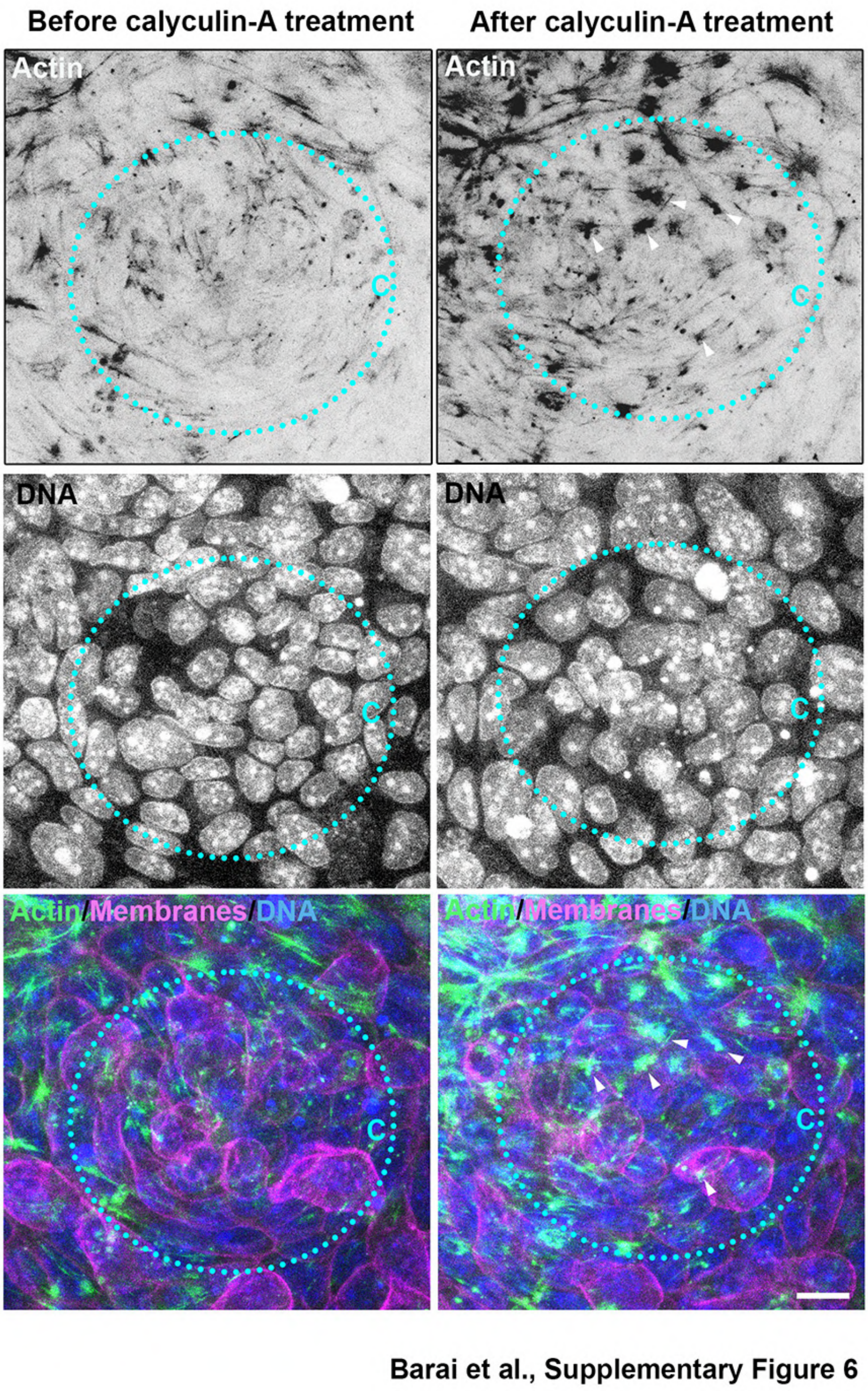


**Supplementary Figure 6:** Time-lapse images of actin (green), membranes (magenta) and DNA (blue) in a crypt-like domain before and after 50nM calyculin-A treatment. Crypt-like domains are delimited in blue. White arrowheads point to AcS-like structures. C, crypt-like domain. Corresponding time-lapse is shown in Video S4. Scale bar, 10 µm.

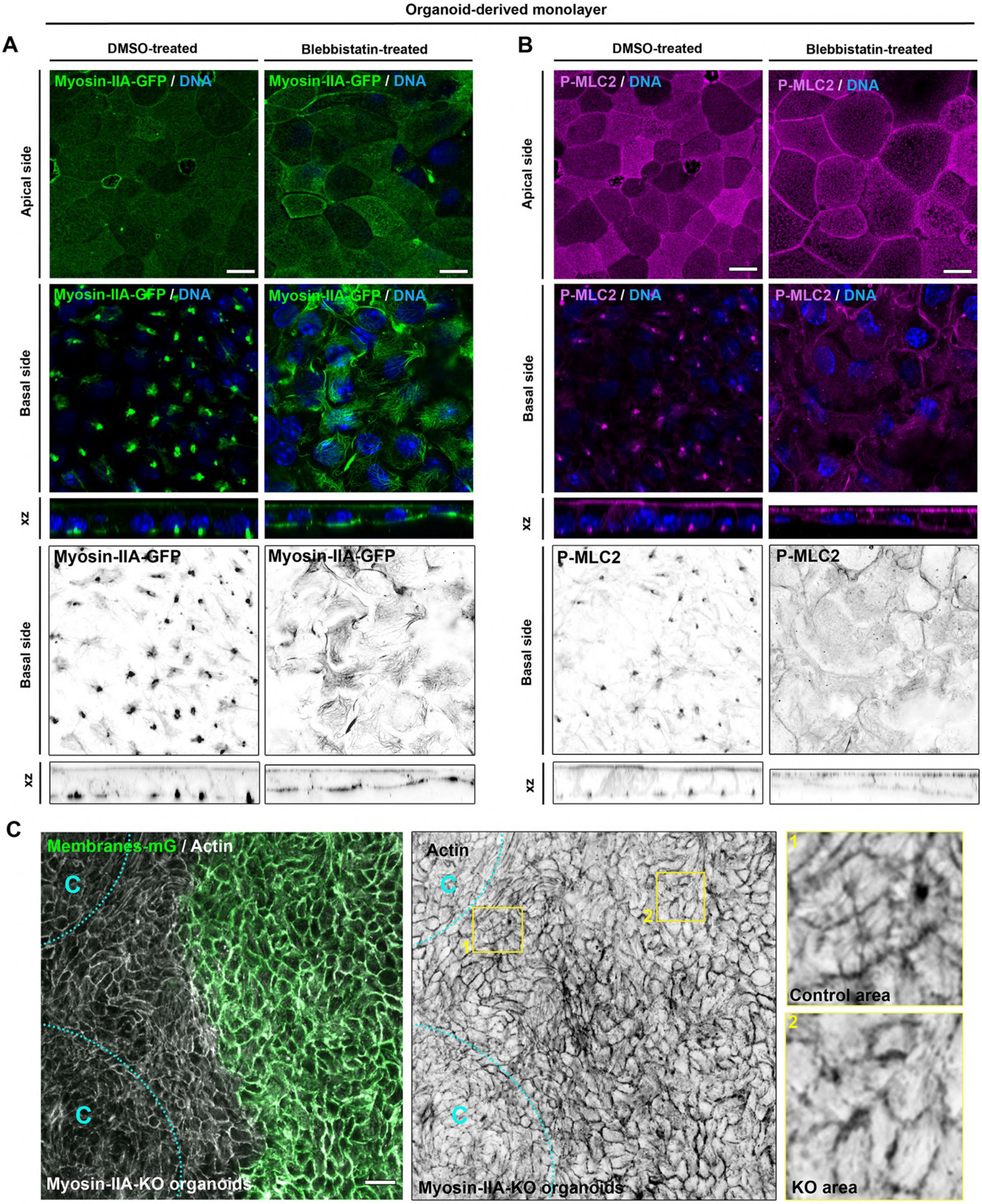


Barai et al., Supplementary Figure 7

**Supplementary Figure 7: (A-B)** Confocal analysis of the distribution of myosin-IIA-GFP (green) or P-MLC2 (magenta) with nuclei (blue) on the apical and basal sides of differentiated cells after treatment of organoid-derived monolayers with DMSO (A) or blebbistatin (B). xz views are also presented. Scale bar, 10 µm. **(C)** Confocal analysis of actin distribution in mosaic myosin-IIA-KO-mT/mG organoid-derived monolayers. Tamoxifen-induced myosin-IIA-KO cells are positive for membranes-mG (green). Crypt-like domains are delimited in blue. C, crypt. Areas boxed in yellow are presented on the right. Scale bar, 20 µm.

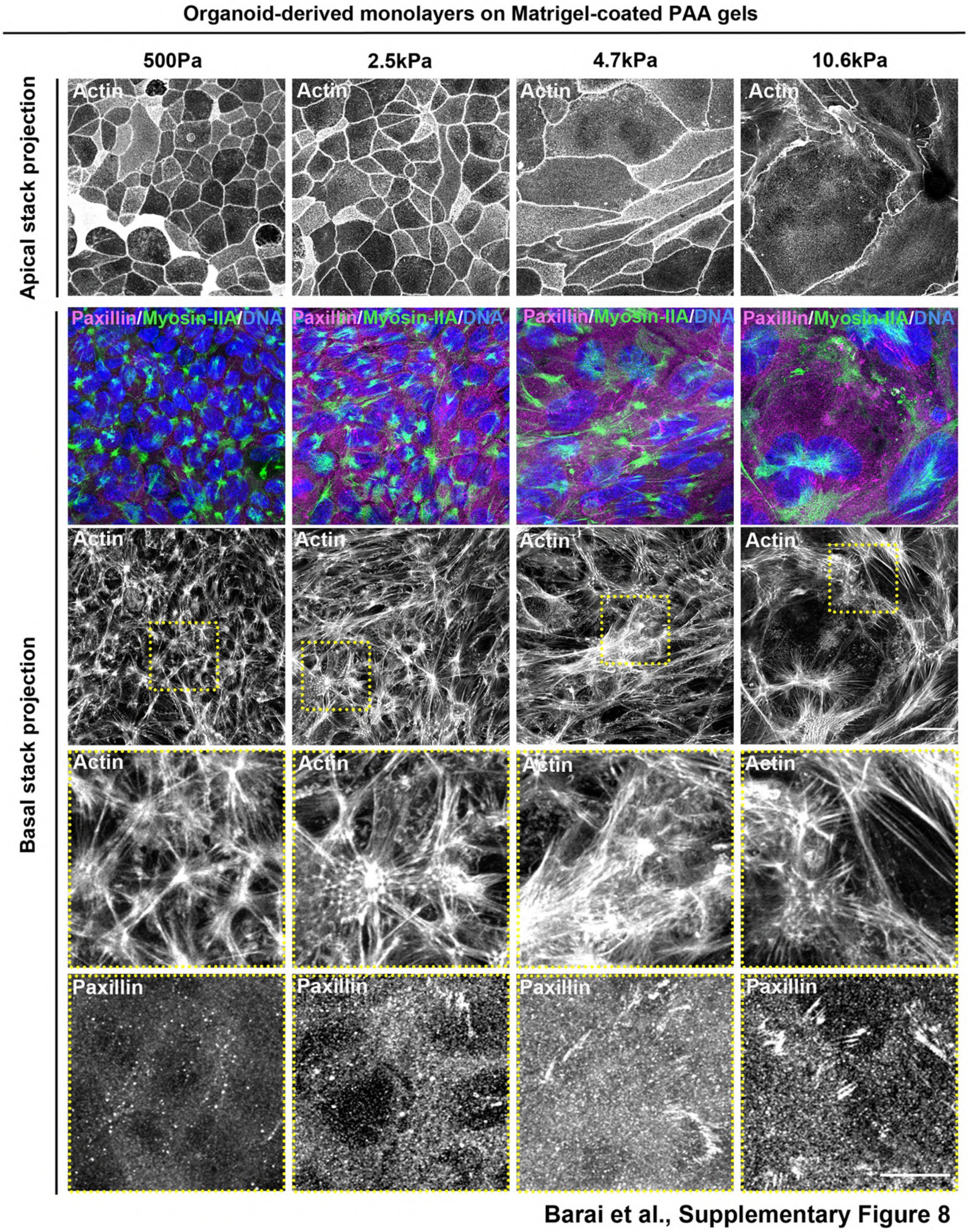


**Supplementary Figure 8:** Confocal analysis of paxillin (magenta)-positive focal adhesions together with the basal actin (green) arrangement in organoid-derived monolayer grown on 300Pa, 2.5, 4.7kPa or 10.6kPa PAA gels. Boxed regions are enlarged in lower panels. Scale bars, 20μm (top) and 10μm (enlarged lower panel).

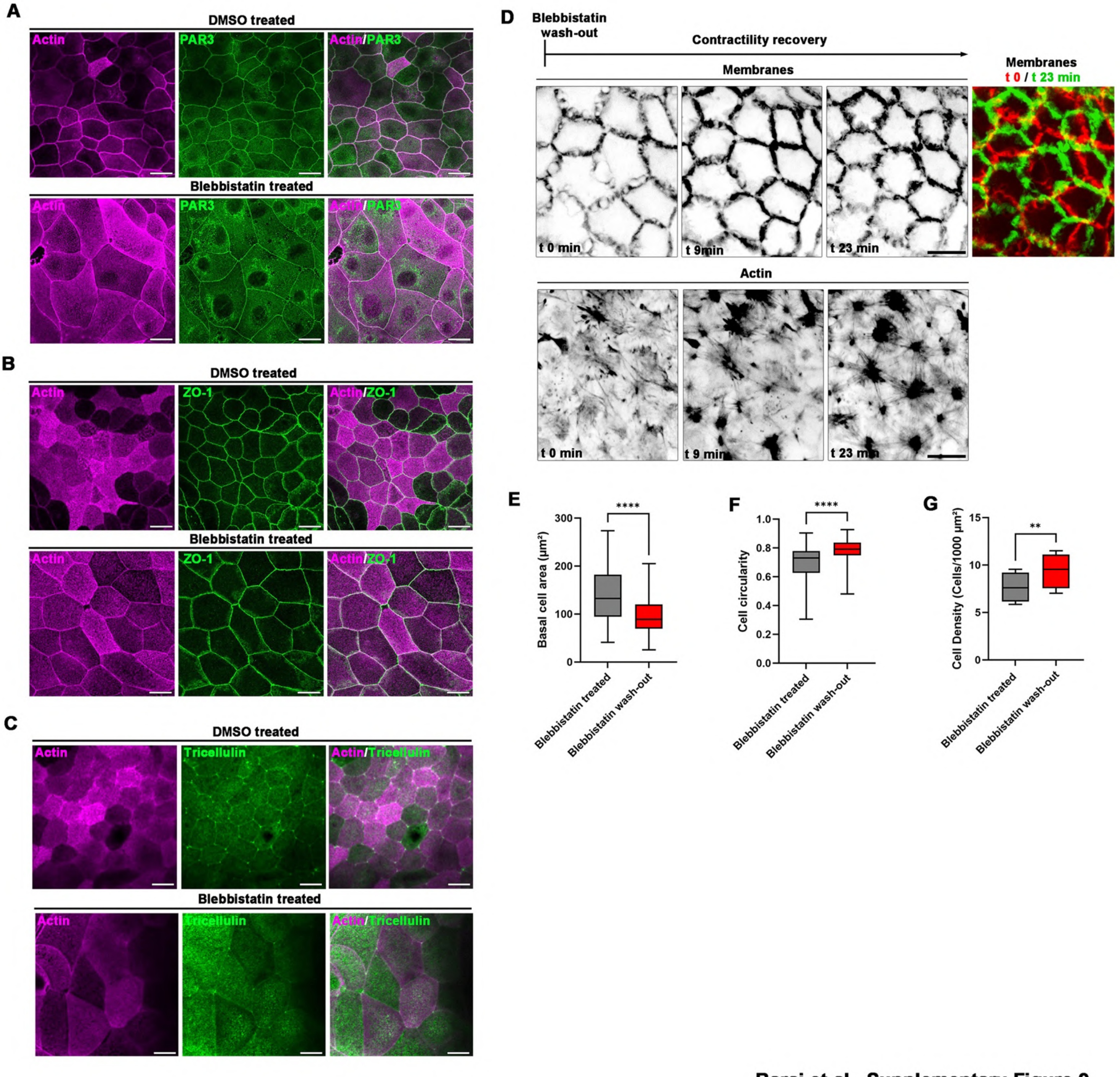


**Supplementary Figure 9: (A-C)** Confocal analysis of the localization of actin (magenta), Par3 (green, **A**), ZO-1 (green, **B**) and tricellulin (green, **C**) in the apical domain of control or blebbistatin-treated organoid-derived monolayers. Scale bar, 10 µm. **(D)** Time-lapse of CellMaskActin and membranes-tdTomato organoid-derived monolayer after 1h blebbistatin treatment and then wash-out (t = 0min). Color-coded t-projection of time-lapse series of membranes-tdTomato signal at the basal domain during contractility recovery in organoid-derived monolayer is shown (t=0min in green, t = 46 min in red). Scale bar, 10 µm. **(E)**

Statistical analysis of basal cell area before and after blebbistatin wash-out. Basal area before = 132.4 (94.45-181.9) (median (IQR)), after = 89.15 (69.62-120.1). N = 4 experiments, n = 100 cells. Wilcoxon test, ****$p<0.0001$. **(F)** Statistical analysis of basal cell circularity before and after blebbistatin wash-out. Basal circularity before = 0.729 (0.626-0.778) (median (IQR)), after = 0.792 (0.749-0.836). n = 100 cells. Wilcoxon test, ****$p<0.0001$. **(G)** Statistical analysis of cell density before and after blebbistatin wash-out. Mean cell density before = 7.65±0.79 (mean±S.E.M), after = 9.40±0.93. Paired t-test, **$p=0.004$.

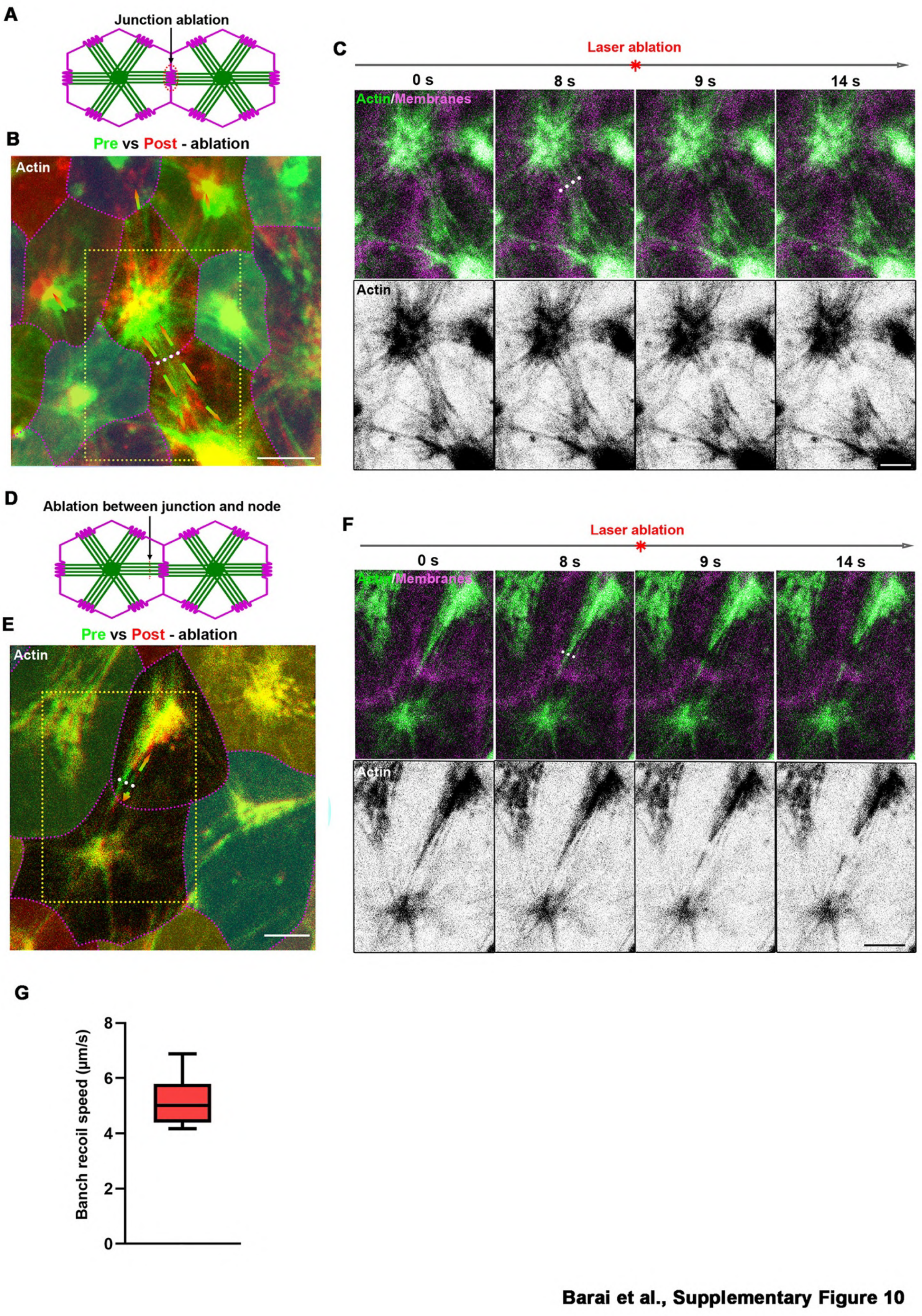


**Supplementary Figure 10:** Time-lapse of CellMaskActin and membranes-tdTomato organoid-derived monolayer after laser ablation of either a cell-cell junction **(A-C)** or a AcS branch close between AcS node and cell-cell junction **(D-F)**. White dotted lines point to the location of laser ablation. B and E show actin before (green) and after (red) ablation along the white dotted line. Arrows indicates the movement of AcS branch and node before and after ablation. B and E are also overplayed with segmented cells with magenta dotted lines indicating cell membranes. C and F show time-lapse images corresponding to the yellow dotted area in B and E, respectively. In C and F, merged images of membranes (magenta) and actin (green) are shown in top panel, and images of inverted grayscale actin on the bottom panel. Laser ablation is done along the white dotted line between 8 s and 9 s. Scale bars, B,E-F, 5 μm; C, 3 μm. Time-lapses corresponding to A-C and D-F are shown in Videos S8 and S9, respectively. **(G)** AcS branch instantaneous ($\Delta t$ = 10 s) recoil speed after laser ablation. Average recoil speed = 5.18 ± 0.88 μm/s (mean±S.D.). n = 9 ablated branches from 3 independent experiments.

**Laser ablation post blebbistatin treatment**

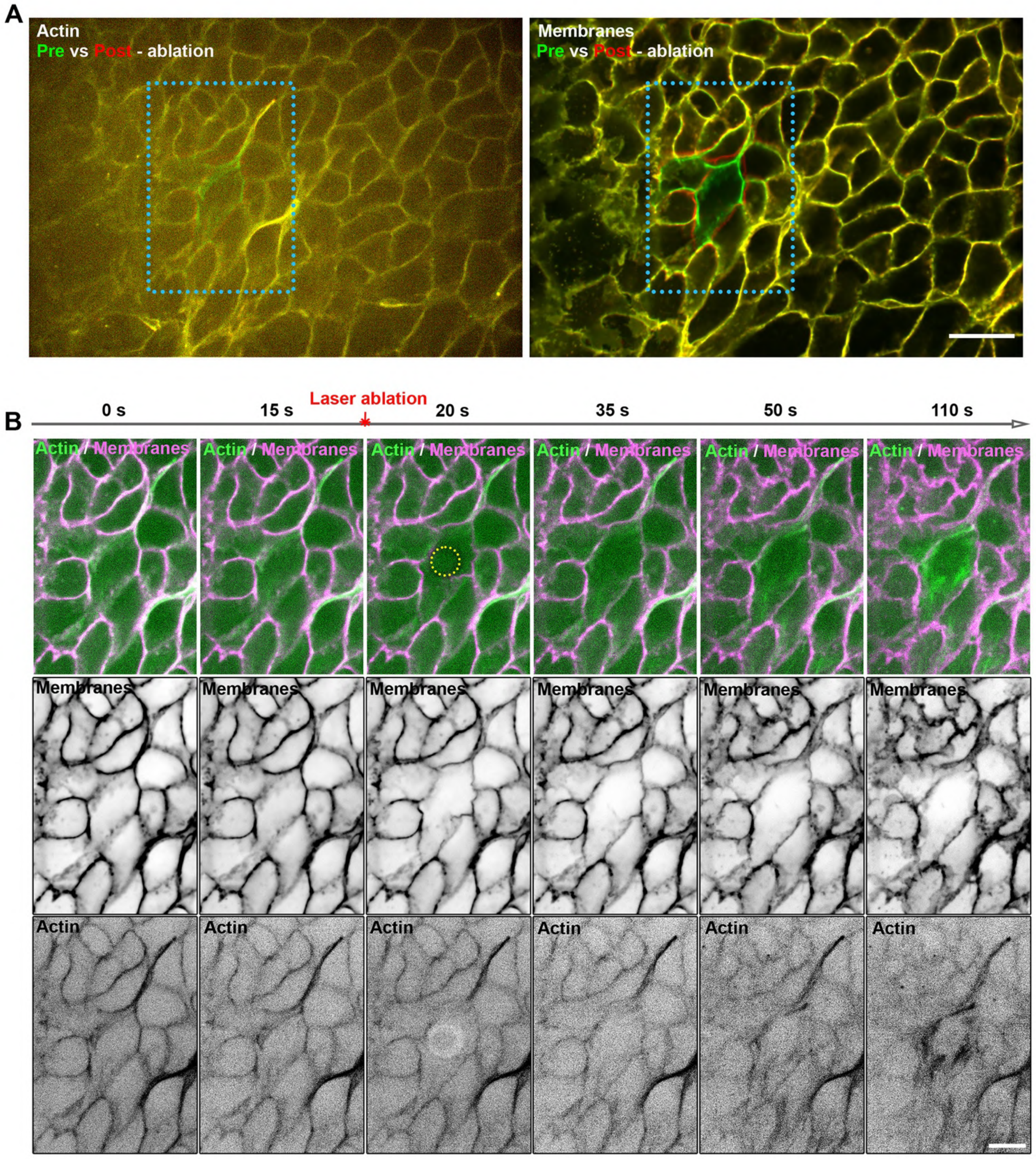


**Barai et al., Supplementary Figure 11**

**Supplementary Figure 11: (A)** Time-lapse of CellMaskActin and membranes-tdTomato organoid-derived monolayer treated for 1h with 10µM blebbistatin, before (green) and after (red) laser ablation of actin (right) and membranes (left) in a given cell**.** Scale bar, 20 µm. **(B)** Enlarged inset showing time-lapse of CellMaskActin (green or inverted grey) and membranes-tdTomato (magenta) organoid-derived monolayer treated for 1h with 10 μM blebbistatin and then laser ablation of actin in a given cell. Laser ablation is carried out along the yellow circle between 8 s and 9 s. Scale bar, 10 μm.

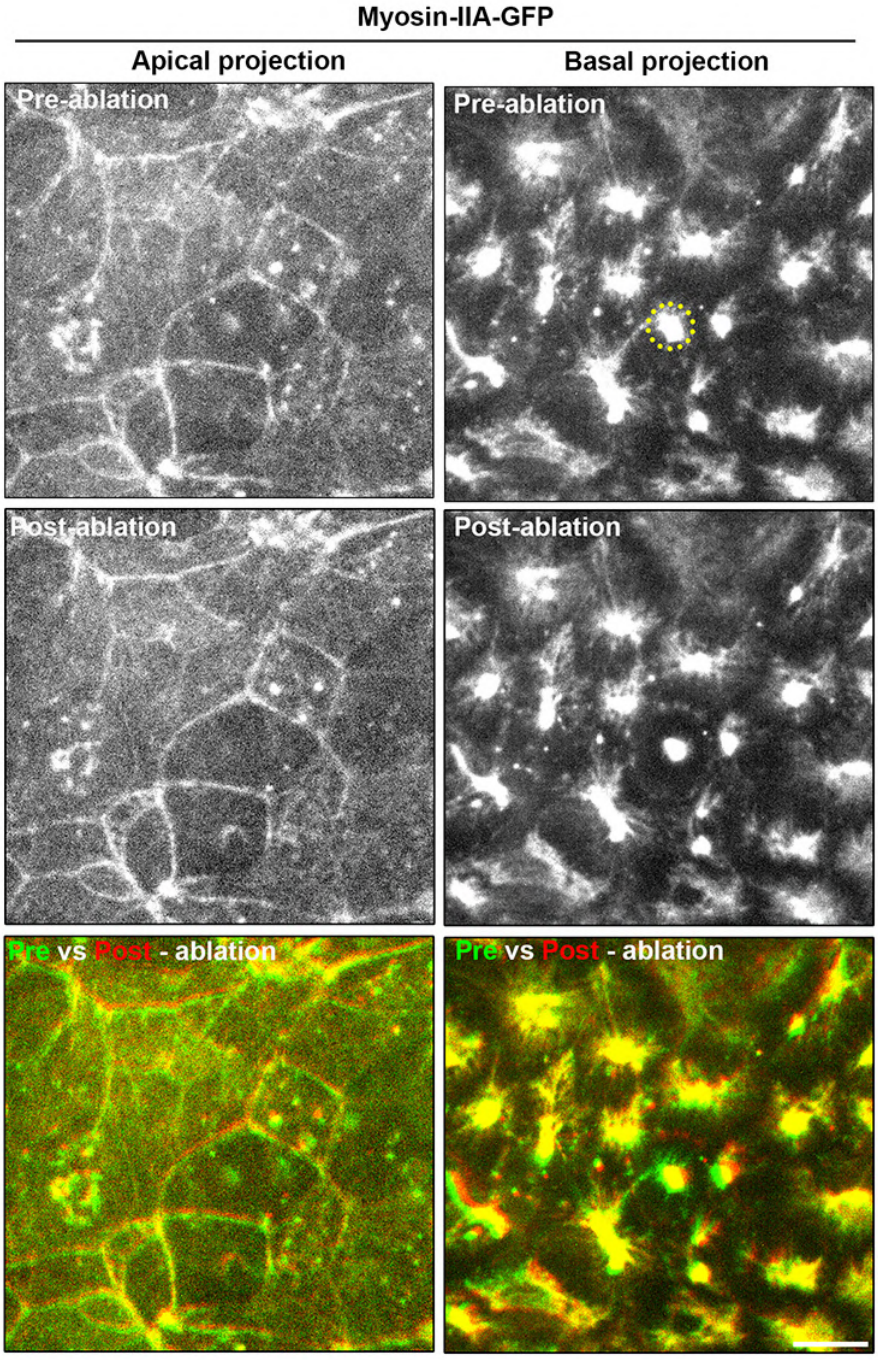


Barai et al., Supplementary Figure 12

**Supplementary Figure 12:** Confocal apical and basal plane projections of myosin-IIA GFP organoid-derived monolayer before and after laser ablation. Yellow dotted circle points to the location of laser ablation. Bottom panel apical and basal myosin-IIA distribution before (green) and after (red) ablation along. Scale bar, 10 μm.

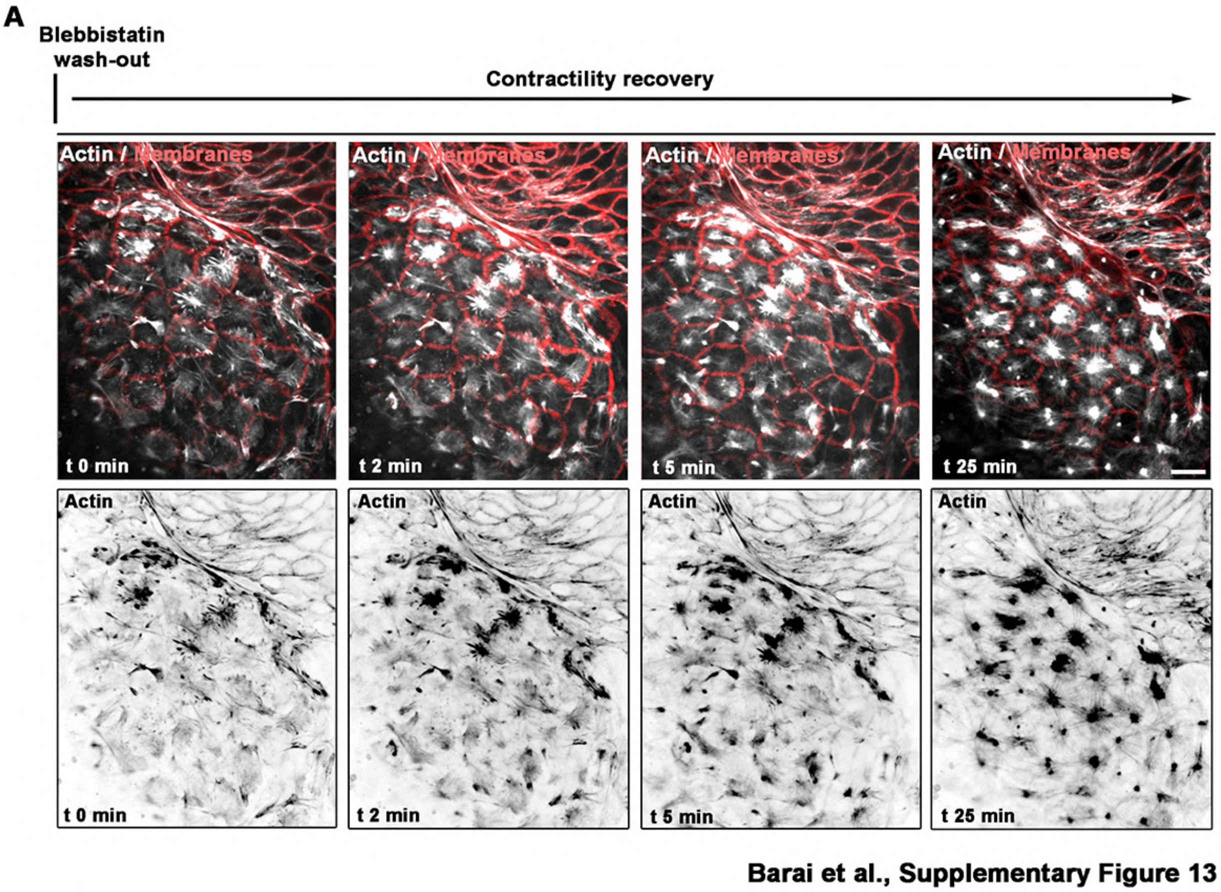


**Supplementary Figure 13:** Time-lapse of CellMaskActin (white) and membranes-tdTomato (red) organoid-derived monolayer after 1h blebbistatin treatment and then wash-out (t = 0 min). Actin-based protrusive structures are observed in the black and white panel. Scale bar, 10 µm.

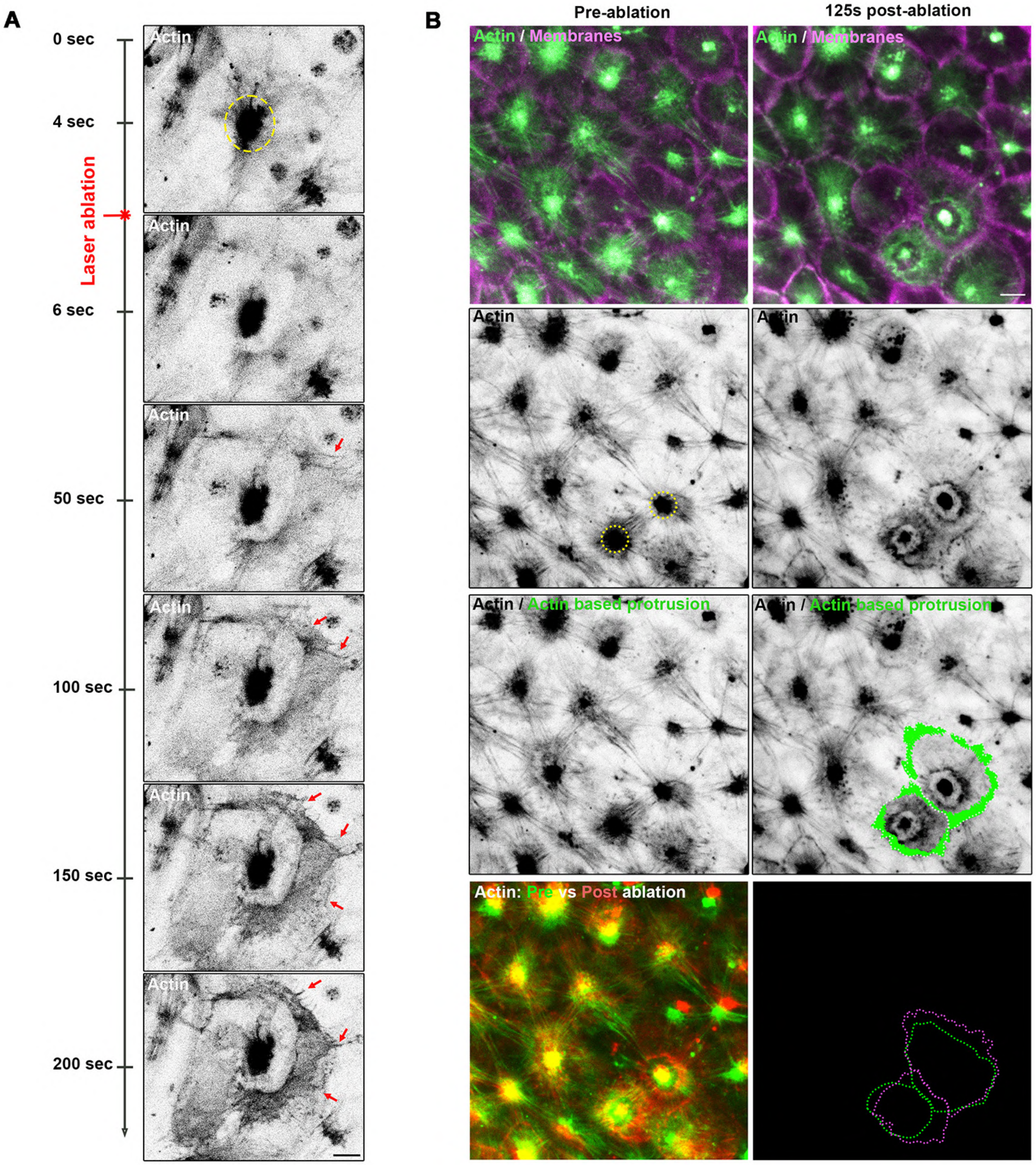


**Barai et al., Supplementary Figure 14**

**Supplementary Figure 14: (A)** Time-lapse of laser ablation of one AcS and actin protrusion formation. Red arrows point to actin-based cell protrusion. Confocal images of the basal plane show CellMaskActin staining as inverted greyscale images. Yellow circle points to the location of laser ablation. Scale bar, 5 μm. **(B)** Time-lapse of laser ablation of 2 neighbouring AcSs and protrusion formation in both cells. Actin pre-ablation (green) and post-ablation (red) are shown as overlayed image, bottom left. Actin-based protrusive structures are outlined in green. Cell contours pre- and post-ablation are outlined in green and magenta, respectively in bottom right. Yellow circles point to the location of laser ablation. Scale bar, 5 μm. Time-lapses corresponding to A and B are shown in Videos S12 and S13, respectively.

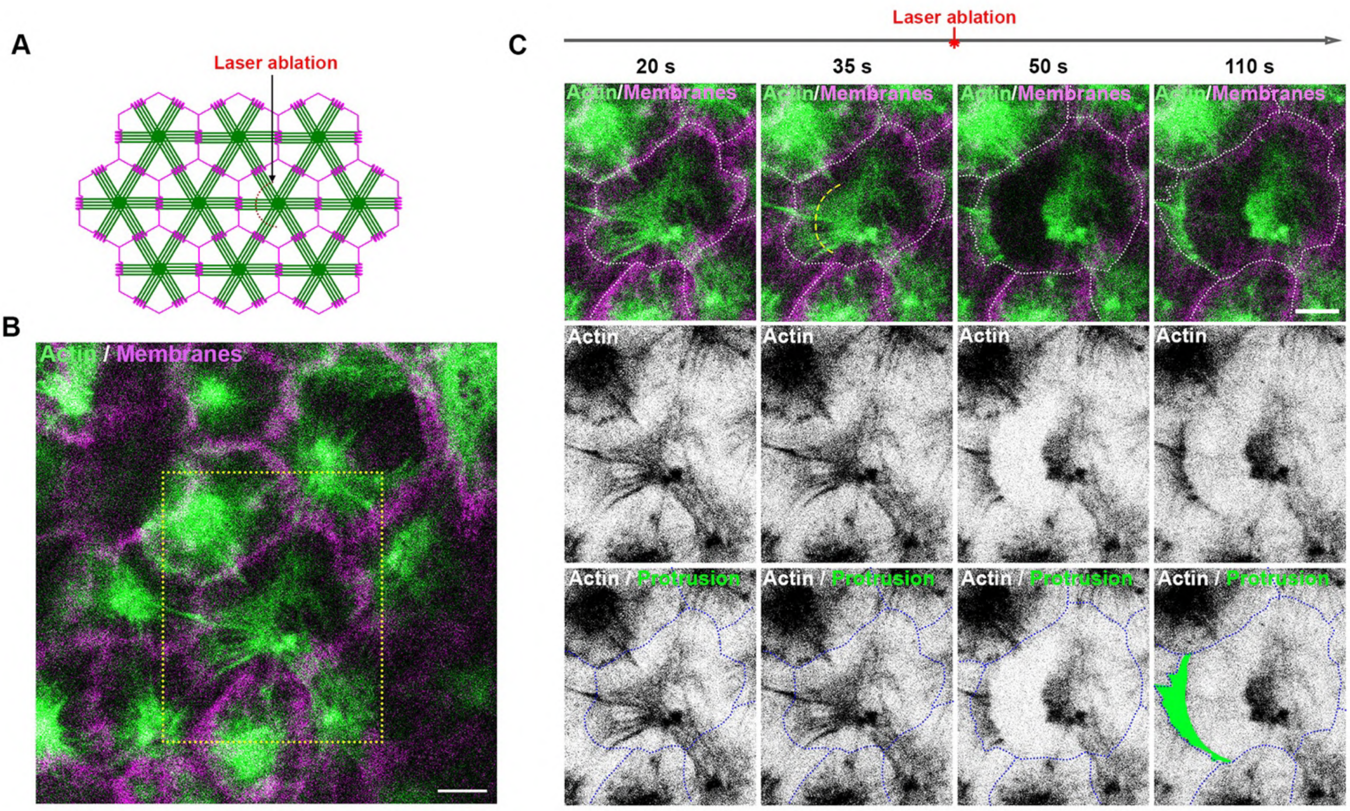


Barai et al., Supplementary Figure 15

**Supplementary Figure 15:** Time-lapse of laser ablation of half of actin branches connected to one AcS node and actin protrusion formation. **(A)** Schematics of the site of ablation during the experiment. **(B)** Confocal images of the basal plane showing insert area of interest in yellow dotted box. **(C)** Time-lapse of laser ablation of half of the actin branches connected to one AcS node. Actin-based protrusive structures are outlined in green. Segmented cell membranes are outlined with dotted white (top panel) and blue (bottom panel) lines. Yellow dotted line indicates the location of laser ablation. Scale bars, 5 µm. Corresponding time-lapse shown in Video S14.

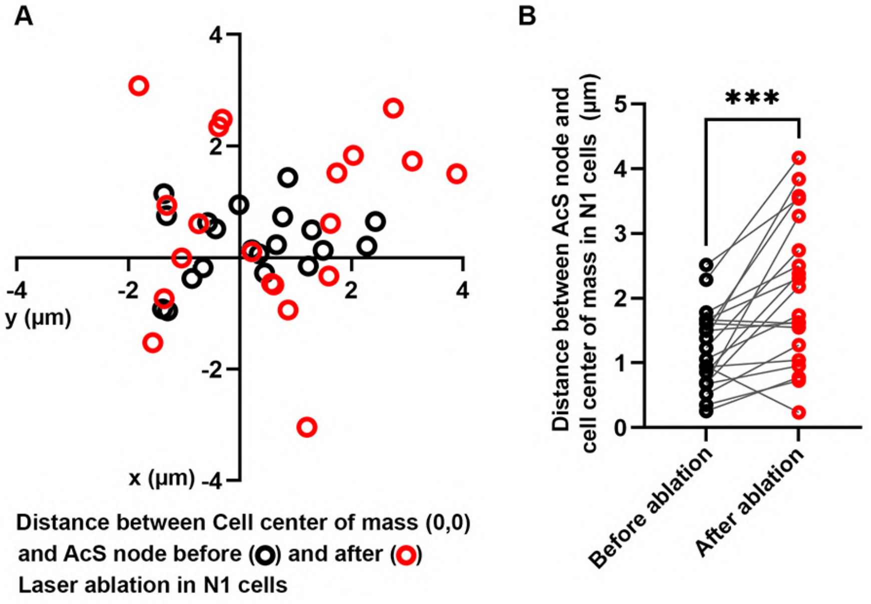


Barai et al., Supplementary Figure 16

**Supplementary Figure 16:** Quantification of the position of AcS nodes in the neighbours of laser ablated cell. **(A)** Position of the AcS nodes compared to the cell center of mass (0,0). AcS node position before laser ablation in black circle and after laser ablation in red circle. **(B)** Distance of the AcS node from cell center of mass before and after laser ablation. Distance before laser ablation 1.21 ± 0.62 µm and after ablation = 2.10 ± 1.14 µm (mean±S.D.). Paired t-test, ***p = 0.0003. N = 3 experiments, n = 20 cells.

### Supplementary Materials
### A multicellular actin star network underpins epithelial organization and connectivity

A. Barai et al.

(Dated: April 14, 2025)

In these notes, we present additional details about our numerical vertex model implementation.

#### Contents

I. Model 1: AcS formation modelled as a bulk active stress and increased junction tension 1
A. Computational model 1
B. Theoretical analysis: active-stress driven liquid-to-solid phase transition 3
C. Numerical simulations 5
1. Simulation 1: laser ablation of an actin star node 5
2. Simulation 2: blebbistatin treatment (contractility inhibition) and wash-out (contractility recovery) 5
3. Simulation 3: cell differentiation from crypt cells 6

II. Model 2: AcS formation modelled as elastic cables 7

References 18

#### I. MODEL 1: ACS FORMATION MODELLED AS A BULK ACTIVE STRESS AND INCREASED JUNCTION TENSION

##### A. Computational model

**Dynamical equation** We consider a vertex model implementation [1–3], with the following dynamics for each vertex,

$$\underbrace{\boldsymbol{F}_i^{(\text{friction})}}_{\text{Friction}} + \underbrace{\boldsymbol{F}_i^{(\text{viscous})}}_{\text{Cell viscosity}} + \underbrace{\boldsymbol{F}_i^{(\text{elastic})}}_{\text{Cell elasticity}} + \underbrace{\boldsymbol{F}_i^{(\text{AcS})}}_{\text{Actin star network contractility}} = \boldsymbol{0}, \tag{S1}$$

where we consider:

1. $\boldsymbol{F}_i^{(\text{friction})} = -\xi \boldsymbol{v}_i$ is the friction force between the monolayer and the substrate, $\xi$ being called the friction coefficient and $\boldsymbol{v}_i = \mathrm{d}\boldsymbol{r}_i/\mathrm{d}t$ being the velocity of the vertex $i$.

2. $\boldsymbol{F}_i^{(\text{viscous})}$ encompasses both dissipation at the cell–cell interfaces (viscous modulus $\eta_{ij}^{(\text{s})}$) and within the cell bulk (viscous modulus $\eta_J^{(\text{b})}$). We consider:

$$\boldsymbol{F}_i^{(\text{viscous})} = \sum_{j\in V_i} (\eta_{ij}^{(\text{s})} \boldsymbol{t}_{i,j} \cdot (\boldsymbol{v}_j - \boldsymbol{v}_i))\boldsymbol{t}_{i,j} + \sum_{J\in C_i} (\eta_J^{(\text{b})} \boldsymbol{t}_{i,J} \cdot (\boldsymbol{v}_J - \boldsymbol{v}_i))\boldsymbol{t}_{i,J}, \tag{S2}$$

where $\boldsymbol{t}_{i,\alpha} = (\boldsymbol{r}_\alpha - \boldsymbol{r}_i)/\left|\boldsymbol{r}_i - \boldsymbol{r}_\alpha\right|$ is a unit vector from the vertex $i$ to, either a neighbouring vertex ($\alpha = j$), or to the cell geometric centre ($\alpha = J$) with coordinates

$$\boldsymbol{r}_J = \frac{1}{n_J} \sum_{j\in cell} \boldsymbol{r}_j, \tag{S3}$$

where $n_J$ the number of vertices of $J$-th cell. We refer to ref. [4] for more details on the numerical implementation. In this study, we focus on the case of a uniform viscosity $\eta_{ij}^{(\text{s})} = \eta_J^{(\text{b})} = \eta$.

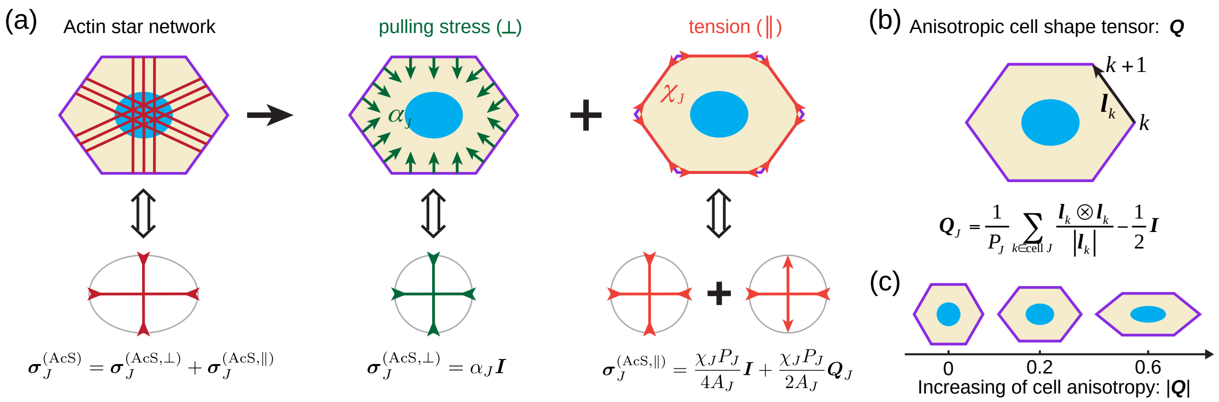


Figure S1: Mechanical characterization of the contractility of the actin star (AcS) network with a cell by an added AcS-induced pulling stress and an AcS-induced tension. (a) Mechanical description of the actin star network contractility within a cell by active stresses, $\boldsymbol{\sigma}^{(\mathrm{AcS})} = \boldsymbol{\sigma}^{(\mathrm{AcS},\perp)} + \boldsymbol{\sigma}^{(\mathrm{AcS},\parallel)}$ with $\boldsymbol{\sigma}^{(\mathrm{AcS},\perp)} = \alpha_J \boldsymbol{I}$ quantifying the pulling force normal to cell edges and $\boldsymbol{\sigma}^{(\mathrm{AcS},\parallel)} = \chi_J P_J/(4A_J)\boldsymbol{I} + \chi_J P_J/(2A_J)\boldsymbol{Q}_J$ quantifying an added active tension parallel to cell edges. (b) Sketch of the anisotropic cell shape tensor $\boldsymbol{Q}$. (c) Examples of cell shapes as a function of the magnitude of the cell anisotropic tensor $|\boldsymbol{Q}|$.

3. $\boldsymbol{F}_i^{(\mathrm{elastic})}$ is the elastic force stemming from the variation of the cell shape, which can be expressed by, $\boldsymbol{F}_i^{(\mathrm{elastic})} = -\partial E/\partial \boldsymbol{r}_i$ with the mechanical energy [2, 5–8]:

$$E = \underbrace{\sum_{J=1}^{N} \frac{1}{2} K_A (A_J - A_0)^2}_{\text{Cell area elasticity}} + \underbrace{\sum_{J=1}^{N} \frac{1}{2} K_P (P_J - P_0)^2}_{\text{Cell perimeter elasticity}}, \tag{S4}$$

where $K_A$ and $K_P$ are the rigidities associated with cell area and cell perimeter; $A_0$ and $P_0$ are the preferred cell area and the preferred cell perimeter, respectively; $A_J$ and $P_J$ are the actual area and perimeter of the $J$-th cell, respectively.

4. the AcS-induced force $\boldsymbol{F}_i^{(\mathrm{AcS})}$ corresponds to the contribution of the multicellular actin; we decompose the actin star network into two contributions, resulting from:

- (4.1) an active pulling stress $\boldsymbol{\sigma}_J^{(\mathrm{AcS},\perp)} = \alpha_J \boldsymbol{I}$ with $\alpha_J \geq 0$ quantifies the intensity of the active pulling stress induced by the contractility of the actin star network within the $J$-th cell, which describes the active pulling force normal to cell edges, see Fig. S1(a);
- (4.2) an added active tension $\chi_J > 0$ parallel to edges of the $J$-th cell, see Fig. S1(a), which results in the following $\boldsymbol{F}_k^{(\mathrm{AcS},\parallel)}$ force applied on the $k$-th vertex of the $J$-th cell:

$$\boldsymbol{F}_k^{(\mathrm{AcS},\parallel)} = \frac{1}{2}\chi_J \frac{\boldsymbol{l}_k}{|\boldsymbol{l}_k|} - \frac{1}{2}\chi_J \frac{\boldsymbol{l}_{k-1}}{|\boldsymbol{l}_{k-1}|}, \tag{S5}$$

where $\boldsymbol{l}_k = \boldsymbol{r}_{k+1} - \boldsymbol{r}_k$ the $k$-th edge vector of the $J$-th cell.

##### B. Theoretical analysis: active-stress driven liquid-to-solid phase transition

In this section, we show that the active pulling stress $\alpha_J$ is equivalent to renormalizing the preferred area of the $J$-th cell as: $A_{0,J} = A_0 - \alpha_J/K_A$; and that the AcS-induced tension $\chi_J$ is equivalent to renormalizing the preferred perimeter of the $J$-th cell as: $P_{0,J} = P_0 - \chi_J/(2K_P)$. Our derivation is based on the analytical derivation of the total stresses within cells.

We express the active stress induced by the AcS-network by (Fig. S1(a)):

$$\boldsymbol{\sigma}_J^{(\mathrm{AcS})} = \boldsymbol{\sigma}_J^{(\mathrm{AcS},\perp)} + \boldsymbol{\sigma}_J^{(\mathrm{AcS},\parallel)}, \tag{S6}$$

where

$$\boldsymbol{\sigma}_J^{(\mathrm{AcS},\perp)} = \alpha_J \boldsymbol{I}, \tag{S7}$$

quantifies the active stress induced by the AcS network (i.e., pulling forces normal to the cell edges, Fig. S1(a)), and, using the Batchelor formula [8–11],

$$\boldsymbol{\sigma}_J^{(\mathrm{AcS},\parallel)} = -\frac{1}{A_J}\sum_{k\in\mathrm{cell}\ J} \boldsymbol{r}_k \otimes \boldsymbol{F}_k^{(\mathrm{AcS},\parallel)}, \tag{S8}$$

quantifies the active stress induced by the AcS tension parallel to cell edges (Fig. S1(a)). Substituting Eq. (S5) into Eq. (S8), we obtain

$$\begin{aligned}\boldsymbol{\sigma}_J^{(\mathrm{AcS},\parallel)} &= -\frac{1}{A_J}\sum_{k\in\mathrm{cell}\ J} \boldsymbol{r}_k \otimes \left(\frac{1}{2}\chi_J\frac{\boldsymbol{l}_k}{|\boldsymbol{l}_k|} - \frac{1}{2}\chi_J\frac{\boldsymbol{l}_{k-1}}{|\boldsymbol{l}_{k-1}|}\right) = -\frac{\chi_J}{2A_J}\sum_{k\in\mathrm{cell}\ J} \boldsymbol{r}_k \otimes \left(\frac{\boldsymbol{l}_k}{|\boldsymbol{l}_k|} - \frac{\boldsymbol{l}_{k-1}}{|\boldsymbol{l}_{k-1}|}\right)\\ &= -\frac{\chi_J}{2A_J}\sum_{k\in\mathrm{cell}\ J}\left(\boldsymbol{r}_k \otimes \frac{\boldsymbol{l}_k}{|\boldsymbol{l}_k|} - \boldsymbol{r}_k \otimes \frac{\boldsymbol{l}_{k-1}}{|\boldsymbol{l}_{k-1}|}\right) = -\frac{\chi_J}{2A_J}\sum_{k\in\mathrm{cell}\ J}\left(\boldsymbol{r}_k \otimes \frac{\boldsymbol{l}_k}{|\boldsymbol{l}_k|} - \boldsymbol{r}_{k+1} \otimes \frac{\boldsymbol{l}_k}{|\boldsymbol{l}_k|}\right)\\ &= \frac{\chi_J}{2A_J}\sum_{k\in\mathrm{cell}\ J}\frac{\boldsymbol{l}_k\otimes\boldsymbol{l}_k}{|\boldsymbol{l}_k|} = \frac{\chi_J P_J}{2A_J}\left(\boldsymbol{Q}_J + \frac{1}{2}\boldsymbol{I}\right) = \frac{\chi_J P_J}{4A_J}\boldsymbol{I} + \frac{\chi_J P_J}{2A_J}\boldsymbol{Q}_J,\end{aligned} \tag{S9}$$

where $\boldsymbol{Q}_J$ is a traceless anisotropic cell shape tensor (Fig. S1(b)), defined as [8, 12]

$$\boldsymbol{Q}_J = \frac{1}{P_J}\sum_{k\in\mathrm{cell}\ J}\frac{\boldsymbol{\ell}_k\otimes\boldsymbol{\ell}_k}{|\boldsymbol{\ell}_k|} - \frac{1}{2}\boldsymbol{I}. \tag{S10}$$

For isotropic cell shape, e.g., regular hexagonal shape, $\boldsymbol{Q}_J = \boldsymbol{0}$; the magnitude $|\boldsymbol{Q}_J|$ quantifies cell anisotropy (Fig. S1(c)). Overall, the active stress induced by the AcS network can be expressed as:

$$\boldsymbol{\sigma}_J^{(\mathrm{AcS})} = \boldsymbol{\sigma}_J^{(\mathrm{AcS},\perp)} + \boldsymbol{\sigma}_J^{(\mathrm{AcS},\parallel)} = \left(\alpha_J + \frac{\chi_J P_J}{4A_J}\right)\boldsymbol{I} + \frac{\chi_J P_J}{2A_J}\boldsymbol{Q}_J. \tag{S11}$$

We recall that the elastic stress corresponding to the mechanical energy Eq. (S4) reads [8, 13]:

$$\boldsymbol{\sigma}_J^{(\mathrm{elastic})} = \left[K_A\left(A_J - A_0\right) + \frac{1}{2}\frac{K_P P_J\left(P_J - P_0\right)}{A_J}\right]\boldsymbol{I} + \frac{K_P P_J\left(P_J - P_0\right)}{A_J}\boldsymbol{Q}_J. \tag{S12}$$

Therefore, the total stress within the $J$-th cell reads:

$$\begin{aligned}\boldsymbol{\sigma}_J &= \boldsymbol{\sigma}_J^{(\mathrm{elastic})} + \boldsymbol{\sigma}_J^{(\mathrm{AcS})}\\ &= \left[K_A\left(A_J - A_0\right) + \alpha_J + \frac{1}{2}\frac{K_P P_J\left(P_J - P_0\right)}{A_J} + \frac{1}{4}\frac{\chi_J P_J}{A_J}\right]\boldsymbol{I} + \left[\frac{K_P P_J\left(P_J - P_0\right)}{A_J} + \frac{1}{2}\frac{\chi_J P_J}{A_J}\right]\boldsymbol{Q}_J\\ &= \left[K_A\left(A_J - A_{0,J}\right) + \frac{1}{2}\frac{K_P P_J\left(P_J - P_{0,J}\right)}{A_J}\right]\boldsymbol{I} + \frac{K_P P_J\left(P_J - P_{0,J}\right)}{A_J}\boldsymbol{Q}_J,\end{aligned} \tag{S13}$$

where

$$A_{0,J} = A_0 - \frac{\alpha_J}{K_A}, \tag{S14}$$

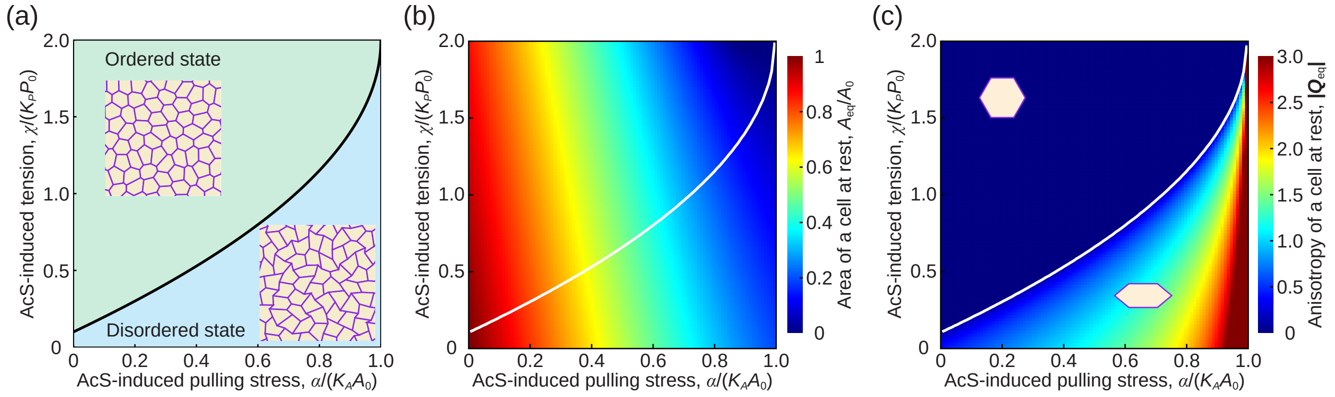


Figure S2: Theoretical analysis of cell area and cell anisotropy dictated by the AcS-induced active pulling stress $\alpha$ and the AcS-induced tension $\chi$. (a) Critical rigidity transition line Eq. (S19) (solid black line) and typical morphology of a cell monolayer within each phase. (b) The area of a hexagonal cell at rest (white line: rigidity transition line Eq. (S19)) (c) Theoretical prediction [8] of the anisotropy $|\boldsymbol{Q}|$ of a single cell at rest. (white line: rigidity transition line Eq. (S19)). Parameters: $K_A = 1$, $A_0 = 1$, $K_P = 0.02$, and $P_0 = 4$.

is a rescaled preferred area of the $J$-th cell, modulated by the isotropic active stress level $\alpha_J$, and

$$P_{0,J} = P_0 - \frac{\chi_J}{2K_P}, \tag{S15}$$

is a rescaled preferred perimeter of the $J$-th cell, modulated by the AcS-induced tension level $\chi_J$.

**Effective mechanical energy formalism.** The total stress Eq. (S13) corresponds to an effective mechanical energy of the $J$-th cell,

$$E_J^{(\mathrm{eff})} = \frac{1}{2}K_A(A_J - A_{0,J})^2 + \frac{1}{2}K_P(P_J - P_{0,J})^2, \tag{S16}$$

which satisfies

$$\boldsymbol{\sigma}_J = -\frac{1}{A_J}\sum_{k\in \mathrm{cell}\ J} \boldsymbol{r}_k \otimes \left(-\frac{\partial E_J^{(\mathrm{eff})}}{\partial \boldsymbol{r}_k}\right). \tag{S17}$$

Therefore, the contractility of the actin star network can be mimicked by a renormalized mechanical energy, $E_J^{(\mathrm{eff})}$.

**Cellular target shape index modulated by the actin star network.** Equations (S14) and (S15) show that the actin star network within cells plays a role in modulating the target shape index of the cell [6],

$$s_{0,J} = \frac{P_{0,J}}{\sqrt{A_{0,J}}} = \frac{P_0}{\sqrt{A_0}} \frac{1 - \dfrac{\chi_J}{2K_P P_0}}{\sqrt{1 - \dfrac{\alpha_J}{K_A A_0}}}. \tag{S18}$$

An increase in the AcS-induced active pulling stress $\alpha_J$ leads to an increase in the target shape index $s_{0,J}$, making cells more elongated; while an increase in the active tension $\chi_J$ leads to a decrease in the target shape index $s_{0,J}$, making cells more rounded.

Following [6], letting $s_{0,J} = s_0^\star$ leads to a critical line of rigidity transition in the $(\alpha_J\ , \chi_J)$ space:

$$\chi_J^\star = 2K_P P_0 \left(1 - s_0^\star \frac{\sqrt{A_0}}{P_0}\sqrt{1 - \frac{\alpha_J}{K_A A_0}}\right), \tag{S19}$$

where $s_0^\star$ is a critical target shape index beyond which the cell monolayer behaves as a disordered liquid; $s_0^\star \approx 3.81$ for a disordered cellular system [6]. When the AcS-induced tension is weak enough, i.e., $\chi_J < \chi_J^\star$, the cell monolayer behaves as a liquid, exhibiting a disordered and elongated cell shape pattern, see Fig. S2.

##### C. Numerical simulations

We perform numerical simulations to explore the roles of the multicellular actin star network in the context of

1. the laser ablation of an actin star node,
2. the blebbistatin treatment and wash-out experiment,
3. the progressive cell differentiation.

In all these simulations, we simulated a cell monolayer consisting of $N = 400$ cells in a square box of size $L = \sqrt{NA_0}$, using periodic boundary conditions, see Fig. S4(a). We initialize our simulations from a random Voronoi cell pattern and let the system relax toward an equilibrium state, using the procedure as described in our previous study [8]. We then (i) solve the force balance equation to obtain the motion velocity of each vertex $\{\boldsymbol{v}_i\}$; (ii) move the vertices to new positions using a forward Euler scheme,

$$\boldsymbol{r}_i(t + \Delta t) = \boldsymbol{r}_i + \boldsymbol{v}_i \Delta t, \tag{S20}$$

and (iii) perform T1 topological transitions for all short cell–cell junctions ($l_{ij} < \ell_{\mathrm{T1}} = 0.01\sqrt{A_0}$).

###### 1. *Simulation 1: laser ablation of an actin star node*

**Method** In such simulations, the activities of all cells are set to be the same before the laser ablation, $\alpha_J = \alpha^{(\mathrm{control})} > 0$ and $\chi_J = \chi^{(\mathrm{control})}$ for $J = 1, 2, 3, \cdots, N$. To model the effect of the laser ablation, we select a cell at the center of the simulation box and set its activity to be zero $\alpha_J = \alpha^{(\mathrm{ablation})} = 0$ and $\chi_J = \chi^{(\mathrm{ablation})} = 0$, see Fig. S4(a). We then let the system relax to a new equilibrium state. We track and measure the displacement of each cell during such a relaxation process. We provide the parameter values in simulations in Table I.

**Result** We find that the delay in the strain propagation scales with the distance to the ablated cell according to $\tau \sim L^2/K$, i.e. with an effective diffusion of elasticity, denoted $K$, which is inversely proportional to the friction, i.e. $K \propto 1/\xi$ (see Fig. S4(e)). In the presence of a finite friction, the evolution of the strain is only marginally dependent on the cell-cell viscosity value, see Fig. S4(d). In the limit of vanishing friction ($\xi \to 0$), the strain is propagated at large distances ($N = 8$) in less than a second (blue curve in Fig. S4(e)).

**Discussion** Diffusion of elasticity is also observed in an optical tweezer mechanical perturbation of cell membranes in the Drosophila embryo [14]. There, the time delay in the strain is measured at $\tau_1 = 150 \pm 85\,\mathrm{ms}$ at a one-cell distance ($\sim 7\mu$m) to the mechanical perturbation site. Such value lies within the time resolution of our experiments. Based on the diffusion of elasticity scaling, we expect that the corresponding value at the six-cell distance should scale as $\tau_6 = 36\tau_1$, corresponding to $\tau_6 \approx 5 \pm 0.5\,\mathrm{s}$. Such an estimate is lower than the six-cell distance measured in the experiments, around $\tau_6 \approx 40$s, see Fig. 6. Overall, the observation that the strain propagation is less rapid than in the Drosophila embryo is compatible with a higher friction to the substrate. This is expected since, at this stage, there is limited adhesion of Drosophila cells to the vitelline membrane.

###### 2. *Simulation 2: blebbistatin treatment (contractility inhibition) and wash-out (contractility recovery)*

**Method** Here, the activities of all cells are set to be the same before the blebbistatin treatment, $\alpha_J = \alpha^{(\mathrm{control})} > 0$ and $\chi_J = \chi^{(\mathrm{control})} > 0$ for $J = 1, 2, 3, \cdots, N$. To model the effect of the blebbistatin treatment, we select a group of cells with a number $N_{\mathrm{blebbistatin}} = 50$ at the center region of the simulation box and set their activities to zero $\alpha_J = \alpha^{(\mathrm{blebbistatin})} = 0 < \alpha^{(\mathrm{control})}$ and $\chi_J = \chi^{(\mathrm{blebbistatin})} = 0 < \chi^{(\mathrm{control})}$ for cells being treated with blebbistatin, see Fig. S5(a). We then let the system relax to a new equilibrium state. Subsequently, to model the effect of blebbistatin wash-out, we recover the activity level of these cells being treated with blebbistatin. We provide the parameter values in simulations in Table II.

**Result** With our set of parameter values, the cell area and cell anisotropy are both increased in the blebbistatin-treated cells; the cell area increase is chosen to match the values observed in experiments. In the blebbistatin wash-out experiment, the recovery of the actin star network leads to a progressive return of the cell area to the pre-blebbistatin treatment value (Fig. 5D). This experiment is important in setting the values of $\alpha$ and $\chi$ associated with the onset of the actin star network.

##### 3. *Simulation 3: cell differentiation from crypt cells*

**Method** Here, to mimic the process of cell differentiation, the activities of all cells are set to be the same before differentiation, $\alpha_J = \alpha^{(\text{undifferentiated})}$ and $\chi_J = \chi^{(\text{undifferentiated})}$ for $J = 1, 2, 3, \cdots, N$.

We keep constant the $(\alpha_J, \chi_J)$ parameters within a group of cells with a number $N_{\text{differentiated}} = 50$ at the center region of the simulation box - to mimic the onset of crypt-like cells domains - while switching the $(\alpha_J, \chi_J)$ parameters to $\alpha_J = \alpha^{(\text{differentiated})}$ and $\chi_J = \chi^{(\text{differentiated})}$ within the rest of the tissue, mimicking differentiation into villus-like cells, see Fig. S6(a).

We then let the system relax to a new equilibrium state. We provide the parameter values in simulations in Table III and IV.

**Result** We test two possible cell differentiation paths in the $(\alpha, \chi)$ space, see Fig. S6(b,c).

- in the cell differentiation path I differentiation path, the cells within the interior domain increase their area, akin to cells subjected to the blebbistanin treatment experiments.
- in the cell differentiation path II, the cells within the interior domain display a lower area than differentiated ones, as crypt-like cells. We also observe a progressive increase of the triangular order within the differentiated cells; see Fig. S6(g). As in experiments, the hexatic order remained relatively unchanged throughout the cell differentiation process, while the triangular order evolved significantly; see Fig. S6(f-g).

#### II. MODEL 2: ACS FORMATION MODELLED AS ELASTIC CABLES

So far, we have considered a generic model in which the AcS active pulling stress $\alpha_J$ and AcS-induced tension $\chi_J$ are two independent parameters.

Here, we propose to focus on a more specific microscopic model which encompasses the observation that the actin star seemingly connects each of the junction mid-points to the cell center, see Fig. S3. We express the relation of this new model to the former one described above in terms of a linear relation between the actin star tension $\xi$ (new model) with the active stress $\alpha_J$ and AcS-induced tension $\chi_J$ (former model).

**New model description** Here, we propose to model the actin star network through a set of additional forces, denoted $\boldsymbol{T}_i$, each applied to the junction mid-points and pointing toward the cell center, $\boldsymbol{r}_C = \sum_{i \in cell} \boldsymbol{r}_i/n$, with $n$ the number of edges of the cell. Based on the Batchelor formula [8–11], the coarse-grained active stress reads

$$\boldsymbol{\sigma}^{(\mathrm{AcS})} = -\frac{1}{A} \sum_{i \in cell} \boldsymbol{s}_i \otimes \boldsymbol{T}_i. \tag{S21}$$

where $\boldsymbol{s}_i = \boldsymbol{\rho}_i + \boldsymbol{l}_i/2 = \boldsymbol{r}_i - \boldsymbol{r}_C + (\boldsymbol{r}_{i+1} - \boldsymbol{r}_i)/2 = (\boldsymbol{r}_i + \boldsymbol{r}_{i+1})/2 - \boldsymbol{r}_C$ is the length vector $\boldsymbol{s}_i$ of the $i$-th actin cable, assumed to connect the cell center to the junction mid-point.

Here, we focus on the case of a linear relationship between the active force $\boldsymbol{T}_i$ and the $i$-th actin cable vector $\boldsymbol{s}_i$

$$\boldsymbol{T}_i = -\zeta \boldsymbol{s}_i, \tag{S22}$$

where $\zeta > 0$ quantifies the contractility of the actin star network. Substituting Eq. (S22) into Eq. (S21), the active stress then takes the expression:

$$\boldsymbol{\sigma}^{(\mathrm{AcS})} = \zeta \boldsymbol{W}, \tag{S23}$$

where $W$ is what we call the mid-point cell shape tensor (Fig. S3), defined as:

$$\boldsymbol{W} = \frac{1}{A} \sum_{i \in \mathrm{cell}} \boldsymbol{s}_i \otimes \boldsymbol{s}_i. \tag{S24}$$

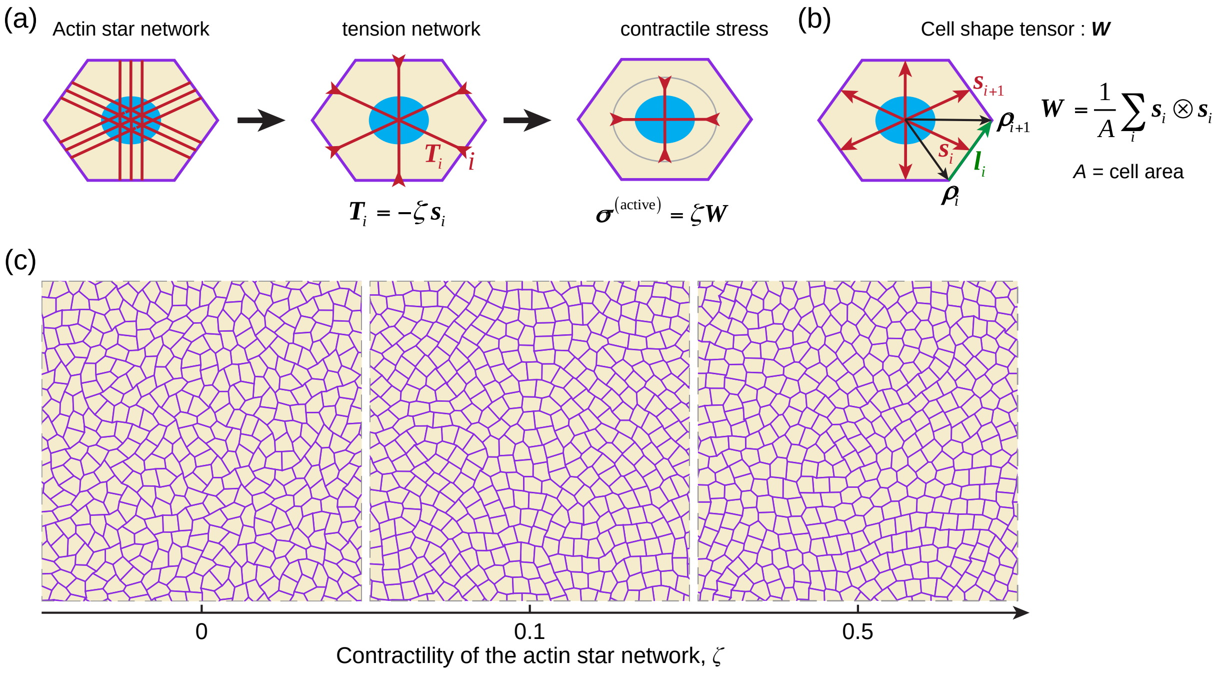


Figure S3: Alternative stress description of the actin star (AcS) network. (a, b) Model sketches of the AcS network, active stresses, and cell shape tensor. (a) Mechanical description of the actin star network contractility within a cell by active stresses, $\boldsymbol{\sigma}^{(\mathrm{AcS})} = \zeta \boldsymbol{W}$ with $\zeta > 0$ quantifying the contractility level and $\boldsymbol{W}$ a cell shape tensor. (b) Sketch of the cell shape tensor $\boldsymbol{W}$. (c) Numerical simulations using the alternative stress description (Eq. (S23)). An increase in the contractility parameter $\zeta$ leads to more rounded cell shapes. Parameters: $K_A = 1$, $A_0 = 1$, $K_P = 0.02$, and $P_0 = 4$.

In particular, for a regular hexagonal cell shape, $\boldsymbol{W} = (\sqrt{3}/2)\boldsymbol{I}$.

Let us here consider a regular hexagonal cell undergoing affine transformation, $(x, y) \to (\lambda_1 x, \lambda_2 y)$, with $\lambda_1$ and $\lambda_2$ stretches along the $x$-axis and the $y$-axis. In this case, the mid-point cell shape tensor $\boldsymbol{W}$ reads

$$\boldsymbol{W} = \frac{\sqrt{3}}{2}\frac{\lambda_1}{\lambda_2}\hat{\boldsymbol{x}} \otimes \hat{\boldsymbol{x}} + \frac{\sqrt{3}}{2}\frac{\lambda_2}{\lambda_1}\hat{\boldsymbol{y}} \otimes \hat{\boldsymbol{y}}, \tag{S25}$$

while the $\boldsymbol{Q}$ tensor defined in Eq. (S10) reads:

$$\boldsymbol{Q} = \left(\frac{\lambda_1}{\sqrt{\lambda_1^2 + 3\lambda_2^2}} - \frac{1}{2}\right)\hat{\boldsymbol{x}} \otimes \hat{\boldsymbol{x}} - \left(\frac{\lambda_1}{\sqrt{\lambda_1^2 + 3\lambda_2^2}} - \frac{1}{2}\right)\hat{\boldsymbol{y}} \otimes \hat{\boldsymbol{y}}. \tag{S26}$$

For small pure shear deformations, $\lambda_1 = 1 + \varepsilon$ and $\lambda_2 = 1 - \varepsilon$ with $\varepsilon \ll 1$, we find that

$$\boldsymbol{W} \approx \frac{\sqrt{3}}{2}\boldsymbol{I} + \sqrt{3}\varepsilon(\hat{\boldsymbol{x}} \otimes \hat{\boldsymbol{x}} - \hat{\boldsymbol{y}} \otimes \hat{\boldsymbol{y}}), \tag{S27}$$

while

$$\boldsymbol{Q} \approx \frac{3}{4}\varepsilon(\hat{\boldsymbol{x}} \otimes \hat{\boldsymbol{x}} - \hat{\boldsymbol{y}} \otimes \hat{\boldsymbol{y}}). \tag{S28}$$

Thus, we find that, at first order in the pure shear amplitude,

$$\boldsymbol{\sigma}^{(\mathrm{AcS})} \approx \frac{\sqrt{3}\zeta}{2}\boldsymbol{I} + \frac{4\sqrt{3}\zeta}{3}\boldsymbol{Q}. \tag{S29}$$

Comparing it to Eq. (S11) and taking $A_J = A_0$ and $P_J = P_0$, we get relationships between the contractility activity parameter $\zeta$, $\alpha_J$ and $\chi_J$:

$$\begin{cases} \alpha_J \approx -\dfrac{\sqrt{3}}{6}\zeta \\ \chi_J \approx \dfrac{8\sqrt{3}}{3}\dfrac{A_0}{P_0}\zeta \end{cases} \tag{S30}$$

We, therefore, expect that the forces defined in Eq. (S22) can be recast into the formalism of Eq. (S11): an increase in the contractility parameter $\zeta$ then maps into a decrease in $\alpha$ and an increase in $\chi$; a sufficiently large increase in $\zeta$ thus can trigger a transition from a liquid to a solid regime, see Fig. S2(a) and Fig. S3(c).

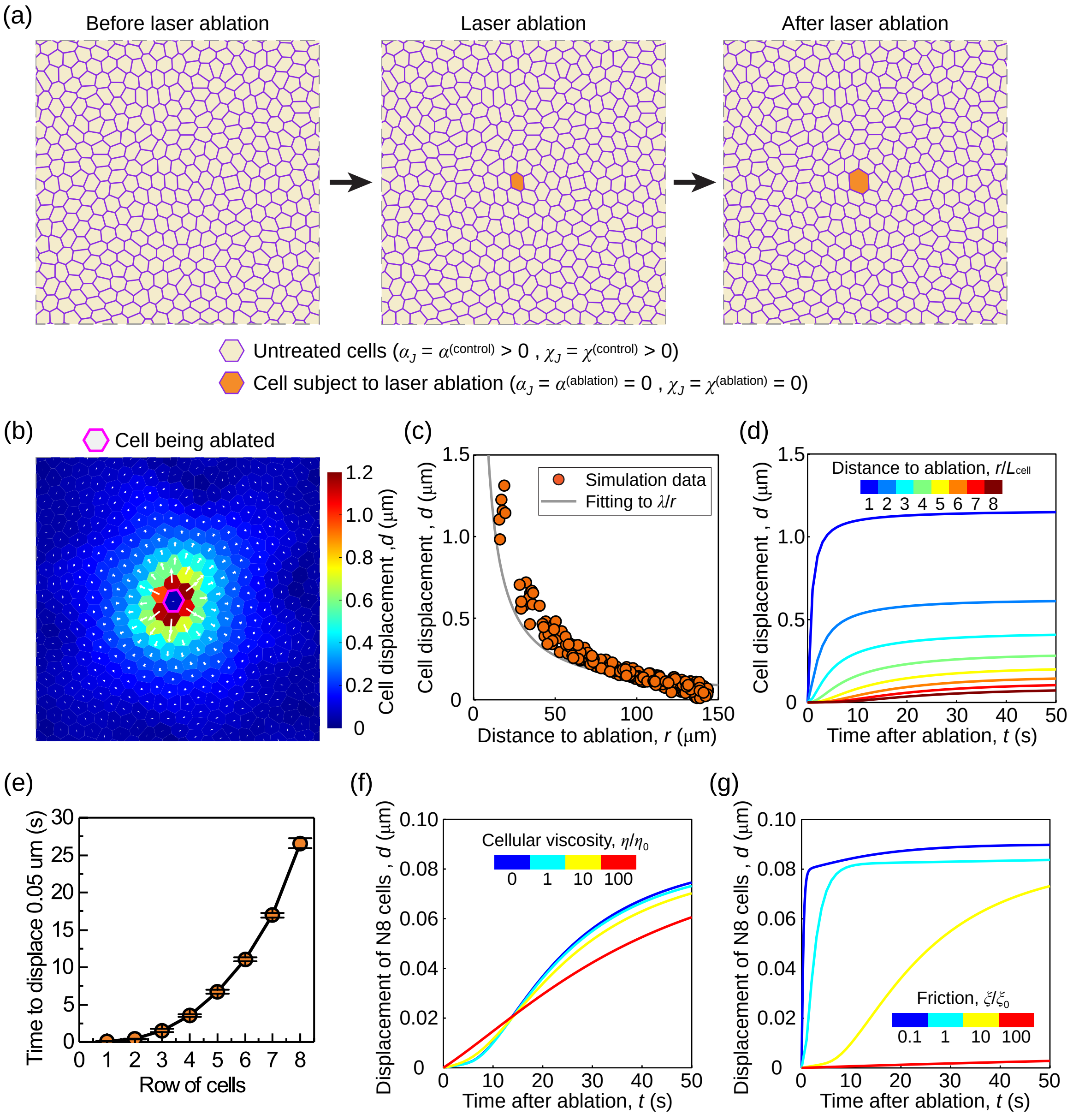


Figure S4: **Simulation 1: the laser ablation of an actin star node.** (a) Sketch of the active vertex model to simulate the laser ablation response of an intestinal epithelial monolayer. Vertex model description of an intestinal epithelial cell monolayer by a two-dimensional tiling of polygons. To mimic the effect of laser ablation, we set the activity of the cell being ablated to zero (i.e., $\alpha_J = 0$ and $\chi_J = 0$). (b-f) Simulation results of the laser ablation response of an intestinal epithelial monolayer. (b) The displacement of cells at time $t = 50$ s in response to the laser ablation (applied at $t = 0$). The color code represents the magnitude of cell displacement, while the white arrows refer to the cell displacement vector. The cell being ablated is marked by the magenta contour. (c) Scatter plot of the cell displacement magnitude as a function of the distance of the cell center to the ablation site. Symbols: simulation data. Gray solid line: fitting to $\lambda/r$ with $\lambda = 13.5\ \mu\text{m}^2$ fitted by the least squares method. (d) The average displacement magnitude of cells at the $k$-th row as a function of time $t$ after ablation. The cells at the $k$-th row are defined by the distance $d$ of the cell center to the ablation site if it satisfies $k - 1/2 < d/L_{\text{cell}} < k + 1/2$ with $L_{\text{cell}} = 10\ \mu$m being the cell size. (e) Time to displace 0.05 $\mu$m, $t_{0.05\ \mu\text{m}}$, as a function of the row of cells (i.e., distance to the laser ablation site). Averaged over $n = 5$ independent simulations. Data = mean $\pm$ SD. (f) The average displacement magnitude of cells at the 8-th row (denoted N8 cells) as a function of time $t$ after ablation, for different levels of cellular viscosities expressed in the unit of $\eta_0 = 0.01\ \text{nN} \cdot \text{s} \cdot \mu\text{m}^{-1}$, with fixed friction $\xi = 0.1\ \text{nN} \cdot \text{s} \cdot \mu\text{m}^{-1} = 10\eta_0$; (g) The average displacement magnitude of cells at the 8-th row (denoted N8 cells) as a function of time $t$ after ablation, for different levels of friction expressed in units of $\xi_0 = 0.01\ \text{nN} \cdot \text{s} \cdot \mu\text{m}^{-1}$, with fixed cellular viscosity $\eta = 0.01\ \text{nN} \cdot \text{s} \cdot \mu\text{m}^{-1} = \xi_0$. Other parameters are set in Table I.

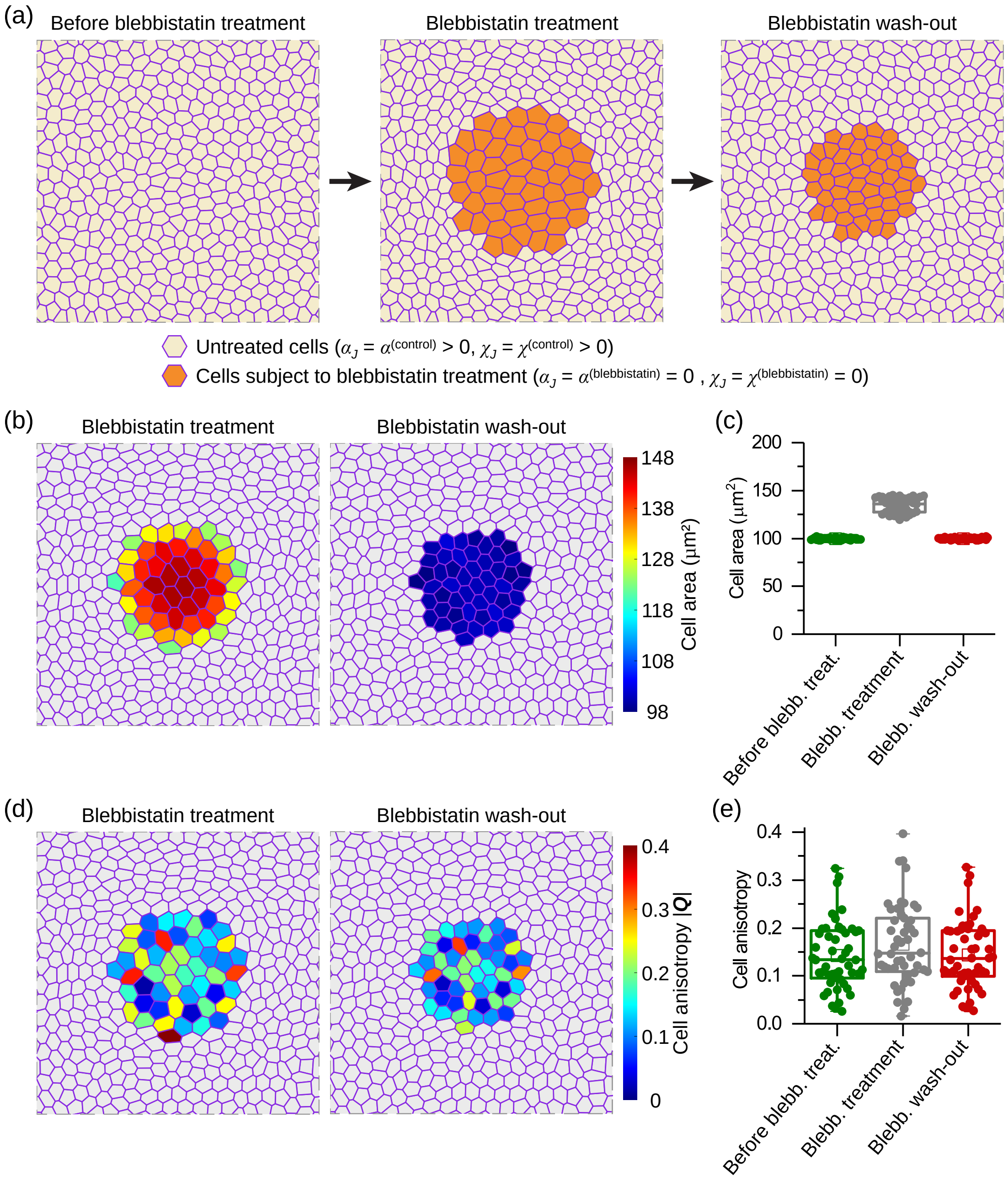


Figure S5: **Simulation 2: contractility recovery by blebbistatin wash-out.** (a) Sketch of the active vertex model to simulate the effect of blebbistatin treatment and the subsequent contractility recovery process by blebbistatin wash-out. The blebbistatin treatment is mimicked by a decrease in the contractility activity of the actin star (AcS) network within the cells being treated. The blebbistatin wash-out effect is mimicked by recovering the contractility activity in the cells being treated. (b-e) Simulation results. (b) The cell area field after blebbistatin treatment (*left*) and the cell area field after blebbistatin wash-out (*right*). Here, we show the area of cells being treated. (c) Statistical analysis of cell area before blebbistatin treatment, after blebbistatin treatment, and after blebbistatin wash-out. The cell area are $99.8 \pm 1.0$ $\mu$m$^2$ (mean $\pm$ S.D.), $134.1 \pm 7.7$ $\mu$m$^2$ (mean $\pm$ S.D.) and $99.9 \pm 1.0$ $\mu$m$^2$, in turn. $n = 50$ cells. (d) The cell anisotropy field after blebbistatin treatment (*left*) and the cell area field after blebbistatin wash-out (*right*). Here, we show the anisotropy of cells being treated. (e) Statistical analysis of cell anisotropy before blebbistatin treatment, after blebbistatin treatment, and after blebbistatin wash-out. The cell anisotropy are $0.142 \pm 0.070$ $\mu$m$^2$ (mean $\pm$ S.D.), $0.165 \pm 0.084$ $\mu$m$^2$ (mean $\pm$ S.D.) and $0.143 \pm 0.070$ $\mu$m$^2$, in turn. $n = 50$ cells. Simulation parameters are provided in Table II.

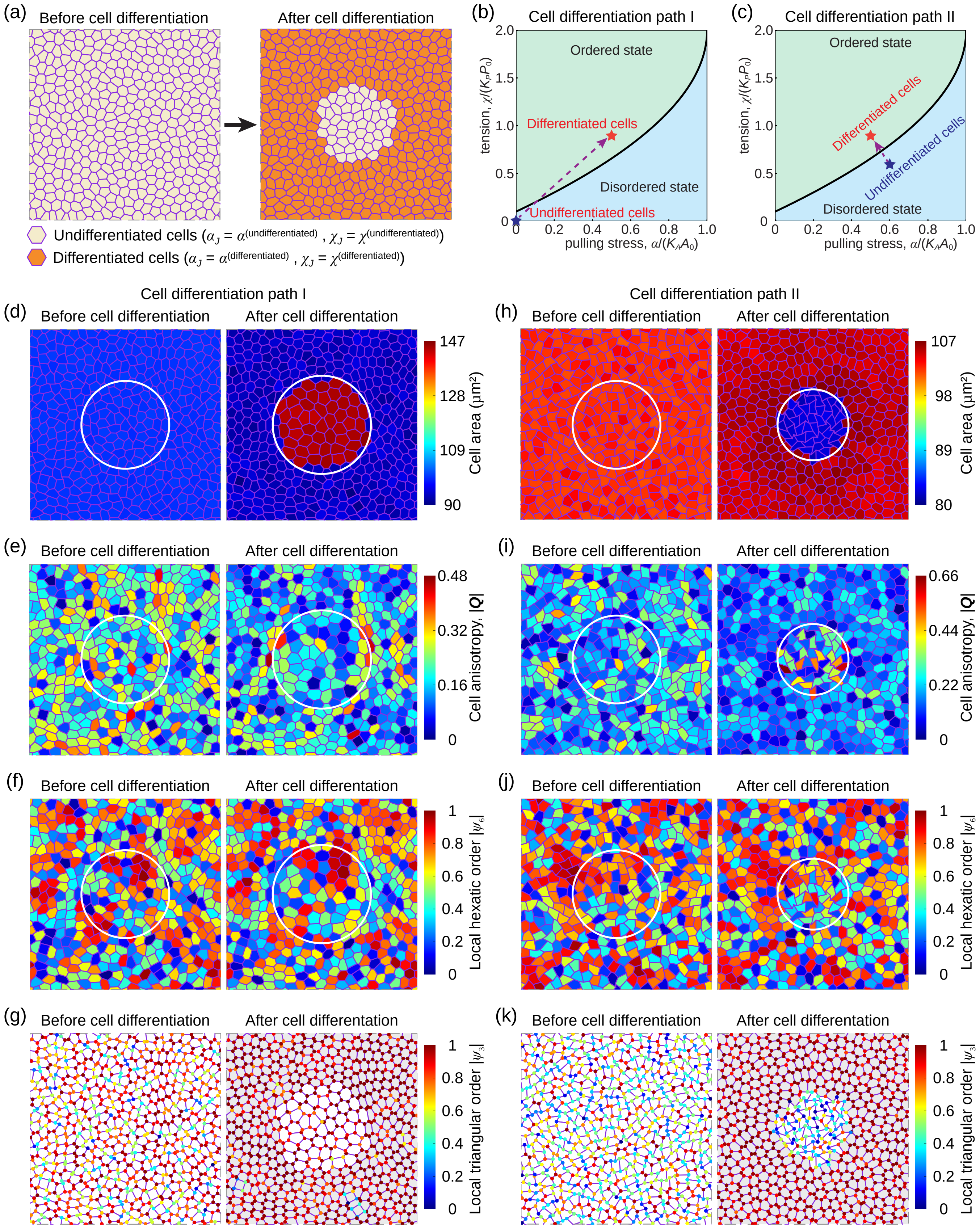


Figure S6: (Caption on the next page.)

Figure S6: (Previous page.) **Simulation 3: cell differentiation from crypt-like cells to villus-like cells.** (a) Sketch of the active vertex model to simulate the process of cell differentiation from crypt-like cells (light yellow) to villus-like cells (orange). Initially, cells are all the same. We mimic the cell differentiation effect by selecting a group of cells and setting up a different level of activity. (b, c) Two possible cell differentiation paths in the ($\alpha$ , $\chi$) space. (d-k) Simulation results of (d, h) the cell area field, (e, i) the anisotropy field, (f, j) the local hexatic order parameter field, and (g, k) the local triangular order parameter field (*left*) before and (*right*) after cell differentiation. Here, the cell anisotropy is quantified by the magnitude of the anisotropic cell shape tensor $|\boldsymbol{Q}|$. In (g, k), differentiated cells are marked as grey. Results (d-g) correspond to the path shown in (b); while results (h-k) correspond to the path shown in (c). Parameter values used in such simulations are given in Table III and IV.

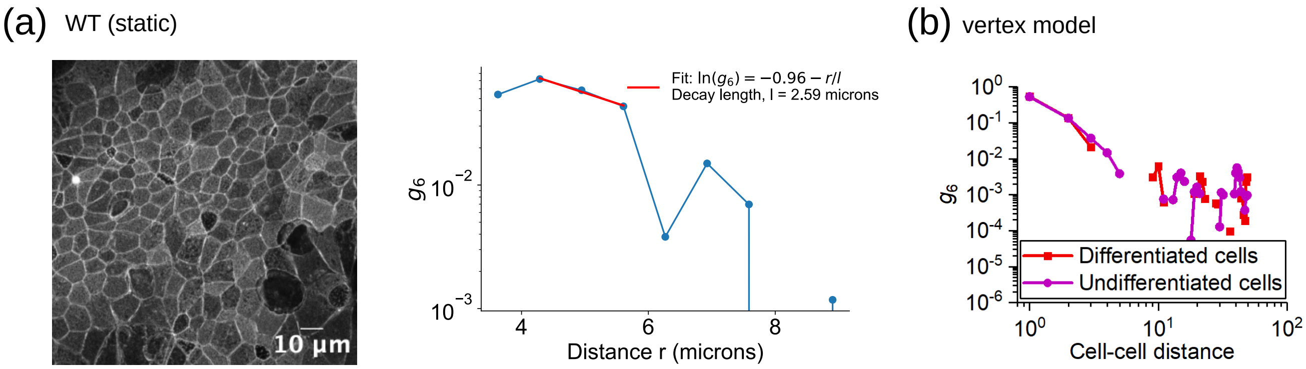


Figure S7: Spatial decay of the normalized spatial correlation function $g_6(r = r_i - r_j) = \langle \Psi_i \Psi_j^\star \rangle / \langle \Psi_i \Psi_i^\star \rangle$, where $\Psi = \psi_6$ is the local hexatic order defined in the main text, Method section, in: (a) a WT experiment, with exponential fit with a $l = 2.5\,\mu$m decay length, which corresponds to 1 cell radius, approximatively; (b) vertex model, in the differentiated (red) and undifferentiated (magenta) case. The decay appears to be faster than algebraic with a $-1/4$ exponent, indicating that there is no long-range order. Default parameter sets.

TABLE I: List of default parameter values used in our simulation 1 of laser cell ablation, Sec. I C 1.

| Parameter | Description | Value |
|---|---|---|
| $\ell = \sqrt{A_0}$ | Length scale | 10 $\mu$m |
| $\tau$ | Time scale | 0.1 s |
| $f = K_A A_0^{3/2}$ | Force scale | 1 nN |
| $K_A$ | Cell area stiffness | $10^6$ N $\cdot$ m$^{-3}$ |
| $A_0$ | Preferred cell area | 100 $\mu$m$^2$ |
| $K_P$ | Cell perimeter stiffness | 0.002 nN $\cdot$ $\mu$m$^{-1}$ |
| $P_0$ | Preferred cell perimeter | 40 $\mu$m |
| $\xi$ | Cell-substrate friction | 0.1 nN $\cdot$ s $\cdot$ $\mu$m$^{-1}$ |
| $\eta$ | Viscosity of cells | 0.01 nN $\cdot$ s $\cdot$ $\mu$m$^{-1}$ |
| $\alpha^{(\text{control})}$ | AcS-induced pulling stress level of control cells | 0.05 nN $\cdot$ $\mu$m$^{-1}$ |
| $\alpha^{(\text{ablation})}$ | AcS-induced pulling stress level of the ablated cell | 0 |
| $\chi^{(\text{control})}$ | AcS-induced tension level of control cells | 0.064 nN |
| $\chi^{(\text{ablation})}$ | AcS-induced tension level of the ablated cell | 0 |
| $\Delta t$ | Simulation time step | 0.001 s |

TABLE II: List of default parameter values used in simulation 2, contractility recovery, Sec. I C 2.

| Parameter | Description | Value |
|---|---|---|
| $\ell = \sqrt{A_0}$ | Length scale | 10 $\mu$m |
| $\tau$ | Time scale | 0.1 s |
| $f = K_A A_0^{3/2}$ | Force scale | 1 nN |
| $K_A$ | Cell area stiffness | $10^6$ N $\cdot$ m$^{-3}$ |
| $A_0$ | Preferred cell area | 100 $\mu$m$^2$ |
| $K_P$ | Cell perimeter stiffness | 0.002 nN $\cdot$ $\mu$m$^{-1}$ |
| $P_0$ | Preferred cell perimeter | 40 $\mu$m |
| $\xi$ | Cell-substrate friction | 0.1 nN $\cdot$ s $\cdot$ $\mu$m$^{-1}$ |
| $\eta$ | Viscosity of cells | 0.01 nN $\cdot$ s $\cdot$ $\mu$m$^{-1}$ |
| $\alpha^{(\text{blebbistatin})}$ | AcS-induced pulling stress level of cells after blebbistatin treatment | 0 |
| $\alpha^{(\text{control})}$ | AcS-induced pulling stress level of cells before blebbistatin treatment or after blebbistatin wash-out | 0.05 nN $\cdot$ $\mu$m$^{-1}$ |
| $\chi^{(\text{blebbistatin})}$ | AcS-induced tension level of cells after blebbistatin treatment | 0 |
| $\chi^{(\text{control})}$ | AcS-induced tension level of cells before blebbistatin treatment or after blebbistatin wash-out | 0.064 nN |
| $\Delta t$ | Simulation time step | 0.001 s |

TABLE III: List of default parameter values used in simulation 3, cell differentiation (path I), Sec. I C 3.

| Parameter | Description | Value |
|---|---|---|
| $\ell = \sqrt{A_0}$ | Length scale | 10 $\mu$m |
| $\tau$ | Time scale | 0.1 s |
| $f = K_A A_0^{3/2}$ | Force scale | 1 nN |
| $K_A$ | Cell area stiffness | $10^6$ N $\cdot$ m$^{-3}$ |
| $A_0$ | Preferred cell area | 100 $\mu$m$^2$ |
| $K_P$ | Cell perimeter stiffness | 0.002 nN $\cdot$ $\mu$m$^{-1}$ |
| $P_0$ | Preferred cell perimeter | 40 $\mu$m |
| $\xi$ | Cell-substrate friction | 0.1 nN $\cdot$ s $\cdot$ $\mu$m$^{-1}$ |
| $\eta$ | Viscosity of cells | 0.01 nN $\cdot$ s $\cdot$ $\mu$m$^{-1}$ |
| $\alpha^{(\text{undifferentiated})}$ | AcS-induced pulling stress level of undifferentiated cells (crypt-like cells) | 0 |
| $\alpha^{(\text{differentiated})}$ | AcS-induced pulling stress level of differentiated cells (villus-like cells) | 0.05 nN $\cdot$ $\mu$m$^{-1}$ |
| $\chi^{(\text{undifferentiated})}$ | AcS-induced tension level of undifferentiated cells (crypt-like cells) | 0 |
| $\chi^{(\text{differentiated})}$ | AcS-induced tension level of differentiated cells (villus-like cells) | 0.064 nN |
| $\Delta t$ | Simulation time step | 0.001 s |

TABLE IV: List of default parameter values used in simulation 3, cell differentiation (path II), Sec. I C 3.

| Parameter | Description | Value |
|---|---|---|
| $\ell = \sqrt{A_0}$ | Length scale | 10 $\mu$m |
| $\tau$ | Time scale | 0.1 s |
| $f = K_A A_0^{3/2}$ | Force scale | 1 nN |
| $K_A$ | Cell area stiffness | $10^6$ N $\cdot$ m$^{-3}$ |
| $A_0$ | Preferred cell area | 100 $\mu$m$^2$ |
| $K_P$ | Cell perimeter stiffness | 0.002 nN $\cdot$ $\mu$m$^{-1}$ |
| $P_0$ | Preferred cell perimeter | 40 $\mu$m |
| $\xi$ | Cell-substrate friction | 0.1 nN $\cdot$ s $\cdot$ $\mu$m$^{-1}$ |
| $\eta$ | Viscosity of cells | 0.01 nN $\cdot$ s $\cdot$ $\mu$m$^{-1}$ |
| $\alpha^{(\text{undifferentiated})}$ | AcS-induced pulling stress level of undifferentiated cells (crypt-like cells) | 0.06 nN $\cdot$ $\mu$m$^{-1}$ |
| $\alpha^{(\text{differentiated})}$ | AcS-induced pulling stress level of differentiated cells (villus-like cells) | 0.05 nN $\cdot$ $\mu$m$^{-1}$ |
| $\chi^{(\text{undifferentiated})}$ | AcS-induced tension level of undifferentiated cells (crypt-like cells) | 0.056 nN |
| $\chi^{(\text{differentiated})}$ | AcS-induced tension level of differentiated cells (villus-like cells) | 0.064 nN |
| $\Delta t$ | Simulation time step | 0.001 s |
